# Benchmarking niche identification via domain segmentation for spatial transcriptomics data

**DOI:** 10.64898/2026.02.27.708202

**Authors:** Yuxuan Wang, Yuhan Chen, Luqi Yang, Cheng Wang, Jinpu Cai, Hongyi Xin

## Abstract

Tissue niches are spatially organized microenvironments in which coordinated multicellular interactions shape cellular states and biological functions. Currently, niche identification is routinely performed using domain segmentation frameworks. While interrelated, spatial domains and niches are not fundamentally equivalent. The former emphasizes intra-domain compositional consistency and transcriptomic homogeneity, whereas the latter is defined by the emergent properties of localized signaling gradients and the functional reciprocity between key cell lineages. Here, we present a high-resolution reference by thoroughly annotating single-cell resolution CosMx ST data of a human follicular lymphoid hyperplasia lymph node, a dynamic, non-compartmentalized tissue containing several critical immune niches defined by specific lineage architectures. We systematically benchmarked 16 contemporary domain segmentation algorithms, demonstrating that most methods in their default configurations fail to recapitulate biologically defined niche boundaries. Our analysis reveals that the definitive, disjoint spatial distributions of key functional lineages are frequently obscured by the stochastic infiltration of peripheral cell types. Such reduction in the spatial signal-to-noise ratio represents a primary bottleneck for existing algorithms, which prioritize local transcriptomic variance over global architectural logic. Following this observation, we demonstrate that strategic weighting of core functional lineages can restore the resolution of spatial niches in select domain segmentation frameworks. Cross-comparison against compartmentalized tissues further underscores the unique challenges of niche identification in non-mechanically separated environments and clarifies the fundamental divergence between structural domain segmentation and functional niche discovery. Our work delineates the limitations of current paradigms and advocates for the development of specialized computational approaches tailored specifically to the complexity of functional microenvironments.

## 1 Introduction

Tissue niches are spatially organized, local microenvironments within tissues that maintain and regulate cellular behavior through coordinated interactions between multiple cell types [1, 2]. These microenvironments consist of diverse sets of cellular components which collectively influence cell fate decisions and tissue function. These niches are fundamentally important in developmental biology, adult tissue homeostasis, regeneration, and disease progression, as they determine whether stem and progenitor cells undergo self-renewal or differentiation, and they establish the phenotypic identity and functional properties of mature cell populations [2]. In immune organs such as lymph nodes, tonsils, and spleen, immune cell phenotype, function, and activation state are highly dynamic and constantly remodeled by neighboring cells within the local microenvironment [3, 4]. The plasticity of immune cells creates a bidirectional communication loop wherein the microenvironment shapes immune function while immune cells simultaneously remodel their local niche. In tissue regeneration, the regenerative niche serves as a temporally dynamic microenvironment that coordinates stem and progenitor cell activation, proliferation, and lineage specification to restore tissue structure and function following injury. During embryonic development, spatially organized microenvironments coordinate cell fate specification and tissue patterning across multiple scales, from early blastocyst symmetry-breaking to organogenesis, through well-characterized morphogen gradients including Wnt, TGF-*β*/Nodal, and FGF signaling emanating from localized signaling centers [5].

The advent of spatial transcriptomics (ST) has revolutionized the in situ interrogation of tissue niches by preserving the physical coordinates of gene expression [6]. Current ST technologies span a broad resolution continuum. Spot-based capturing (e.g., 10x Visium) profiles multicellular mixtures, offering robust tissue-wide coverage but requiring deconvolution to infer niche composition. Conversely, single-cell and subcellular platforms, encompassing targeted multiplexed FISH-based methods (e.g., MERFISH [7], Xenium, CosMx) and highdensity sequencing arrays (e.g., Stereo-seq), provide granular maps of precise cell-to-cell interactions, yet introduce analytical challenges like signal sparsity and localized transcriptional noise. While these diverse modalities have rapidly accelerated the discovery of spatially restricted immune hubs and progenitor zones, extracting functionally cohesive, biologically meaningful niches from such complex, multi-resolution data remains a critical computational challenge.

Biologically, tissue niches can be defined as local pockets of cells with characteristic compositions that collectively perform specific biological functions. Within each niche, particular cell types are enriched and arranged in organized spatial configurations that support specialized roles, such as stem cell maintenance, immune regulation, or tissue repair. The same cell type can adopt distinct functional states depending on the niche it resides in, reflecting local differences in signaling cues, extracellular matrix, and neighboring cell populations. For instance, in secondary lymphoid organs such as lymph nodes and spleen, the tissue can be anatomically partitioned into functionally distinct regions, including T cell zones, B cell zones, and germinal centers, where T cells, B cells, and antigen-presenting cells are respectively enriched. Within these regions, lymphocytes undergo extensive activation, proliferation, selection, and differentiation, illustrating how niche organization orchestrates immune cell fate and function. In small intestine, the epithelium is organized into large numbers of repetitive crypt-villi units, in which stem cells and differentiated terminal cells are spatially segregated to support continuous tissue renewal [8]. At the base of each crypt, Lgr5^+^ intestinal stem cells are intercalated with Paneth cells, which provide niche signals such as Wnt ligands, EGF, and Notch ligands that maintain stemness and proliferative capacity [8, 9]. As progeny of these stem cells migrate upward along the crypt-villus axis, they exit the stem cell niche, experience decreasing Wnt and increasing BMP signaling, and progressively differentiate into absorptive enterocytes, goblet cells, and enteroendocrine cells. Notably, Paneth cells are specifically restricted to the crypt base and do not appear in villi, creating a sharply delimited stem cell compartment that ensures both homeostatic turnover and rapid regeneration after injury.

Computationally, niche identification, segmentation and characterization are most commonly implemented by applying spatial domain identification algorithms to spatial transcriptomics data [10]. These methods cluster cells or spots into spatially coherent spatial domains, usually manifest as continuous patches in a tissue slice, based on joint patterns of gene expression, spatial proximity, and sometimes histological features, and these domains are then interpreted as putative tissue niches [10, 11]. Once domains are defined, downstream analyses, including cell type deconvolution, niche-specific differential expression, and ligand-receptor or pathway enrichment analysis, are used to functionally characterize each niche and link it to underlying biological processes. Although individual methods differ in how they represent transcriptomes and define similarity, most follow a shared design principle: they explicitly balance transcriptomic homogeneity, spatial continuity, and similarity in local cell-type composition to obtain biologically meaningful domains.

While related, in our view spatial domain segmentation and niche identification are not entirely equivalent. Spatial domain segmentation is fundamentally a partitioning operation in which cells or spots are assigned to non-overlapping regions, with a primary emphasis on maximizing intra-domain similarity in cell composition and transcriptional profiles while enhancing divergence between domains. Niche identification, by contrast, centers on characterizing microenvironmental function, intercellular signaling, and context-dependent cell states [12, 13]. A tissue niche does not necessarily encompass all cells within a local tissue pocket, and its defining interactions may involve only specific subsets of neighboring cells. Consequently, functionally distinct niches can overlap within the same physical region of tissue, effectively superimposing multiple microenvironments in the same anatomical space. Likewise, tissue niches are less constrained by spatial continuity, and the same niche type can naturally recur as multiple, spatially scattered microenvironments across a tissue section [13]. Current literature extensively validates spatial clustering through its concordance with histological structures, but it remains unclear whether this success translates to the identification of functional tissue niches, particularly in heterogeneous environments lacking discrete anatomical landmarks. Here, we systematically evaluate 16 domain segmentation algorithms for their ability to delineate nuanced niches formed by biochemical signaling gradients rather than physical barriers, such as basement membranes. We investigate the capacity of these algorithms to autonomously recapitulate biological knowledge and quantify their sensitivity to extrinsic guidance, including feature selection, spatial resolution, and the strategic weighting of key cell types.

First, we performed a comprehensive expert annotation of large-scale, single-cell resolution, FISH-based spatial transcriptomics data from human lymph nodes undergoing reactive follicular lymphoid hyperplasia (RFH). By leveraging the distinct zonal enrichment of key cellular lineages, we manually partitioned the tissue into the medulla, T-cell zone, and B-cell follicles. Within the B-cell follicles, we further delineated overlapping functional niches, specifically focusing on germinal center and B-cell maturation zones. Our evaluation revealed that while most domain segmentation algorithms effectively maximize intra-domain homogeneity and inter-domain disparity, the resulting boundaries consistently deviated from our expert-curated ground truth. This discrepancy is likely driven by the confounded signals from irregular and pervasive peripheral cell populations.

Next, we investigated whether domain segmentation could be refined through extrinsic guidance by mitigating the interference of less informative peripheral cell populations. While most algorithms exhibited varying degrees of sensitivity to these augmentations, we found that strategic weighting of key cell lineages enabled GraphST and MENDER to produce segmentations that closely recapitulated our expert-curated ground truth. This observation underscores a fundamental divergence between standard domain segmentation, which prioritizes global transcriptomic homogeneity across all cells, and functional niche identification, which necessitates the prioritization of highly conserved local microenvironments defined by a core set of lineage-specific cell types. Our *in silico* spatial simulations further reinforced this finding, demonstrating that without targeted correction, domain segmentation algorithms remain highly susceptible to interference from pervasive, non-canonical peripheral cell populations.

To assess the platform-agnostic performance of these algorithms, we extended our benchmark to include public 10x Visium and Stereo-seq datasets. Our analysis revealed significant variance in algorithmic accuracy across platforms, highlighting a critical linkage between computational performance and the underlying sequencing chemistry. Interestingly, our results show that intra-domain heterogeneity often outweighs inter-domain disparity. This finding suggests that the transcriptomic signatures of tissue domains are inherently noisy due to diverse cellular compositions, presenting a persistent challenge for algorithms that rely on maximizing global cluster separation.

Finally, we benchmarked the computational scalability of current algorithms, revealing that only a select few are equipped to process datasets exceeding one million observations, a scale already attainable with contemporary high-resolution platforms. Our findings provide a critical assessment of algorithmic utility for the burgeoning era of ultra-large-scale spatial transcriptomics. These results highlight the urgent need for computationally efficient frameworks that can maintain biological resolution without succumbing to the prohibitive memory and time requirements of burgeoning dataset volumes.

In summary, this work addresses a pivotal question in the field of spatially resolved transcriptomics: can standard domain segmentation algorithms be readily adapted for niche identification within complex tissues? Our results demonstrate that in environments where niches are defined by selective subsets of key cellular lineages, their transcriptomic signals are frequently obscured by the prevalence of less cohesively distributed peripheral cells. We show that without the strategic weighting of these core lineages, conventional algorithms fail to recapitulate functional architectures. Coupled with our manually curated RFH annotation, we envision that these findings will provide a foundation for the next generation of spatial algorithms, which are capable of navigating the nuanced divergence between global domain partitioning and targeted niche identification.

## 2 Results

### 2.1 Overview of the benchmarking task and framework

Spatial niche identification aims to partition tissue into multicellular microenvironments that are defined by coordinated cell type composition and spatially structured cellular states, using spatially resolved expression measurements. Across tissues, such microenvironments can reflect recurrent cellular neighborhoods, context-dependent state programs, and local cell-cell interactions [14] that are not fully captured by gene expression alone or by coarse anatomical landmarks. Tissue niches may align with the anatomical structure of a tissue in some settings, but they can also be sharply separated yet internally heterogeneous, appear as non-contiguous islands embedded within larger compartments, or vary continuously along gradients. These properties make it difficult to infer method performance from a single reference setting, motivating a benchmark that jointly probes multiple niche geometries and practical data regimes under a consistent task definition. (Fig. 1).

**Fig. 1.**
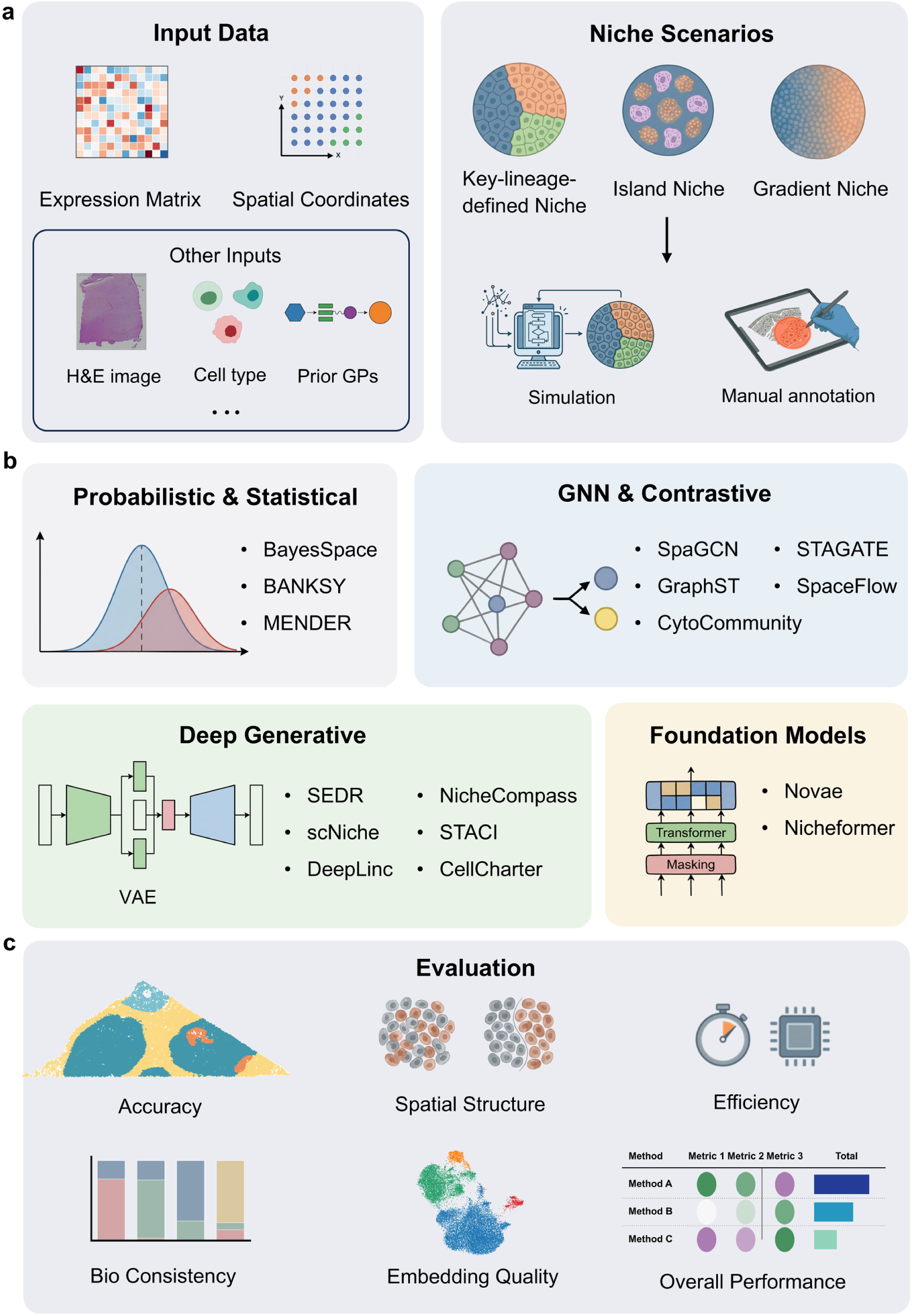
Benchmarking framework for spatial niche identification. (a) Inputs and benchmark scenarios. Expression matrix and spatial coordinates are treated as the minimal common inputs across methods, while optional auxiliary information may include histology images, cell-type annotations, or prior gene programs. To reflect diverse niche organization in tissues, we evaluate three canonical niche geometries: key-lineage-defined niches with sharp boundaries, island niches with spatially separated microdomains, and gradient niches with continuous transitions, using both simulation and expert manual annotation. (b) Method landscape. Sixteen niche identification methods are grouped into four methodological families: probabilistic and statistical models, GNN and contrastive methods, deep generative models, and foundation models. (c) Evaluation suite. Methods are assessed along complementary metrics, including agreement with reference niches (accuracy), spatial structure and boundary fidelity (connectivity), biological consistency of inferred niches (composition similarity), quality of learned embeddings (silhouette score), and computational efficiency (runtime and memory), which together support an overall performance summary for cross-method comparison.

We treat the expression matrix and spatial coordinates as the minimal common inputs across methods, while explicitly accommodating approaches that incorporate auxiliary modalities such as histology images, cell-type annotations, or prior gene programs (Fig. 1a). We benchmark 16 representative algorithms spanning probabilistic and statistical models (BayesSpace [15], BANKSY [16], MENDER [17]), GNN (graph neural network) [18] and contrastive methods (SpaGCN [11], GraphST [19], CytoCommunity [20], STAGATE [21], SpaceFlow [22]), deep generative models (SEDR [23], scNiche [24], DeepLinc [25], NicheCompass [12], STACI [26], CellCharter [27]), and foundation models (Novae [28], Nicheformer [29]) (Fig. 1b; Table 1). Probabilistic and statistical models typically encode spatial dependence through explicit priors or structured regularization and infer niches by optimizing likelihood-based or composition-based objectives. GNN and contrastive methods learn spatially informed embeddings by message passing on neighborhood graphs, often coupled with smoothness constraints or contrastive alignment to preserve local structure. Deep generative models use latent-variable formulations to capture complex, nonlinear variation and denoise technical or biological confounding before deriving niche partitions from the learned latent space. Foundation models transfer representations learned through large-scale pre-training across tissues, gene panels, and technologies. Novae learns local cellular-neighborhood graph representations using a prototype-based self-supervised objective adapted from SwAV and supports zero-shot spatial-domain assignment. Nicheformer instead applies masked-token pre-training to expression-ranked gene sequences from more than 110 million dissociated and spatially profiled cells; spatial coordinates are not used during pre-training, and spatial labels are learned through downstream probing or fine-tuning. Here, foundation model therefore refers to the large-scale pre-training and transfer paradigm.

**Table 1.** Overview of the 16 niche identification methods evaluated in this study.

| Method | Language | Core model | Inputs | GPU | Year |
| --- | --- | --- | --- | --- | --- |
| BayesSpace [15] | R | Bayesian Markov Random Field (MRF) | expression, coordinates | no | 2021 |
| BANKSY [16] | Python/R | Spatial feature augmentation | expression, coordinates | no | 2024 |
| MENDER [17] | Python | Multi-scale neighborhood representation | cell type, coordinates | no | 2024 |
| SpaGCN [11] | Python | Graph Convolutional Network (GCN) | expression, coordinates, histology | yes | 2021 |
| GraphST [19] | Python | Graph self-supervised contrastive learning | expression, coordinates | yes | 2023 |
| CytoCommunity [20] | Python+R | MinCut-based GNN | cell type, coordinates | yes | 2024 |
| STAGATE [21] | Python | Graph Attention Auto-encoder | expression, coordinates | yes | 2022 |
| SpaceFlow [22] | Python | Spatially regularized Deep Graph Networks | expression, coordinates | yes | 2022 |
| SEDR [23] | Python | VGAE + masked self-supervised learning | expression, coordinates | yes | 2024 |
| scNiche [24] | Python | Multi-view GAE + mutual-information maximization | expression, coordinates, cell type | yes | 2025 |
| DeepLinc [25] | Python | Variational Graph Autoencoder (VGAE) | expression, coordinates | yes | 2022 |
| NicheCompass [12] | Python | Variational Graph Autoencoder (VGAE) | expression, coordinates, gene programs | yes | 2025 |
| STACI [26] | Python | Graph-based autoencoder | expression, coordinates, images | yes | 2022 |
| CellCharter [27] | Python | Variational Autoencoder | expression, coordinates | yes | 2024 |
| Novae [28] | Python | Graph-based foundation model | expression, coordinates | yes | 2025 |
| Nicheformer [29] | Python | Transformer-based foundation model | expression, coordinates, metadata | yes | 2025 |

Beyond these broad model families, the methods differ in the spatial scale and biological signal that their objectives preferentially preserve. BayesSpace, SpaGCN, SpaceFlow, and SEDR place substantial weight on agreement between neighboring observations and are therefore expected to favour smooth, contiguous territories, potentially absorbing small embedded structures when their molecular contrast is weak. GraphST, STAGATE, and STACI retain finer local variation through contrastive learning, adaptive attention, or multimodal graph reconstruction, making them better suited to compact microdomains and sharper boundaries; BANKSY and CellCharter similarly use neighborhood-augmented representations that can expose local subdivisions. MENDER, scNiche, and CytoCommunity are driven more directly by local cell-type context and thus favour recurrent neighborhoods with similar cellular composition, whereas NicheCompass can prioritize signaling-related gene programs. Novae transfers local graph-neighborhood representations, while Nicheformer is primarily transcriptomic during pre-training and may consequently follow strong cell-state variation in the absence of downstream spatial supervision. These tendencies are not deterministic: their expression depends on neighborhood scale, input features, spatial regularization, and clustering resolution.

Performance is quantified along complementary axes (Fig. 1c), including agreement with reference niches, spatial coherence and boundary fidelity, biological consistency of inferred microenvironments, utility of learned embeddings, and practical runtime and memory costs. The results are organized to reflect how niche identification is used in practice. We first establish a baseline on a manually annotated CosMx [30] human lymph node reference (19,718 cells), and then dissect how input preprocessing modulates identifiability by evaluating feature selection, pseudo-spot aggregation, and core cell type refinement as distinct data processing strategies, each yielding a corresponding benchmark input. We next quantify performance on island-like niches by evaluating the ability to detect compact, embedded germinal center regions within B cell zone subregions drawn from two distinct lymph node areas (5,521 and 7,319 cells). On the 5,521-cell subset, we additionally assess sensitivity to gradient-like microenvironments by testing whether methods recover a continuous transition in B cell states along a spatial trajectory. In parallel, we use controlled lymph node simulations with increasing diffusion to emulate progressively mixed niches (4,000 cells per setting) and quantify robustness as niche boundaries become less separable. To assess generalizability beyond immune microenvironments, we evaluate methods on anatomy-driven references, including a human dorsolateral prefrontal cortex (DLPFC) [31] and a MOSTA [32] E16.5 mouse brain. Finally, we perform a dedicated scalability analysis using six dataset sizes spanning 20k to 1.8M cells, enabling feasibility comparisons under standardized resource constraints and highlighting which approaches remain practical at atlas scale.

### 2.2 Annotation and curation of a high-resolution human lymph node reference from a key lineage perspective

Lymph nodes are central hubs of adaptive immunity, strategically positioned along lymphatic vessels to filter lymph-borne material and coordinate when and where immune cells meet, enabling efficient immune surveillance and response initiation (Fig. 2a) [33]. Their hallmark is a highly ordered yet dynamic spatial architecture, in which stromal scaffolds and local molecular cues compartmentalize immune processes into distinct microenvironments that can rapidly remodel during inflammation. This combination of stereotyped organization and stimulus-driven reconfiguration makes lymph nodes an especially valuable tissue for studying niches, where spatial context is tightly coupled to multicellular function [34].

**Fig. 2.**
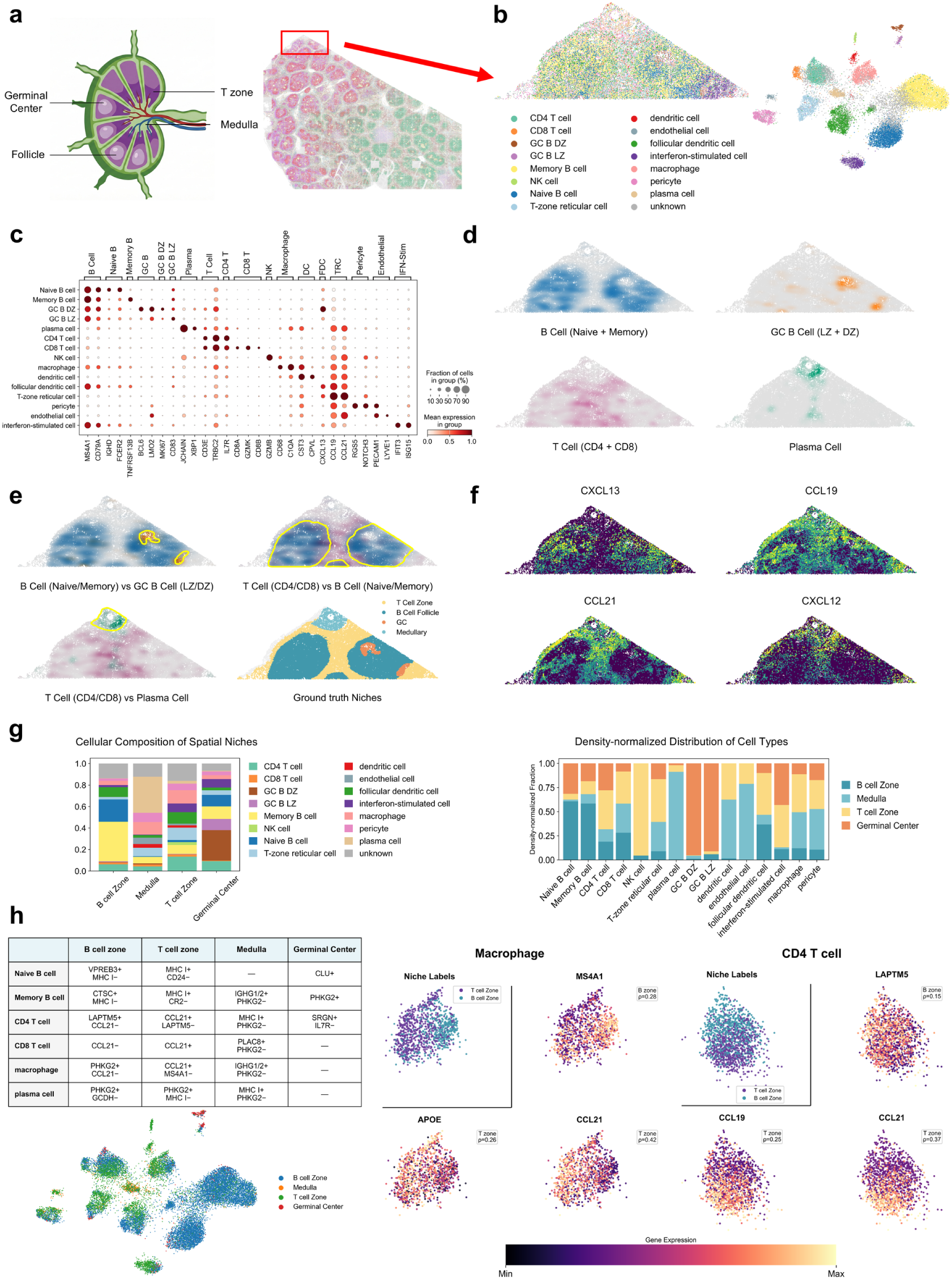
Annotation of cellular identities and canonical niches in the CosMx human lymph node dataset (a) Schematic of lymph node spatial architecture (left) and spatial cell-type map of the full CosMx human lymph node dataset (right). (b) Selection of a representative region (n=19,718 cells) (left) and UMAP of 16 refined cell types for cells in the selected region (right), displayed with a shared legend. (c) Dot plot of differential marker gene expression across subtypes. (d) Spatial density maps of representative cell types. (e) Lineage-specific density maps and spatial visualization of the four manually annotated niches (B cell zone, GC, T cell zone, and Medulla); yellow outlines indicate manually annotated niches (inset). (f) Spatial expression patterns of canonical chemokines (CXCL13, CCL19, CCL21, and CXCL12) highlighting the molecular basis of lymph node zonation. (g) Cellular composition of defined niches (left) and the density-normalized spatial distribution of each cell type across niches (right). (h) Niche-specific expression analysis on multiple cell types. Left: table of niche-enriched markers by cell type and a UMAP plot across all cells colored by niche type (+/− denote upregulation/downregulation vs. all other regions). Right: UMAP plots of macrophages and CD4 T cells colored by niche type (top-left plot), and the expressions of niche-specific genes, with Spearman’s *ρ* attached in legend.

We analyzed a CosMx spatial transcriptomics dataset comprising *∼*1.85 million cells, from which we curated a representative region of interest (ROI) of 19,718 cells spanning all major functional zones (Fig. 2a). This dataset was derived from human lymph node tissue exhibiting reactive follicular lymphoid hyperplasia (RFH), a common benign and reversible adaptive response to immune stimulation that reflects the capacity of the humoral immune system to remodel upon antigen challenge [35]. Under persistent stimulation, RFH is characterized by compensatory expansion of lymphoid follicles and associated stromal remodeling and cellular migration, yet it preserves the fundamental lymph node architecture, distinguishing it from malignant lymphoproliferative disorders such as lymphoma [36]. Anatomically, these activation-driven spatial patterns are readily apparent in our tissue section (Fig. 2b): plasma cells showed near-complete accumulation within the medulla and medullary cords, whereas naïve B cells expanded outward from the mantle zone and partially encroached upon memory B cell territories, consistent with follicular remodeling during sustained stimulation. Together, these configurations suggest progression toward a late-stage response in which antigen presentation is attenuated and the system shifts toward an effector program dominated by antibody production, with antibody egress likely concentrated through the medullary cords. In parallel, antigen-presenting regions, primarily follicles, appear to enter a restoration phase involving local homeostatic rebalancing, factor recycling, and stromal reorganization.

In anatomical terms, classical histology partitions the lymph node into distinct structural zones with specific functions: the follicle (cortex/B cell zone), the T cell zone (paracortex), and the medulla comprising medullary cords and sinuses (Fig. 2a) [34]. During immune activation, follicles expand to form germinal centers (GCs) [37], a feature notably pronounced in RFH. These primary zones contain specialized sub-compartments: the GC, serving as the site of antigen-specific clonal expansion; the mantle zone, a circumferential ring of displaced naïve B cells [37]; and the T-B border, where activated B cells upregulating *CCR7* and *GPR183*/*EBI2* interact with pre-Tfh cells, a decisive checkpoint governing GC entry [38]. The medulla comprises medullary cords, where plasma cells reside and secrete antibodies, and medullary sinuses serving as channels for lymph and antibody egress [34]. The GC is functionally polarized into a dark zone (DZ), where centroblasts undergo clonal expansion and somatic hypermutation, and a light zone (LZ), where centrocytes undergo affinity selection via interactions with FDCs and Tfh cells; only centrocytes that successfully capture antigen receive the critical survival signals CD40L and IL-21 [39]. In parallel, lymph flow imposes a coordinated functional itinerary: antigen-bearing lymph is screened at the subcapsular sinus and relayed via B cells to FDC networks, routed through the paracortical conduit system for small-molecule surveillance, and ultimately accumulated in the medulla for export [34].

We manually annotate the immune niches by first annotating all the cell types in the ROI region. We performed canonical clustering followed by differential expression analysis and canonical marker-based annotation. The established cell type annotation in the ROI is then used as the foundation for subsequent niche delineation. The cell type annotation criteria are summarized in Fig. 2c. Briefly, we classify cells to the following cell types: T lymphocytes, marked by *CD3E*/*TRBC2*, were subdivided into CD4^+^ helper T cells (*IL7R*) and CD8^+^ cytotoxic T cells (*CD8A*/*CD8B*/*GZMK*). B lymphocytes, marked by *MS4A1*/*CD79A*, were stratified into naïve B cells (*IGHD*/*FCER2*), germinal-center B cells (*BCL6*/*LMO2*) with dark-zone cells further indicated by *MKI67* and light-zone cells by *CD83*, memory B cells (*TNFRSF13B*), and plasma cells (*JCHAIN* /*XBP1*). Myeloid cells were predominantly macrophages (*CD68*/*C1QA*). Stromal populations included T-zone reticular cells (TRCs; *CCL19*/*CCL21*). Follicular dendritic cells (FDCs; *CXCL13*), which provide positional cues that structure the paracortex and follicle, respectively.

By conventional knowledge, lymph node compartments are not separated by membrane-like physical barriers but emerge from spatially organized biochemical gradients and stromal scaffolds [40]. We therefore treated each niche as being organized around a small set of function-defining cell populations and delineated canonical regions by mapping their spatial density fields (Fig. 2d, e). Specifically, we used naïve and memory B cells for follicle annotation, GC B cells to mark the germinal center, CD4 and CD8 T cells to delineate the paracortex, and plasma cells to identify the medulla. As shown in Fig. 2e, the above key lineages are enriched in mostly mutually exclusive regions, with their boundaries as natural cues for segmentation (Fig. 2e). The preferential spatial localization of the above cells is likely a result of concerted molecular signaling in the lymph node for immune cell directed migration. Fig. 2f demonstrates the spatial expression strength of key chemokines: *CXCL13* is enriched in the B cell compartment and supports follicular segregation by recruiting *CXCR5*^+^ lymphocytes; *CCL19* and *CCL21* align with the T cell zone to promote *CCR7* -dependent positioning and T cell-APC encounters, complemented by a paracortical *CXCL12* cue that provides additional *CXCR4*-mediated guidance for lymphocyte trafficking [41].

While the immune niches are not physically separated by membrane barriers, their specific biochemical-driven enrichment and interleaved organization is functionally important. Within each niche, these organizing populations participate in tightly coupled cooperation networks. In the follicle, antigen is relayed from subcapsular sinus macrophages to B cells and then to follicular dendritic cell networks that retain immune complexes and support iterative B-cell scanning, with memory B cells near the subcapsular periphery poised for rapid recall responses [42]. In the T cell zone, fibroblastic reticular cells provide a conduit scaffold for antigen and chemokine transport, and *CCL19*/*CCL21* tune T-cell motility to increase productive contacts with antigen-presenting cells. The medulla couples egress with humoral maintenance, as plasma cells occupy medullary cords supported by stromal survival cues and medullary sinus macrophages mediate terminal filtration. In germinal centers, affinity maturation follows the dark-zone-to-light-zone cycle, in which B cells diversify and are selected through competition for follicular dendritic cell-presented antigen and T follicular helper signals [37]. Although key cell types exhibit clear spatial enrichment that supports regional annotation, they do not necessarily constitute an absolute or even relative majority within their respective zones because niche function depends on extensive collaboration with auxiliary populations. For instance, CD4 and CD8 T cells do not dominate the cellular composition of the T cell zone (Fig. 2g), yet density maps show that most T cells concentrate there at markedly elevated local density, underscoring that niche identity is often encoded by lateral spatial distribution rather than vertical abundance. Similarly, while GC B cells, including both dark-zone and light-zone variants are predominately enriched in GC zone, their presence in GC registers only a moderate relative majority, diluted by the participation of cell types across the board. As we will return in later sections, the awkward symbiosis of evident preferential localization of key lineages in tissue while taking only a minor role in local cell type compositions is a major challenge in niche identification through domain segmentation.

Finally, we observe non-trivial within-niche transcriptomic similarity and between-niche disparity among cells of the same type. This indicates functional specialization shaped by the local microenvironment. The niche-specific expressions are summarized in the table of Fig. 2h. Among cell types with significant inter-niche differential expressions, we observe macrophages with the most exemplified plasticity: follicular macrophages upregulate *PHKG2* and *MS4A1*, consistent with active phagocytic engagement with B-cell material; T-zone macrophages express *APOE* together with *CCL21*, aligning with debris clearance and reinforcement of paracortical chemokine structure [43]; and medullary macrophages express *IGHG*, potentially reflecting clearance of antibody-rich material [44]. CD4 T cells similarly show spatial adaptation, with follicular CD4 T cells expressing *LAPTM5* consistent with restrained activation [45], paracortical CD4 T cells expressing *CCL19* and *CCL21*, medullary CD4 T cells exhibiting elevated MHC class I-related signatures, and GC CD4 T cells corresponding to the Tfh subset expressing *SRGN* (Fig. 2h). Additional information with regard to the niche-specific expression of other cell types can be found in Supplementary Fig. S1. It is worth noting that while there is non-trivial niche-specific expression within cell types and could potentially be harnessed for niche extraction, the differences are nuanced and easily surpassed by cell type differences between local microenvironment. Thus, cell composition and concentration shifts remain the dominating forces guiding niche annotation.

### 2.3 Functional niche curation from a cross-lineage proximal gene co-expression perspective

Aside from tissue niche definition based on distinct localization of key lineages, the curation of tissue niches is also supported by conserved proximal gene co-expressions arise from multiple lineages. Here, proximal co-expression does not require genes to be expressed within the same cell; instead, it refers to lineage-restricted genes whose expression repeatedly occurs in close spatial proximity across the tissue. From this perspective, a functional niche is jointly characterized by its spatial context, the niche-associated genes active within that context, and the specialized cell states that carry these genes. This provides a complementary view to the key-lineage-based annotation by asking whether distinct cellular populations converge spatially on a coherent functional program.

At the molecular level, the lymph-node tissue contains several distinct cross-lineage functional programs rather than signals attributable to a single marker or dominant cell population. Four of these programs correspond closely to the major recurrent lymph-node architecture, while an additional interferon-responsive program captures a localized condition-dependent state. We curated 30 representative genes for each niche according to their spatial distributions and associated biological functions, with selected genes shown in Fig. 3a. The reticular-immune support program niche combines lymphocyte-associated *IL7R*/*GIMAP7*, myeloid *FCER1G*/*C1QA*, and TRC-associated *CCL21*, *CXCL12*, *COL1A1*, and *DCN*, jointly reflecting immune-cell positioning, surveillance, clearance, and stromal support. Follicular B-cell organization niche integrates B-cell signaling and state-associated genes such as *BANK1*, *PLCG2*, *TLR10*, *P2RX5*, and *CR2* with the follicular stromal and antigen-presentation programs represented by *CXCL13* and *CIITA*. The medullary secretion-clearance program niche couples plasma-cell secretion (*JCHAIN*, *XBP1*, and *MZB1*) with macrophage-mediated immunoglobulin handling and scavenging (*FCGRT*, *STAB1*, and *CPVL*), endothelial-associated *RNASE1*/*STAB1*, and stromal *TNC*. Finally, the germinal-center reaction program niche combines GC B-cell activation and proliferation (*BCL6*, *RGS13*, *AICDA*, and *MKI67*) with helper-T-cell activity (*PDCD1* and *ICOS*), FDC-associated *CR2*, and stromal *SERPINE2*. In addition, a spatially restricted interferon-responsive program is marked by coordinated expression of canonical interferon-stimulated genes, including *IFIT2*, *IFIT3*, *MX1*, *RSAD2*, *ISG15*, *OAS1*, *PLSCR1*, and *STAT1*, across immune, stromal, and vascular-associated populations. Unlike the four architecture-associated programs, this pattern represents a condition-dependent inflammatory state superimposed on the underlying lymph-node organization. Their related but non-identical spatial distributions indicate that each niche represents a coordinated functional program assembled across several lineages.

**Fig. 3.**
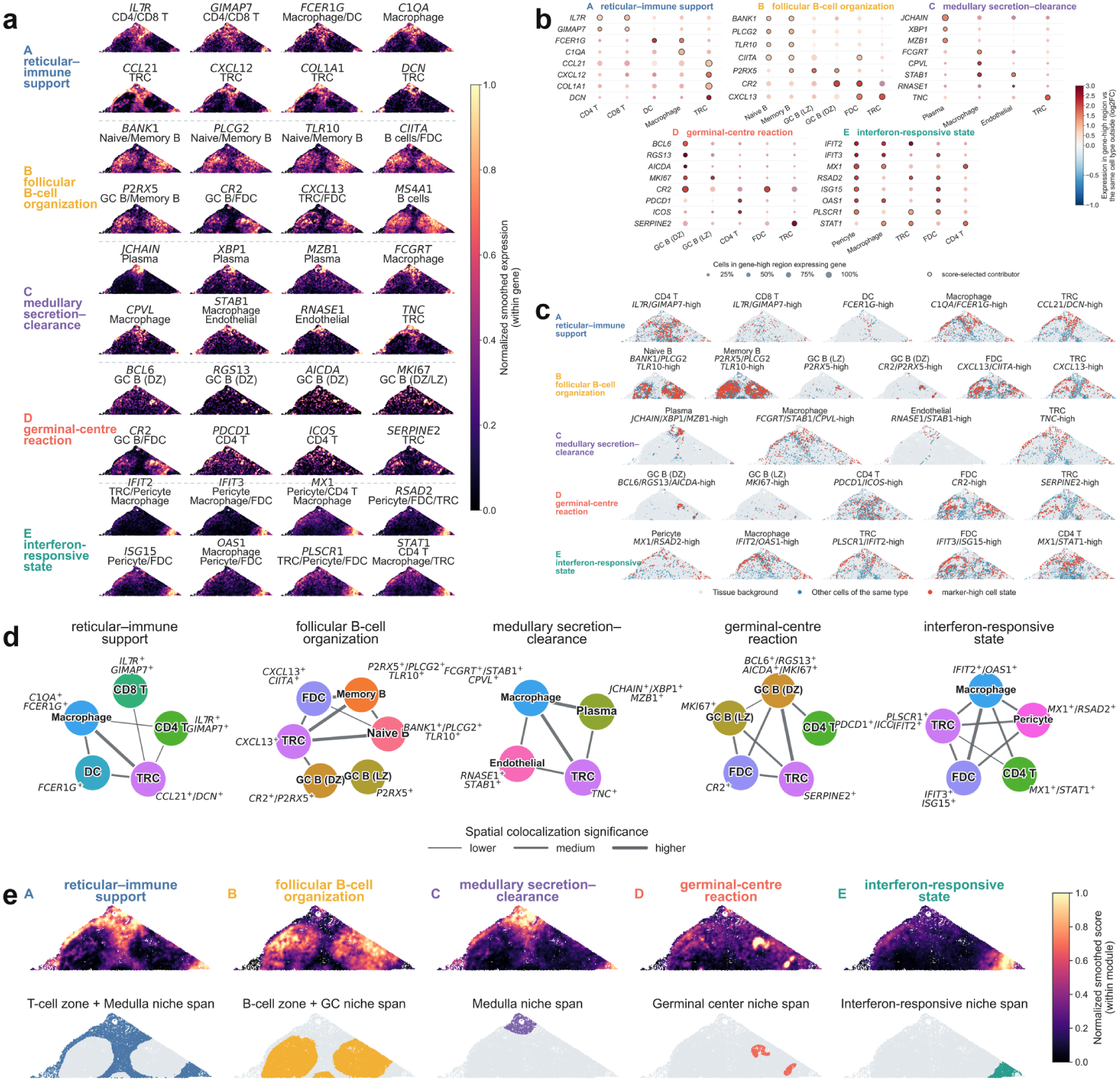
Multicellular and molecular characterization of functional niches from a cross-lineage proximal gene co-expression perspective. (a) Spatial expression of representative genes contributing to four architecture-associated niche programs and an additional interferon-responsive state: reticular–immune support, follicular B-cell organization, medullary secretion–clearance, germinal-center reaction, and interferon response. Representative genes arise from multiple cell lineages and show coordinated spatial distributions. (b) Cell-type-resolved expression of niche-associated genes. Dot size indicates the fraction of cells in the corresponding gene-high region expressing the gene, and colour denotes expression enrichment relative to cells of the same type outside that region; outlined dots indicate score-selected contributors. (c) Spatial distributions of niche-associated cell states defined by representative gene combinations. Marker-high cells are shown in red, other cells of the same type in blue, and tissue background in grey. (d) Multicellular co-localization networks for the five functional programs. Nodes represent niche-associated cell states, and thicker edges indicate stronger statistical support for spatial co-localization. (e) Continuous spatial activity of the five gene modules (top) and their corresponding spatial spans (bottom). The first four programs broadly follow the major lymph-node architecture, whereas the interferon-responsive state forms a localized functional microenvironment superimposed on the T-cell-zone context, illustrating functional organization beyond discrete anatomical partitions.

Importantly, these cross-lineage programs are carried by specialized states within their parent cell populations rather than uniformly by entire cell types. Cell-type-resolved analysis showed that niche-associated genes were preferentially enriched in restricted subsets of the corresponding lineages (Fig. 3b). In the reticular-immune niche, for example, *IL7R*/*GIMAP7* - high CD4 and CD8 T cells coexist with *FCER1G*-high dendritic cells, *C1QA*/*FCER1G*-high macrophages, and *CCL21*/*DCN* -high TRCs. Follicular organization niche similarly involves distinct naïve-, memory-, and GC-B-cell states together with *CXCL13*/*CIITA*-high follicular stromal populations. In the medulla region, *JCHAIN* /*XBP1*/*MZB1*-high plasma cells are accompanied by specialized macrophage, endothelial, and TRC states, whereas the germinal-center program combines activated or proliferative GC B cells with *PDCD1*/*ICOS*-high CD4 T cells, *CR2*-high FDCs, and *SERPINE2*-high TRCs. The interferon-responsive program provides an especially clear example of such cross-lineage state specialization, comprising *MX1*/*RSAD2*-high pericytes, *IFIT2*/*OAS1*-high macrophages, *PLSCR1*/*IFIT2*-high TRCs, *IFIT3*/*ISG15*-high FDCs, and *MX1*/*STAT1*-high CD4 T cells (Fig. 3b). Thus, functional niche identity reflects context-dependent specialization within lineages in addition to their regional abundance; the shared interferon response further demonstrates that a local functional state can arise synchronously across otherwise distinct cell populations.

These specialized states, in turn, occupy coordinated but non-identical spatial territories, providing a direct spatial link between lineage-specific transcriptional states and local niche organization. As shown in Fig. 3c, marker-high cells frequently occupy restricted subsets of the broader distribution of their parent cell types. Follicular naïve-, memory-, and GC-B-cell states are spatially accompanied by follicular stromal states, while the medulla contains highly localized antibody-secreting plasma cells embedded among more broadly distributed macrophage, endothelial, and stromal states associated with clearance and transport. Within germinal centers, specialized GC B-cell states form compact foci, whereas helper T-cell, FDC, and stromal states occupy complementary distributions within and around these structures. A similar but more condition-dependent pattern is observed for the interferon-responsive states, which were concentrated in a localized portion of the tissue despite the broader distributions of their parent cell types. Pericyte, macrophage, TRC, FDC, and CD4 T-cell subsets expressing the interferon stimulation genes (ISG), which indicate the nucleus proliferation of ISGF3, a heterotrimer assembled by combining *STAT1*, *STAT2* and *IRF9* downstream of the *JAK*-*STAT* pathway activation by interferons. The ISGF3 binds directly to the ISRE motif, which is an enhancer upstream of most ISG genes that activates the ISG expression. The converged ISG spatial expressions within this region, signify a locally coordinated collective response of interferon stimulation rather than changes in cellular identity and composition. The same broad cell type can therefore practice different biological functions across tissue contexts, while selected states from distinct lineages repeatedly converge within related microenvironments. At the multicellular level, these spatially coordinated states form coherent niche-associated assemblies rather than independent collections of marker-high cells. Spatial co-localization analysis revealed statistically supported networks connecting complementary cell states from different lineages within each niche program (Fig. 3d). The reticular-immune network links lymphocyte states with TRCs, dendritic cells, and macrophages; the follicular network connects B-cell states with FDC- and TRC-associated states; the medullary network couples plasma-cell secretion with macrophage clearance, endothelial transport, and stromal support; and the germinal-center network brings together GC B cells, helper T cells, FDCs, and TRCs. The interferon-responsive network similarly connected ISG-high macrophage, pericyte, TRC, FDC, and CD4 T-cell states, supporting a coordinated multicellular response shared across distinct lineages. These recurrent proximal relationships support a multicellular view of niche organization in which distinct lineages collectively contribute complementary local functions, conceptually consistent with the emerging framework of spatial ecotypes as recurrent multicellular neighborhoods composed of co-associated cell states [46], which has recently been operationalized at scale to identify conserved multicellular ecosystems across diverse human tumors [47]. Although spatial co-localization does not by itself establish direct molecular interaction between every connected pair, it demonstrates that these niches cannot be reduced to the presence of any single lineage or cell state.

At the tissue scale, the resulting functional niche activity is graded, locally heterogeneous, and hierarchically overlapping rather than confined to uniformly filled, mutually exclusive territories. We integrated the complete niche-associated gene sets into continuous spatial module scores and compared their distributions with the benchmark reference regions (Fig. 3e). Reticular-immune support niche extends across both the T-cell-zone and medullary reference regions, within which medullary secretion–clearance forms a more spatially concentrated program. Likewise, follicular B-cell organization spans both the B-cell-zone and germinal-center regions, whereas the germinal-center reaction forms compact high-activity foci nested within this broader follicular field. The interferon-responsive module showed a different form of organization: its activity was strongly concentrated in a localized region predominantly embedded within the broader T-cell-zone environment. Rather than defining an additional anatomical compartment, this pattern represents a condition-dependent functional state superimposed on the reticular-immune support architecture. The same location can therefore retain a broad tissue-organizing program while simultaneously participating in a more specialized or condition-dependent functional state, and the strength of these programs varies continuously even within regions assigned the same anatomical label.

This molecular and multicellular organization clarifies how the functional niche definition relates to the discrete reference used for benchmarking. The B-cell zone is predominantly characterized by follicular B-cell organization, whereas the germinal center retains this broader follicular program while adding the specialized germinal-center reaction. Similarly, the T-cell zone is dominated by reticular-immune support, while the Medulla combines this shared support environment with a more concentrated secretion-clearance program. The interferon-responsive niche represents a different, condition-dependent level of organization: it is locally superimposed on the broader T-cell-zone context rather than forming a mutually exclusive anatomical compartment. The cross-lineage gene programs, lineage-specific cell states, multicellular co-localization networks, and continuous activity maps thus provide a complementary functional view, revealing graded and overlapping niche organization that cannot be fully represented by a single hard partition.

### 2.4 Niche identification benchmarking on the lymph node spatial transcriptomics data

We benchmarked 16 niche identification methods on the manually annotated human lymph node dataset, using the complete set of 6,195 genes to maximize information retention. The results are summarized in Fig. 4. Overall, we observed a consistent theme across all approaches: while none of the methods fully recapitulated the manual reference, most produced spatial partitions that are internally coherent and biologically interpretable under their own modeling assumptions, highlighting a gap between the method-designated most appropriate segmentation through empirical deliberation between transcriptomic homogeneity and spatial proximity; and our biology-grounded niche definition encoded in the ground truth.

**Fig. 4.**
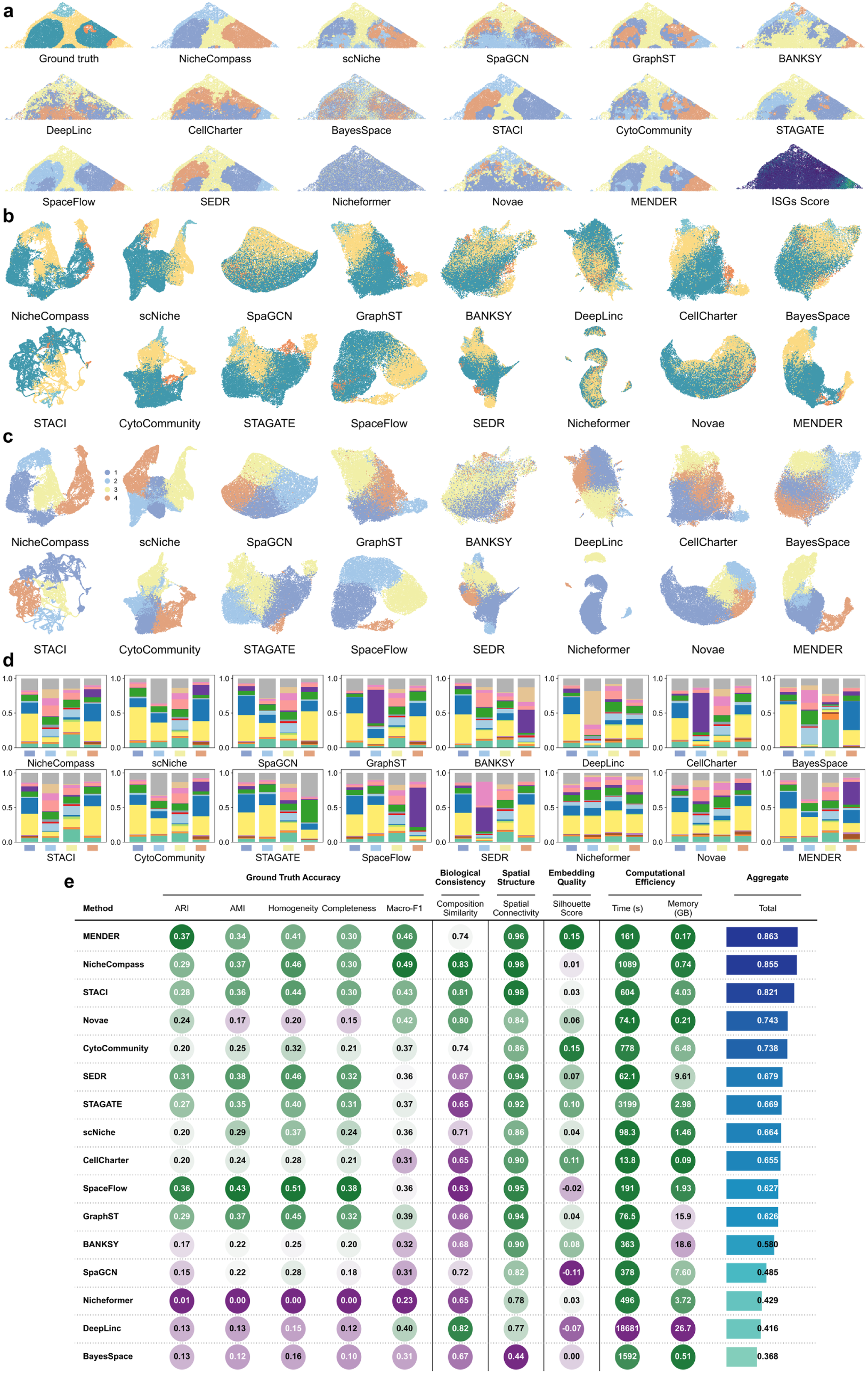
Benchmarking results on the manually annotated human lymph node dataset. (a) Spatial niche assignments for the ground truth and each method, visualized in tissue coordinates; the last panel shows the spatial map of ISGs score computed from interferon-stimulated genes (IFIT2, IFIT3, ISG15, MX1, OAS1, RSAD2) (b-c) UMAP visualization of method-specific latent embeddings, with cells colored by ground-truth niche labels (b) and by each method’s predicted niche labels (c). (d) Cell-type composition of niches for each method, shown as the proportion of annotated cell types within each niche. (e) Multi-criteria quantitative comparison across ground-truth accuracy, biological consistency, spatial structure, embedding quality, computational efficiency, and the aggregated overall score.

Across the spatial niche maps (Fig. 4a), several consistent qualitative patterns emerged. NicheCompass and STACI most clearly recapitulated the internal heterogeneity of the Medulla and T cell zone, in line with their relatively strong recovery of heterogeneous niche labels. Many other methods produced visually smooth partitions that respected broad compartmental continuity but collapsed this intra-compartment structure. At the compartment scale, multiple approaches, including NicheCompass, STACI, CellCharter, CytoCommunity, STAGATE, and SEDR, cleanly separated the central T cell zone from the flanking follicular B cell zone, yet often attribute the left and right follicular regions into separate niches, despite these bilateral regions corresponding to the same compartmental niche in the manual reference. The over-fragmentation in follicles is associated with reduced agreement in accuracy metrics. Out of all methods, MENDER was notably more robust, preserving cross-side concordance of the two B cell zone regions while maintaining the global compartment layout, consistent with its top ARI among methods. Germinal center identification remained challenging overall (Supplementary Fig. S9): from niche geometry and boundaries, only CellCharter and MENDER reliably recovered a coherent germinal-center-like region (typically capturing one complete GC), while most methods homogenized the GC with surrounding niches. These results suggest that germinal center identification in this dataset may require either more sensitive modeling of rare/compact structures or better disentanglement between local composition and broader compartmental context.

Cell-type composition profiles (Fig. 4d) further indicated that inferred partitions were often internally coherent and biologically interpretable even when they diverged from the manual niche reference. Across methods, predicted domains frequently showed sharper compositional contrasts than the ground-truth niches, suggesting that many algorithms preferentially partition tissue along the most separable cell-type or cell-state axes rather than the compartment-informed organization used for annotation.

One recurrent pattern was the emergence of domains dominated by strong, spatially localized cell-state programs, such as the interferon-stimulation response. In GraphST, BANKSY, CellCharter, SpaceFlow, and MENDER, a lower-right region characterized by elevated *IFIT2*, *IFIT3*, *ISG15*, *MX1*, *OAS1*, and *RSAD2* repeatedly influenced the inferred partitioning. Cross-method analysis confirmed that this was a reproducible functional-state signal rather than an algorithm-specific artifact: the consensus region was located predominantly within the manually annotated T-cell zone and showed coordinated ISG induction across multiple immune and stromal cell types, together with strong interferon-response pathway enrichment (Supplementary Fig. S17). Thus, this disagreement does not imply that the anatomical T-cell-zone annotation is incorrect; instead, several methods preferentially resolve a finer interferon-responsive microenvironment nested within that compartment. Because the benchmark requires a fixed four-domain partition, allocating one inferred domain to this localized state also alters the assignment of the remaining cells, thereby reducing agreement with the compartment-based reference.

We next evaluated whether the interferon-responsive state could be recovered directly from the existing domain partitions, rather than introducing it as a fifth mutually exclusive class in the primary benchmark. Representative best-matched domains identified by SpaceFlow, CellCharter, GraphST, SEDR, BANKSY, and MENDER are shown in Supplementary Fig. S19a. Using the 818-cell interferon-responsive region as an independent binary reference, we quantified the precision, recall, and F1 score of the best-matched domain from each method (Supplementary Fig. S19b). Under the predefined detection criteria, SpaceFlow, CellCharter, GraphST, SEDR, and BANKSY recovered the interferon-responsive state, whereas MENDER achieved partial recovery. SpaceFlow and CellCharter showed the strongest agreement with the reference (F1 = 0.90 and 0.88), followed by GraphST (0.74), SEDR (0.73), and BANKSY (0.67); MENDER reached an F1 of 0.54. Several other methods exhibited high recall but substantially lower precision, indicating that they covered much of the interferon-responsive region without isolating it as a spatially specific partition.

Other departures from the reference were more directly driven by dominant cell-type composition. For example, NicheCompass and BayesSpace produced domains with highly distinctive dominant populations, yielding readily interpretable memory-B-cell- or CD4-T-cell-enriched partitions. These observations reinforce an important distinction: strong compositional separation or high within-domain transcriptomic similarity does not necessarily correspond to functional niche delineation. Functional niches instead reflect coordinated, spatially organized multicellular programs that may cut across dominant cell-type boundaries or coexist with broader anatomical structure. Such composition- or transcriptome-driven partitions may still contain biologically relevant information, but they represent a different organizational axis from the niche definition evaluated here. This distinction underlies the broader difference between spatial domain segmentation and functional niche identification.

These contrasting segmentation patterns can be interpreted in light of the methods’ inductive biases. Because most approaches infer representations from local neighborhoods and subsequently impose a hard partition, spatially disconnected but biologically related structures are not explicitly constrained to share the same label. This helps explain why NicheCompass, STACI, CellCharter, CytoCommunity, STAGATE, and SEDR frequently assigned the bilateral follicular regions to different domains. At the opposite resolution limit, strong neighborhood smoothing or sensitivity to dominant tissue-scale variation can absorb compact GC islands into the surrounding B-cell field when their lineage-specific signal is locally diluted. The interferon-responsive domains reflect a distinct preference for high-contrast biological signals: GraphST, BANKSY, CellCharter, and SpaceFlow can organize their representations around the spatially coherent ISG program, whereas MENDER captures the multiscale enrichment of the interferon-stimulated cell population provided in its cell-type input. Thus, disagreement with the compartment-based reference can arise because a method preferentially resolves a different spatial scale or biological axis, rather than from a single uniform mode of segmentation failure.

Latent-space embedding showed a closely related picture. When each method’s embedding was visualized with UMAP [48] and colored by the ground-truth annotation (Fig. 4b), most methods exhibited substantial inter-niche mixing; only MENDER and CytoCommunity produced embeddings in which ground-truth niches formed comparatively coherent, separated groups (silhouette score, or SC, 0.15 and 0.14, respectively), whereas most other methods had silhouette scores generally *≤* 0.10. In contrast, coloring the same embeddings by each method’s predicted labels (Fig. 4c) typically yielded compact, well-separated clusters, indicating that many approaches learn internally consistent latent organizations even when those partitions only partially align with the manual reference. A supervised out-of-fold kNN read-out further showed that many embeddings retained reference-relevant information that was not recovered by the default clustering step (Supplementary Fig. S7); this diagnostic analysis was not used in the unsupervised benchmark rankings.

Finally, we quantitatively assessed the performance of all 16 domain segmentation algorithms across various metrics and their performances are recorded in the table in Fig. 4e. We evaluate the results from a diverse set of perspectives, including ground-truth accuracy, biological consistency, spatial structure, embedding quality, and computational efficiency. The resulting summary (Fig. 4e) indicates that the highest-ranked methods are those that perform consistently across these layers rather than optimizing a single criterion. MENDER shows the most balanced profile (aggregate 0.863), combining the strongest agreement with the biology-grounded manual reference (ARI 0.37; Macro-F1 0.46) with high spatial coherence (spatial connectivity 0.96) and the clearest ground-truth separation in latent space (SC 0.15), while remaining computationally efficient (161 s; 0.17 GB). NicheCompass (aggregate 0.855) and STACI (aggregate 0.821) follow closely. Both excel in spatial and compositional fidelity: each attains the top spatial connectivity (0.98), and both achieve high composition similarity (0.83 and 0.81, respectively). However, their overall agreement with the manual reference remains lower than MENDER (ARI 0.29 and 0.28), consistent with the boundary and fragmentation patterns observed in the spatial maps. Spatial segmentation by other methods topologically deviates from our annotation ground truth. The two foundation models showed distinct transfer behavior in the primary lymph-node benchmark. Novae remained competitive overall (aggregate score 0.743) and achieved moderate reference agreement (ARI 0.24; Macro-F1 0.42), within the ARI range of the GNN and contrastive methods, although below SpaceFlow, GraphST, and STAGATE. Nicheformer showed substantially lower agreement in the same unsupervised setting (ARI 0.01; Macro-F1 0.23). This contrast is consistent with Novae explicitly learning spatial-neighborhood graphs for zero-shot domain inference, whereas Nicheformer pre-trains cell-level transcriptomic representations without physical coordinates and typically learns spatial labels through downstream probing or fine-tuning. More broadly, large-scale pre-training provides transferable representations across heterogeneous datasets, but does not by itself ensure superior recovery of functional lymph-node niches. In this setting, methods trained directly on the target tissue can retain an advantage because their neighborhood message passing and spatial regularization are optimized for the specific topology of the tissue being analyzed. CytoCommunity reaches a similarly high silhouette score (0.15) but does not translate this embedding separability into comparable agreement (ARI 0.20). Practical considerations further differentiate methods: CellCharter is exceptionally lightweight (13.8 s; 0.09 GB), whereas DeepLinc is computationally demanding (18,681 s; 26.7 GB) despite its reasonable biological consistency.

Because global evaluation metrics can obscure structurally distinct errors, we further decomposed the four reference niches into their spatially connected regions and classified their recovery as single-domain, split, merged/hidden, or missed/fragmented (Supplementary Fig. S16). The connected-region analysis revealed distinct failure patterns that were not apparent from the aggregate scores: the broader B- and T-cell compartments were often substantially recovered but could be divided across multiple predicted labels, whereas the smaller Medulla and germinal-center regions were more susceptible to merging, fragmentation, or incomplete recovery. Evaluating cell-type composition only when a reliable spatial correspondence was present further separated niche recovery from compositional fidelity, providing a complementary diagnostic to the global Composition Similarity score.

We further assessed the robustness of the primary comparison to reference-boundary uncertainty and random initialization. Restricting evaluation to progressively more interior niche cores increased absolute agreement with the reference, while relative method rankings remained highly stable across erosion levels (Supplementary Fig. S20), indicating that mixed niche interfaces do not drive the principal cross-method conclusions. Across five pre-specified random seeds, MENDER retained the highest mean ARI (0.4181 *±* 0.0579), whereas SEDR (0.3442 *±* 0.1457) and GraphST (0.3340 *±* 0.1751) achieved the next-highest mean values but showed substantially greater between-seed variation. By comparison, SpaceFlow (0.3071 *±* 0.0339) and STACI (0.2774 *±* 0.0176) were more stable across initializations (Supplementary Fig. S14; Supplementary Tables S2 and S3). Thus, MENDER’s strong average agreement was reproducible, while small performance differences among several subsequent methods should not be interpreted as stable rank differences. We separately examined the sensitivity of the aggregate score to its weighting scheme. Although several reference-agreement metrics were strongly correlated, composition similarity, embedding quality, and computational efficiency captured less redundant aspects of performance. Across six prespecified weighting schemes, MENDER, NicheCompass, and STACI consistently formed the top-three group, while their internal ordering varied with the relative emphasis placed on accuracy, biological and spatial consistency, and efficiency. This leading group was also highly stable across 10,000 plausible weight combinations, with each of the three methods appearing in the top three in more than 90% of cases (Supplementary Fig. S13). Together, these robustness analyses support the aggregate score as a useful descriptive summary of overall multi-criteria performance, while emphasizing that exact method ordering and method-specific trade-offs are more appropriately interpreted from the individual component metrics.

### 2.5 Assessing the impact of feature selection, pseudo-spot aggregation, and core-cell-type-guided refinement on niche identification

Spatial domain segmentation is well documented to be highly volatile, sensitive to gene selection and technology platforms. Thus, we investigate if domain segmentation algorithms can rectify their results through extrinsic augmentations, including feature selection, resolution reduction and strategic weighing of key lineages. For feature selection, we benchmarked three strategies, each restricted to 2,000 genes: (i) HVG (highly variable genes), selecting the top 2,000 highly variable genes; (ii) SVG (spatially variable genes), selecting the top 2,000 spatially variable genes; and (iii) Curated genes, selecting 2,000 niche-informative genes derived from a core-cell-type-guided workflow in which dominant cell types were first used to define coarse groups and then subjected to group-versus-rest differential expression, with markers pooled and ranked to form a compact, niche-focused gene panel (Fig. 5a). Notably, when restricting the view to these dominant core cell types, niche-specific compositional signatures become substantially more distinct (Fig. 5c), providing a direct rationale for both the curated-gene panel and the core-cell-type-anchored refinement strategy. In addition to feature selection, we evaluated a distinct core lineage refinement strategy that leverages the same dominant cell types at the cell level rather than the gene level: niche identification was first performed on the curated subset of core cell types to obtain a stable core niche partition, after which remaining cells were assigned to these niches by K-nearest-neighbor-(KNN)-based label diffusion [49] (Fig. 5a). It is worth noting that we exempt MENDER and CytoCommunity from gene selection benchmarks, due to their unique workflows which perform domain segmentation not directly on the transcriptome but over cluster assignments (which is conducted separately).

**Fig. 5.**
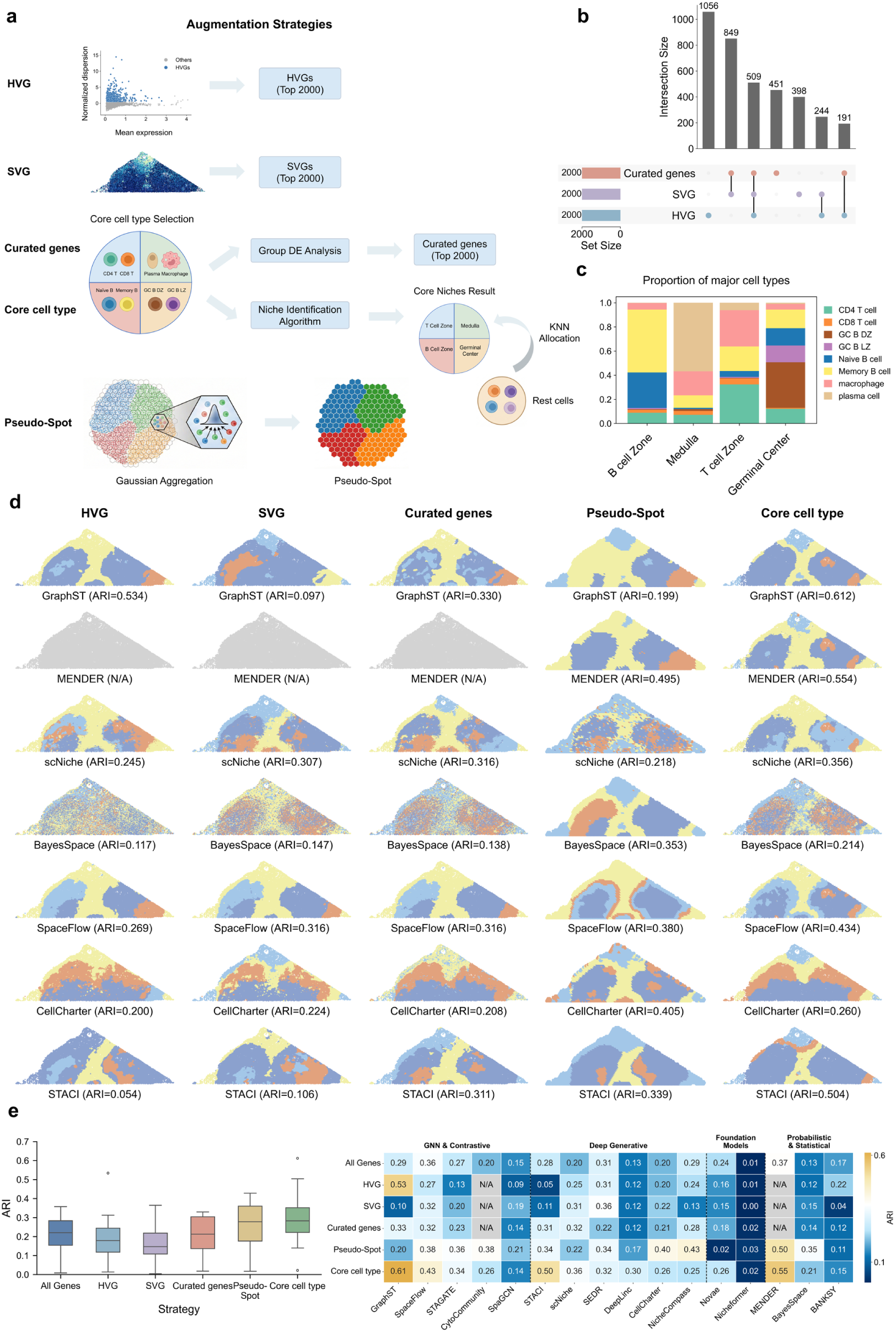
Feature selection, pseudo-spot aggregation, and core-cell-type-guided refinement modulate niche identification on the manually annotated human lymph node dataset. (a) Schematic overview of five augmentation strategies: HVG selection, SVG selection, curated-gene construction via group-wise DE on biologically defined core cell types, core-cell-type-guided refinement (running methods on core populations followed by KNN-based label propagation to remaining cells), and pseudo-spot aggregation via local Gaussian aggregation. (b) UpSet plot summarizing overlap among the 2,000-gene sets derived from HVG, SVG, and curated genes. (c) Proportions of major core cell types within each niche, normalized over the selected core cell-type set. (d) Representative spatial niche maps for seven methods (GraphST, MENDER, scNiche, BayesSpace, SpaceFlow, CellCharter, and STACI) under five augmentation strategies, with ARI appended at the bottom. (e) Summary of global and method-wise performance across strategies: boxplots show the ARI distribution over all benchmarked methods (left), and the heatmap reports method-specific ARI across strategies with methods grouped by algorithm family (right).

Fig. 5b summarizes the relationships between the three gene selection strategies: HVG and SVG each contained substantial strategy-specific components (1,056 HVG-only genes, 398 SVG-only genes); and the curated panel also retained a sizeable unique fraction (451 curated-only genes). This indicates that the three strategies emphasize partially distinct aspects of the expression landscape. The curated panel overlapped more strongly with SVG than with HVG: 1,358 genes were shared between curated genes and SVG (849 shared by curated and SVG only, plus 509 shared by all three sets), whereas only 700 genes overlapped between curated genes and HVG (191 shared by curated and HVG only, plus the same 509 shared core). Only 509 genes were common to all three panels, underscoring that a large fraction of HVG-driven variability is not captured by niche-focused features. Together, these patterns support a straightforward interpretation: HVG and SVG capture broad variability or spatial structure, but part of that signal can be orthogonal to niche-defining programs, whereas the curated panel is explicitly constructed to concentrate on compartment-discriminative transcriptional differences, making it a closer match to niche identification objectives.

In parallel, to investigate if spatial resolution, a primary differentiator among ST platforms, modulates analytical outcome, we evaluated whether reducing effective resolution via pseudo-spot aggregation can amplify niche-level signals. We mapped single-cell profiles onto a Visium-like hexagonal lattice and aggregated expression using Gaussian distance-weighted assignment, producing pseudo-spots that emulate spatial mixing and blurred boundaries (Fig. 5a). This procedure reduced the number of observations from 19,718 cells to 2,637 pseudo-spots, corresponding to an average of 7.48 cells per pseudo-spot, and yielded a multicellular spot-level representation with corresponding pseudo-spot reference labels while preserving the overall tissue geometry and all four niches. By attenuating cell-level idiosyncrasies and emphasizing locally averaged expression programs, pseudo-spot aggregation can enhance niche-level contrast and stabilize spatial patterns, yielding niche assignments that better reflect locally mixed microenvironments and blurred cellular boundaries.

Across strategies, several methods showed marked improvements in ARI relative to the default all-gene, single-cell resolution baseline, corresponding to the “All genes” setting in Fig. 5e, and in multiple cases recovered a niche layout that closely matched the ground truth. Complete method-level spatial assignments, latent embeddings, cell-type compositions, and multi-criteria results for the HVG, SVG, curated-gene, pseudo-spot, and core-cell-type settings are provided in Supplementary Figs. S2–S6, respectively. For HVG and SVG selections, across the six methods shown in Fig. 5d (scNiche, GraphST, BayesSpace, SpaceFlow, CellCharter, and STACI), the impact of input choice was strongly method dependent. Under HVG, GraphST achieved a substantially improved spatial partition (ARI 0.534). Under SVG, SpaceFlow showed a clear gain (ARI 0.316) and scNiche improved to ARI 0.307, whereas several methods remained weak (e.g., GraphST 0.097; STACI 0.106).

Relative to HVG/SVG, curated genes improves the segmentation to be more aligned with the annotation ground truth for multiple methods, with scNiche (ARI 0.316), SpaceFlow (ARI 0.316), and STACI (ARI 0.311) showing robust improvements and clearer separation of the major compartments. This pattern is consistent with the curated panel concentrating on compartment-discriminative programs learned from dominant cell types through niche-versus-rest contrasts, thereby reducing feature capacity spent on within-niche variation. In parallel, pseudo-spot aggregation provided a complementary route to improvement by smoothing cell-level stochasticity and amplifying locally coherent signals. Among the representative methods shown in Fig. 5d, MENDER achieved the highest agreement under the revised pseudo-spot setting (ARI 0.495), followed by CellCharter (0.405), SpaceFlow (0.380), BayesSpace (0.353), and STACI (0.339). scNiche improved only slightly to 0.218, whereas GraphST decreased to 0.199. These results indicate that coarser local aggregation can improve compartment-scale recovery for methods that effectively exploit smoothed neighborhood structure, but its effect remains strongly method dependent.

Core cell type refinement yielded the strongest and most coherent gains overall by anchoring niche inference to dominant populations and then propagating assignments to all remaining cells. This strategy produced the highest single-method agreement in Fig. 5e, led by GraphST (ARI 0.612), followed by MENDER (0.554) and STACI (0.504); scNiche (0.356) and SpaceFlow (0.434) also improved relative to their all-gene baselines. Under this setting, GraphST recovered an overall partition that closely recapitulated the ground-truth compartment layout. Similarly, it helps MENDER deriving a spatial partition that closely resembles the annotation ground truth. Moreover, GraphST, scNiche, and SpaceFlow more clearly delineated a compact, island-like Germinal Center embedded within the B cell zone, a structure that is difficult to resolve under All genes/HVG/SVG.

Quantitative assessment of each individual extrinsic guidance strategy is summarized in Fig. 5e. Relative to the all-genes baseline (median ARI 0.220), pseudo-spot aggregation increased the median to 0.340, and core cell type refinement raised it to 0.280, whereas curated genes yielded a similar median (0.215) but selectively improved a subset of methods. The heatmap makes these method-specific shifts explicit: curated genes increased agreement for scNiche (0.200 *→* 0.320) and STACI (0.280 *→* 0.310), while pseudo-spot aggregation produced large gains for BayesSpace (0.130 *→* 0.353), CellCharter (0.200 *→* 0.405), STACI (0.280 *→* 0.339), MENDER (0.370 *→* 0.495), and scNiche (0.200 *→* 0.218). Core cell type refinement delivered the strongest improvements for methods that can leverage a robust spatial scaffold, exemplified by GraphST (0.290 *→* 0.610), STACI (0.280 *→* 0.500) and scNiche (0.200 *→* 0.360).

Taken together, these results point to three complementary levers for improving niche identifiability in complex microenvironmental tissues. Curated genes sharpens the representation by prioritizing transcriptional programs that best separate niches, which stabilizes inference when expression differences are subtle and confounded by within-niche variation. Pseudo-spot aggregation increases signal-to-noise through local averaging, effectively suppressing cell-level stochasticity and emphasizing region-scale structure, which benefits methods whose objective is sensitive to noisy neighborhoods or fragmented boundaries. Core cell type refinement adds a biologically grounded prior by anchoring niche discovery to dominant populations that define canonical immune compartments, thereby enforcing plausible cell type compositions and spatial co-localization patterns. By fixing this core scaffold and propagating labels to remaining cells in a geometry-aware manner, the strategy both improves recovery of major compartments and increases sensitivity to smaller, embedded niches that arise from selective enrichment of specialized immune subsets, consistent with the concept of microenvironmental aggregation in immune tissues. It is worth noting that, however, response to various extrinsic augmentation strategies are method-specific and there is no one-fit-all solution. While strategic lineage weighing is perhaps the most encompassing augmentation, CellCharter responds best to pseudo-spot augmentation. GraphST, on the other hand, did not improve under the coarser pseudo-spot representation. Our results provide a comprehensive and algorithm-specific road map to analytic enhancement for niche identification.

### 2.6 Benchmark against distributed isolated niches and gradient spatial niches

A key distinction between spatial domains and tissue niches is that while spatial domain emphasizes on spatial continuity, niche prioritizes recurring spatial affinity between specific cell populations which does not necessarily connect into a large continuum. Therefore, the same type of niche could be sporadic and wide-spread in a tissue slice as isolated islands. In the annotated human lymph node data, Germinal Centers (GCs) is a representative of such detached niche. We therefore extracted a B-cell-zone subregion containing two spatially separated GC regions to define an island-niche benchmark (Fig. 6a). In our whole-tissue benchmark, GC zones are among the hardest niches to recover (Supplementary Fig. S9). The poor success rate of identifying GC zones is disappointing given their central role in adaptive immune response. GC is the major microanatomical site where B-cell activation, somatic hypermutation, affinity maturation, and class-switch recombination converge. At the core of GC, rapidly proliferating centroblasts undergo clonal expansion, surrounded by differentiating progeny, follicular dendritic cells (FDCs) that retain antigen for affinity-based selection, and macrophages that clear apoptotic debris from failed selection events. This architectural arrangement generates a discontinuous, polarized microenvironment with compact dimensions and sharp demarcation from surrounding tissue. Geometrically, GCs zones manifest as irregularly sized, spatially sparse islet embedded within the larger B cell zone. This nested, non-uniform organization violates the assumptions of many domain segmentation algorithms which favor globally smooth partitions or allocate clusters to the dominant sources of variance. Consequently, small GC islands are inclined to be absorbed into the surrounding follicular background or to fragmented into separate patches.

**Fig. 6.**
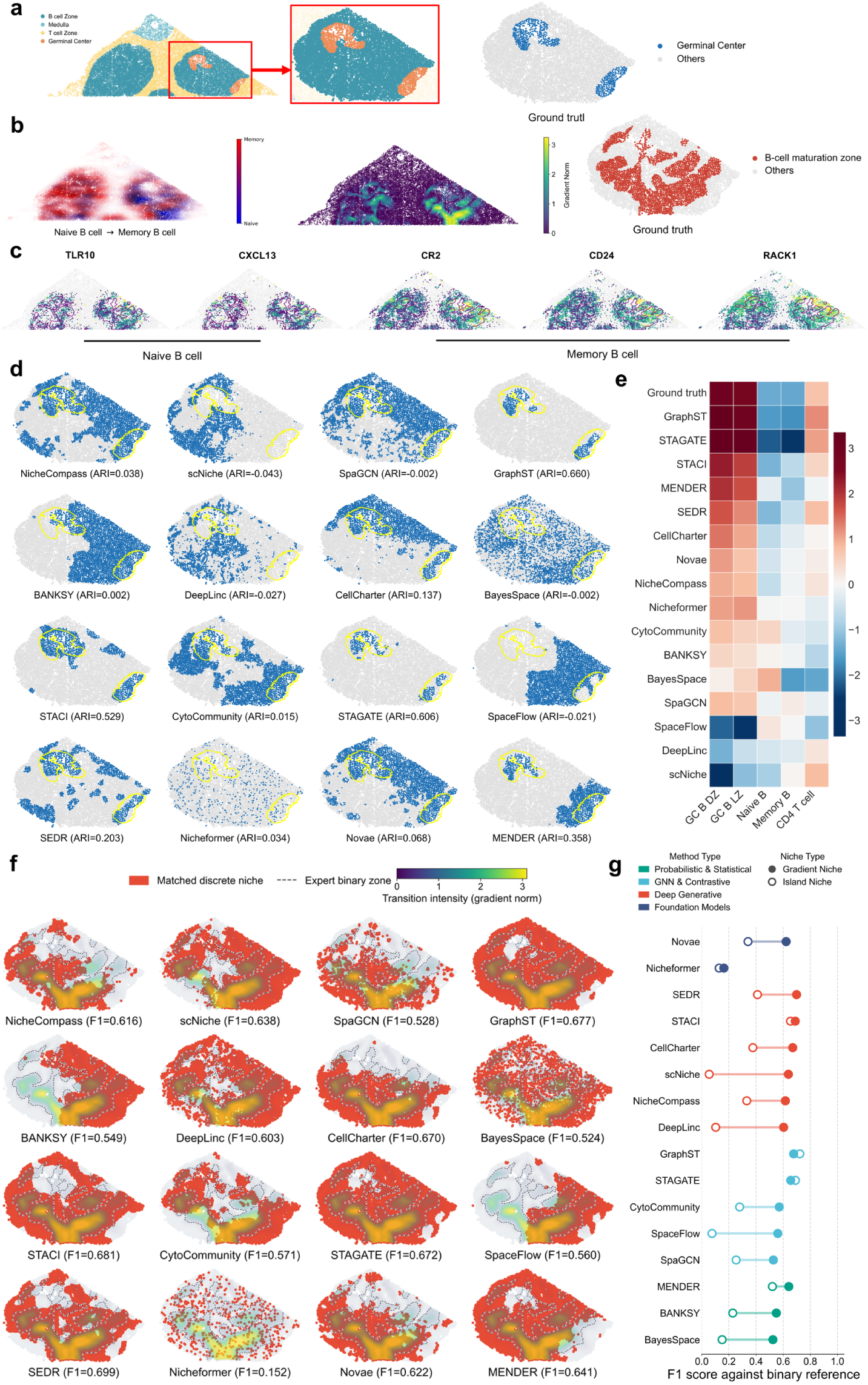
Benchmarking methods on island and transition niches in a selected subregion from the human lymph node dataset. (a) Definition of an island niche by extracting a B cell zone subregion containing two Germinal Center (GC) regions (left, full view; middle, zoom-in), and the binary GC ground truth (right). (b) Gaussian KDE-based normalized density fields for Naïve B and Memory B cells (left), the resulting gradient-norm map capturing local Naïve-to-Memory transition intensity (middle), and the binary B-cell maturation zone ground truth (right). (c) Expression of representative marker genes within the B-cell maturation zone: TLR10 and CXCL13 (Naïve B-associated), CR2, CD24, and RACK1 (Memory B-associated). (d) Island-niche identification results: for each method, the predicted niche with maximal overlap to the GC ground truth is shown (blue), with GC regions outlined in yellow. (e) Cell-type enrichment heatmap for the GC-matched niche from each method. (f) Transition-niche identification results: for each method, the predicted domain with maximal overlap with the expert-defined binary B-cell maturation-zone reference is shown in red over the continuous transition-intensity field; dashed outlines indicate the binary reference, and the corresponding F1 score is reported. (g) Summary of niche-type-specific performance using F1 score against the binary reference: filled circles denote the B-cell maturation zone (transition niche), and open circles denote the GC (island niche); methods are colored by algorithm family.

Beyond the identification of discrete island niches, the enforcement of non-overlapping spatial partitions risks overlooking critical transition zones. In these regions, cells often exhibit high plasticity and undergo functional reprogramming, manifesting as a spatial continuum rather than a set of demarcated domains. Unlike microenvironments in homeostatic stasis, these transition zones are highly dynamic; consequently, they present a fundamental challenge to the hard boundary logic of current algorithms, as the biological transitions are often too gradual to be resolved by discrete segmentation. The B-cell maturation zone in follicle is a typical transition niche, encapsulating continuous transitions from naïve B to memory B cell states. This transition reflects the canonical B cell maturation trajectory: naïve B cells residing in the mantle zone first migrate toward the T-B border, where they receive cognate antigen presentation and T cell assistance. Following activation, a subset of B cell migrates centrally to become GC B cells, undergoing iterative rounds of proliferation, somatic hypermutation, and affinity-based selection. Upon completing the activation program, cells differentiate into memory B cells and exit the germinal center. This migratory and differentiation path generates a complex, folded gradient zone, where the spatial coordinates of a cell directly correlate with its differentiation state. We annotated B-cell maturation zone by integrating the spatial distributions of naïve and memory B cells with a transition-intensity map that highlights where the naïve→memory shift is most pronounced (Fig. 6b).

Characterization of the naïve-to-memory transition zone facilitated the extraction of lineage-specific markers governing B-cell activation and maturation. By isolating the top-ranking genes that specifically co-occur within this spatial interface across both naïve and memory populations (Fig. 6c), we identified the core transcriptomic signatures orchestrating the B-cell maturation program. Within the transition zone, genes elevated in naïve B cells include *TLR10* (a Toll-like receptor family member), which is frequently linked to intrinsic dampening of B-cell immune signaling and is consistent with a locally constrained activation program [50], and *CXCL13* (a canonical follicular chemokine), reflecting preserved follicle-associated spatial organization cues [51]. In contrast, genes elevated in memory B cells include *CR2* (complement receptor 2), a BCR co-receptor that lowers activation thresholds and amplifies antigen-linked signaling, aligning with enhanced responsiveness of memory-leaning states [52]; *CD24*, associated with shifts in B-cell differentiation/activation states and suggesting incomplete stabilization within the transition zone [53]; and *RACK1* (Receptor for Activated C Kinase 1), a signaling scaffold that supports pathway integration and signal propagation, consistent with heightened signaling competence [54]. Additional naïve- and memory-B-cell-associated spatial markers supporting this transition are shown in Supplementary Fig. S8.

At the full ROI scale, neither the discrete, island-like germinal centers (GCs) nor the gradient-like maturation niches were robustly recovered (Fig. 4). To investigate whether attenuating out-of-follicle confounding signals could enhance the contrast between the GCs, B-cell transition niches, and the background, we isolated the B follicles and re-evaluated the performance of all 16 algorithms within this spatially constrained context. The results are summarized in Fig. 6d,e. For the island niche (GC) task, after attending to the follicle region, a small subset of methods now are capable of detecting GC regions. GraphST (ARI 0.660), STAGATE (ARI 0.606), and STACI (ARI 0.529) most consistently recovered compact, spatially coherent GC islands, while MENDER (ARI 0.358) and SEDR (ARI 0.203) captured GC-associated areas with weaker boundary precision. CellCharter (ARI 0.137) yielded broadly plausible GC contours but with limited overall agreement.

We identified the domains most proximal to the germinal centers (GCs) for each method, highlighted in blue in Fig. 6d, and performed a differential enrichment analysis to characterize the cellular composition both within and outside these segmented boundaries. This analysis quantifies the fold-change in relative cell-type abundance within the segmented domain compared to the remaining ROI, providing a measure of the algorithm’s ability to isolate specific cellular niches. Consistent with the ARI scores, GraphST, STAGATE and STACI yield cell-type enrichment fold changes that most akin to the annotation ground truth. To ensure consistency, we repeated the same evaluation on a separate follicle (Supplementary Fig. S10). Once again, we observe GraphST, STAGATE, CellCharter, MENDER and STACI providing segmentation results that are most similar to the GC annotation ground truth.

In contrast, the identification of transition zones presents a more challenging landscape for current algorithms. The GC and B-cell maturation niches are not mutually exclusive, exhibiting a non-trivial spatial overlap that is biologically essential but computationally difficult to resolve. Because most domain segmentation frameworks operate via discrete partitioning, they are inherently unable to accommodate these overlapping architectures. As illustrated in Fig. 6f, the best-matched discrete domains frequently covered substantial portions of the thresholded B-cell maturation-zone reference but extended beyond the strongest transition-intensity ridges or captured only restricted parts of the continuous field. Several methods therefore achieved comparatively high F1 scores against the binary reference, although none reproduced the complete graded and locally heterogeneous structure of the underlying naïve-to-memory transition (Fig. 6g). This distinction indicates that binary-region agreement reflects the approximate spatial localization of the transition zone, but does not by itself demonstrate recovery of the continuous maturation gradient.

Across the two niche geometries summarized in Fig. 6g, GraphST, STAGATE, STACI, and MENDER showed the most balanced performance across both the island-like and transition-like tasks. This performance pattern is consistent with their respective inductive biases: the top methods either learn spatially regularized representations on a neighbor graph (GraphST/STAGATE/STACI), which helps preserve compact islands and maintain local continuity, or explicitly encode neighborhood context statistics (MENDER), which stabilizes signals that are inherently microenvironmental. While the improved performance on isolated B follicles is encouraging, it also highlights the fragility of the analytical framework and its dependence on the tissue context. Overall, these results underscore that tissue niches are inherently different from transcriptome domains: performance on well-bounded compartments does not necessarily translate to biology-centric structures that are nested (islands) or continuous (transitions), and resolving such niches requires models that can retain fine local structure without collapsing it into dominant tissue-scale variation.

### 2.7 Assessing the impact of peripheral cell types through simulation

To confirm and quantify the impact of peripheral cell types in niche identification, we conducted a suit of simulations that gradually reduce the contrast between originally distinct artificial domains. The initialization of the simulation setup is summarized in Fig. 7a. In this simulation, we generated a single-cell resolution de novo lymph node spatial transcriptomics dataset using SRTsim [55], guided by the annotated human lymph node reference. We specified 4 high contrast niches, imitating the four non-overlapping niches in the lymph node: medulla, T cell zone, B cell zone, and germinal center. We set the cell type compositions in each artificial niche to closely follow their counterparts in the real data, while deliberately increasing the percentage of the marker lineages: plasma cell and macrophage for medulla; CD4 and CD8 T cells for T zone; naïve and memory B cells for B zone; GC B cells for germinal center [34]. The initial simulation represents the most ideal case for domain segmentation, where transcriptomic landscapes are maximally different between niches, generating clear-cut boundaries in between zones.

**Fig. 7.**
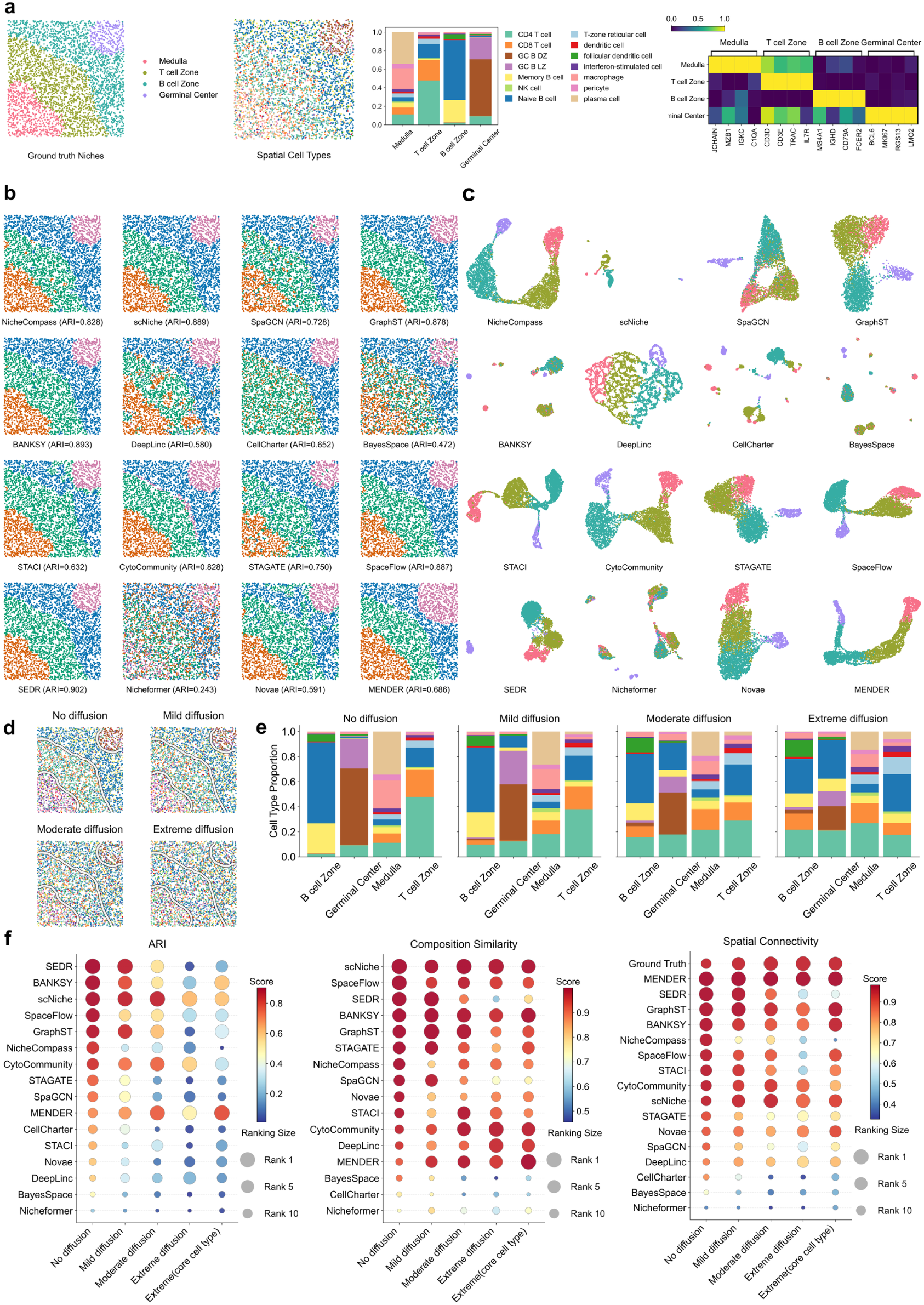
Benchmarking results of niche identification methods on simulated lymph node data across cell-type compositional diffusion settings. (a) Ground-truth niches in the simulated tissue, the corresponding spatial distribution of cell types, niche-wise cell-type composition, and representative marker-gene expression patterns. (b) Spatial niche assignments produced by 16 methods, with ARI scores attached. (c) UMAP visualizations of method-specific latent embeddings colored by ground-truth niche labels. (d) Illustrative cell-type spatial maps under four diffusion settings (no, mild, moderate, and extreme diffusion), showing progressive dispersal of dominant cell types. (e) Niche-wise cell-type composition profiles across diffusion settings. (f) Bubble matrix summarizing ARI, cell-type cosine similarity, and spatial connectivity across diffusion settings and an additional condition applying the core-cell-type refinement strategy under extreme diffusion; bubble color encodes metric value and bubble size indicates rank, with methods ordered by performance in the no-diffusion setting.

Domain segmentation over the initial simulation produces high quality domains for the majority of the 16 algorithms. Their domain segmentation results are summarized in Fig. 7b, c. Most of the algorithms, with CellCharter and Nicheformer slightly struggling, are capable of delineating the 4 domains in correct topological order, in spite of slight over-extensions by some of the domains.

To assess algorithmic robustness, we systematically introduced peripheral cell interference by progressively down-weighting the frequency of niche-specific marker lineages. These vacancies were filled by a combination of marker lineages from adjacent niches and a constant background composition encompassing all cell types. The resulting spatial landscapes following this progressive diffusion are illustrated in Fig. 7d-e. As diffusion increases, marker lineage dominance within each niche diminishes, leading to a convergence toward a shared global background composition. Notably, even at maximum diffusion, evident compositional divergence persists between niches, albeit with a markedly attenuated effect size.

This controlled mixing primarily weakened niche separability by compressing molecular differences between neighboring environments(Supplementary Fig. S18). As diffusion increased, the mean between-niche Euclidean distance among Gaussian-smoothed key-lineage profiles decreased from 14.8 to 11.4, whereas the within-niche distance remained nearly stable at 8.9–9.2, reducing the inter-to-intra niche separability ratio from 1.65 to 1.24 (Supplementary Fig. S18a). Consistently, representative marker programs of the Medulla, T-cell zone, B-cell zone, and germinal center became progressively less spatially confined and less distinct from the surrounding tissue (Supplementary Fig. S18b). This molecular dilution was accompanied by increasing cellular mixing: representative non-core populations rose from 10.1% to 20.4% of cells and became increasingly interspersed across niche boundaries (Supplementary Fig. S18c). Together, these results show that peripheral-cell diffusion progressively erodes the molecular contrast available for distinguishing neighboring niches, providing a mechanistic basis for the decline in domain-segmentation performance.

The performance benchmarks across the simulated diffusion gradient are summarized in Fig. 7f. Complementary Macro-F1 and silhouette-score results across the same diffusion settings are provided in Supplementary Fig. S11. We observed a consistent degradation in segmentation accuracy as diffusion intensified across all evaluated methods. Notably, scNiche, CytoCommunity, and MENDER exhibited superior robustness, maintaining high Adjusted Rand Index (ARI) scores even under extreme diffusion scenarios (scNiche: 0.89 to 0.62; CytoCommunity: 0.83 to 0.57; MENDER: 0.69 to 0.49). In contrast, many methods exhibited a shared sensitivity pattern in which ARI dropped sharply as diffusion increased. The resilience of CytoCommunity and MENDER suggests that explicitly modeling the local microenvironment, rather than relying solely on the spatial transcriptome, is a viable strategy for mitigating peripheral cell interference.

Furthermore, restricting inference to core cell types under extreme diffusion (Fig. 7f) yielded significant performance gains for BANKSY, GraphST, and STACI, indicating that a cell-type-anchored structural prior can partially restore niche separability. While biological consistency remained high for scNiche and BANKSY, spatial connectivity generally weakened as a function of diffusion, reflecting the transition toward more fragmented or stochastic niche assignments.

The diffusion-induced loss of niche contrast has a direct biological analogue in lymph-node tissue. Lymph-node compartments are organized by chemokine gradients and stromal scaffolds but remain permeable to continuous immune-cell trafficking, such that niche-defining populations coexist locally with accessory or transient populations from adjacent regions. This cellular mixing increases within-niche compositional heterogeneity and reduces the contrast between neighboring microenvironments, providing a biological basis for the progressive performance degradation observed in the simulation. To examine how closely this controlled perturbation reflects the real tissue, we compared the within-dataset ARI ranks of all 16 methods rather than their absolute ARI values, because the real and simulated datasets differ in cell number, gene dimensionality, spatial geometry, and overall task complexity (Supplementary Fig. S15). Spearman rank correlations with the real lymph-node benchmark were 0.52, 0.43, 0.59, and 0.26 under no, mild, moderate, and extreme diffusion, respectively, with the highest concordance occurring under moderate diffusion. This partial and non-monotonic agreement indicates that compositional mixing captures an important component of the difficulty encountered in the real lymph node, while additional features of the tissue, including irregular and nested niche geometries, continuous cell-state variation, and single-cell expression heterogeneity, also contribute substantially to method performance.

Collectively, these results prove that peripheral cell type interference, thereby reducing cell type composition contrasts between adjacent niches, is indeed a major challenge for niche identification. While extrinsic augmentations, such as anchoring the analysis to knowledge-guided core lineages, offer a viable interim solution, future niche identification algorithms should incorporate adaptive, multi-scale weighting mechanisms that can autonomously distinguish between functionally conserved core constituents and transient peripheral populations.

### 2.8 Cross comparison to anatomically compartmentalized tissues across spatial technologies

Unlike tissue compartments defined by coarse anatomical boundaries, tissue niches of the lymph node are jointly governed by compositional mixing, transient cellular states, and context-dependent signaling gradients. Previous domain segmentation benchmarks have largely focused on concordance with anatomy-driven annotations, such as those derived from cortical layers or specific brain regions. These datasets typically feature large, spatially contiguous compartments characterized by high transcriptional contrast and strong spatial autocorrelation. While informative, these anatomy-centric benchmarks differ fundamentally from the complex microenvironments in our human lymph node reference, where lineage specialization lacks discrete physical boundaries. To evaluate and cross-compare how segmentation algorithms generalize across these distinct architectural paradigms and measurement modalities, we extended our assessment to two widely utilized anatomical-domain benchmarks: the 10x Visium human dorsolateral prefrontal cortex (DLPFC) [31] slice 151673 (spot-based, low resolution) and the Stereo-seq MOSTA [32] E16.5 mouse brain dataset (single-cell resolution) (Fig. 8a-e).

**Fig. 8.**
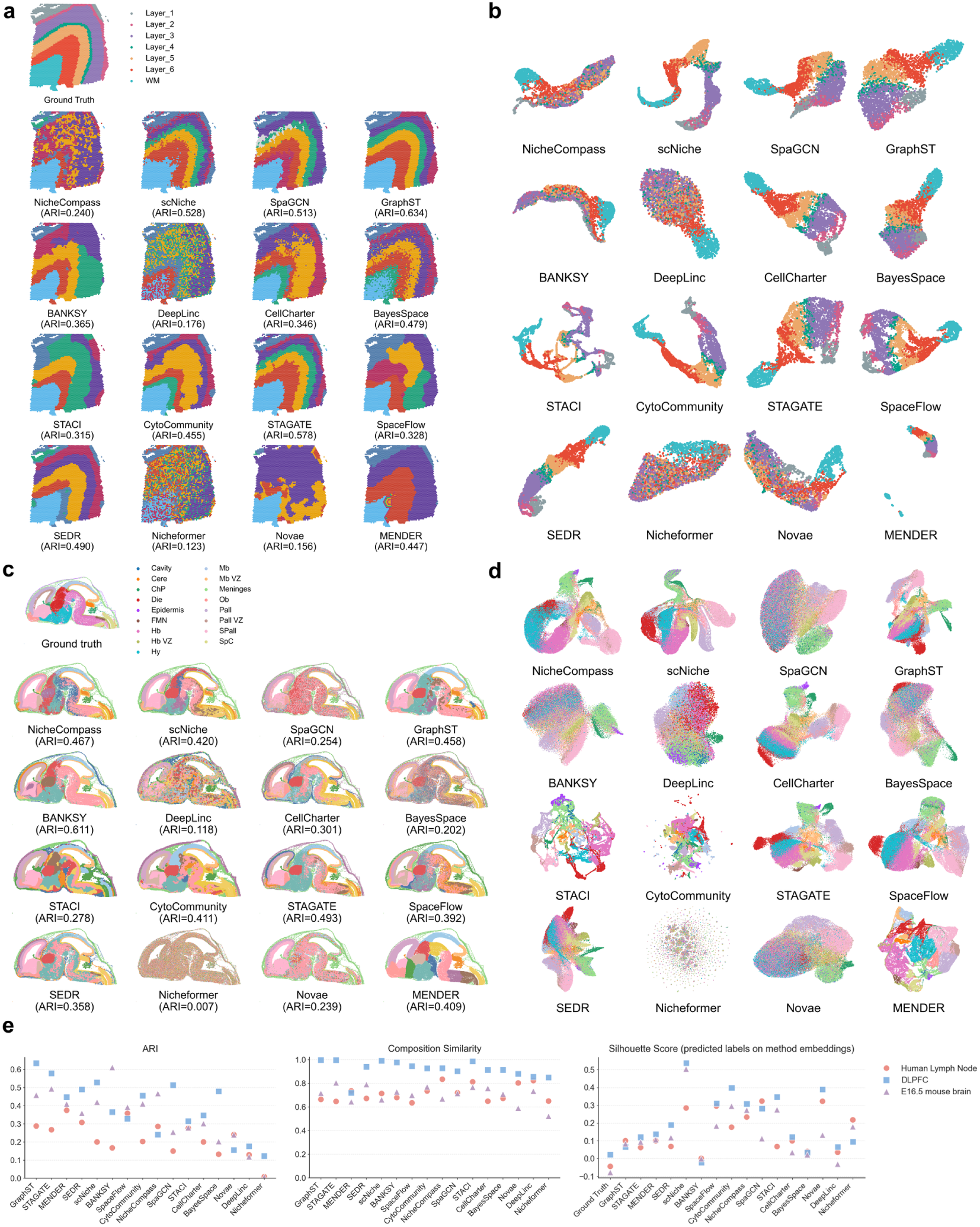
Benchmarking results on the DLPFC slice 151673 and the MOSTA E16.5 mouse brain dataset. (a) Spatial distributions of ground-truth anatomical domains in DLPFC and method-predicted niches, with ARI scores attached. (b) UMAP visualizations of method-specific latent embeddings for DLPFC, colored by the corresponding ground-truth domain labels. (c) Spatial distributions of ground-truth anatomical regions in the E16.5 mouse brain and method-predicted niches, with corresponding ARI scores at the bottom. (d) UMAP visualizations of method-specific latent embeddings for the mouse brain dataset, colored by ground-truth region labels. (e) Cross-dataset summary of ARI, cell-type cosine similarity, and silhouette score across the manually annotated human lymph node reference, DLPFC, and E16.5 mouse brain. Silhouette scores are computed on each method’s latent embedding using that method’s predicted niche labels, whereas the ground-truth baseline is computed on a transcriptome-derived UMAP using ground-truth labels; dataset markers/colors facilitate comparison across tissue contexts and technologies.

The DLPFC dataset represents a prototypical low-resolution setting in which cortical layers form well-separated anatomical domains. We treated layer annotations as niches and evaluated all methods under a harmonized pipeline. For methods requiring cell-type composition features, we followed the scNiche [24] workflow and inferred spot-level cell-type proportions using Cell2location-based deconvolution [56], providing consistent cell-type composition inputs without altering the underlying spatial domain ground truth. Across methods, DLPFC layers were recovered with comparatively high agreement (Fig. 8a). GraphST achieved the strongest concordance (ARI 0.634), followed by STAGATE (ARI 0.578), scNiche (ARI 0.528), and SpaGCN (ARI 0.513); several additional methods also performed competitively, including SEDR (ARI 0.490), BayesSpace (ARI 0.479), CytoCommunity (ARI 0.455), and MENDER (ARI 0.447). This performance profile is consistent with the underlying task structure: layered cortex favors methods that couple representation learning with spatial smoothness and that prioritize large-scale, contiguous partitions. By contrast, approaches that produced more fragmented assignments or learned embeddings misaligned with discrete layers showed reduced agreement (e.g., DeepLinc, ARI 0.176; Nicheformer, ARI 0.123), which was also reflected by entangled latent spaces when UMAPs were colored by layer labels (Fig. 8b). We next evaluated MOSTA E16.5 mouse brain, a higher-complexity, single-cell-resolution Stereo-seq benchmark with a larger number of annotated regions and more intricate tissue geometry (treated here as anatomical-domain niches). Relative to DLPFC, the increased number of regions, finer spatial granularity, and broader cellular diversity reduced performance for many methods (Fig. 8c), underscoring the difficulty of resolving numerous boundaries in a dense single-cell setting. Nevertheless, several approaches still reconstructed major region structure with good fidelity, led by BANKSY (ARI 0.611), followed by STAGATE (ARI 0.493), GraphST (ARI 0.458), NicheCompass (ARI 0.467), and scNiche (ARI 0.420); CytoCommunity (ARI 0.411) and MENDER (ARI 0.409) also remained competitive. Qualitatively, these better-performing methods tended to preserve contiguous anatomical territories while accommodating region-specific subdivision (Fig. 8c). In contrast, DeepLinc (ARI 0.118) and Nicheformer (ARI 0.007) exhibited limited agreement and produced latent embeddings that remained visibly mixed with respect to region labels across both anatomical benchmarks (Fig. 8b,d), suggesting that the dominant variation captured by these representations is less aligned with the anatomical annotations.

Finally, a comparative analysis of intra- and inter-domain variance offers a mechanistic explanation for the performance discrepancies observed across datasets. The cross-dataset comparison of ARI, cell-type cosine similarity, and silhouette score further summarizes these dataset-dependent performance patterns (Fig. 8e). Using Silhouette scores to evaluate ground-truth annotations, we found that both the E16.5 mouse brain and the reactive lymph node datasets registered near-zero or negative values within their transcriptomic embeddings. This suggests that for these complex tissues, intra-domain transcriptomic homogeneity does not significantly exceed inter-domain heterogeneity, a finding that reflects the profound cellular complexity and compositional variance inherent to each domain. Notably, while many algorithms generated substantially higher silhouette scores when evaluated on their own latent embeddings using predicted labels, these scores remained consistently low when mapped back to the original transcriptomic space (Supplementary Fig. S12). This result highlights a critical divergence from a foundational premise of most domain segmentation algorithms: the assumption that spatial domains are defined by cohesive transcriptomic profiles that are distinct from those of neighboring regions. Our results indicate that in biologically complex environments, this assumption is frequently violated, suggesting that successful segmentation requires models that can account for high local variance rather than just global similarity.

Across these settings, spatial resolution changes the unit over which expression and cellular composition are averaged. In Visium, each spot integrates multiple cells, reducing cell-level stochasticity and favouring methods that enforce spatial smoothness over broad, contiguous structures. GraphST and STAGATE therefore provide suitable starting points for layered anatomical data, with BayesSpace offering a spot-oriented statistical alternative and scNiche being useful when reliable deconvolution is available. Cell-resolved CosMx preserves rare populations and fine boundaries but also exposes strong lineage effects and local compositional heterogeneity. In this setting, MENDER performed most strongly when cell-type labels were available, while NicheCompass and STACI remained competitive alternatives. The matched-tissue pseudo-spot experiment reinforces this interpretation: local aggregation increased the median ARI from 0.220 to 0.340, although the gains remained method specific. Stereo-seq combines single-cell granularity with dense sampling and numerous anatomical boundaries; BANKSY, STAGATE, GraphST, and NicheCompass performed most consistently on the E16.5 mouse brain. These recommendations should be regarded as dataset-informed starting points rather than universal rankings, and for atlas-scale single-cell data should be considered together with the scalability limits summarized in Fig. 9.

**Fig. 9.**
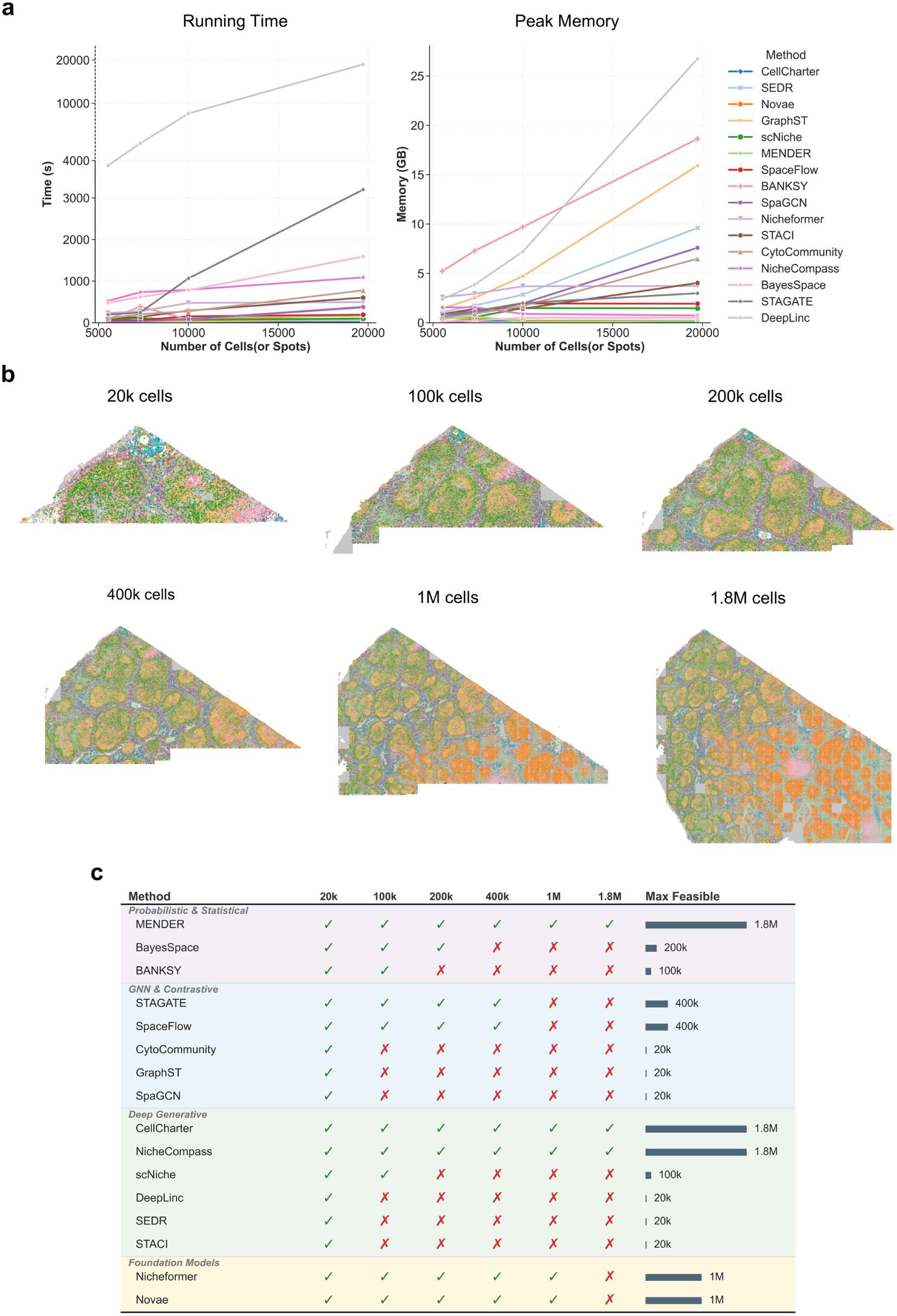
Benchmarking computational efficiency and scalability of niche identification methods across increasing data sizes. (a) Wall-clock runtime (left) and peak memory usage (right) measured when running each method on four progressively larger subsets of the human lymph node dataset (5,521; 7,319; 9,999; and 19,718 cells). (b) Spatial maps of varying-size sub-sections (20k, 100k, 200k, 400k, 1M, and 1.8M cells) from the full CosMx human lymph node field of view, illustrating the increasing tissue coverage and complexity used for large-scale stress testing. (c) Feasibility matrix summarizing whether each method successfully completed under a fixed resource budget (128 GB system RAM and a single NVIDIA GeForce RTX 4090 GPU with 24 GB VRAM) across data sizes (✓, successful; ×, failed due to RAM/CPU memory or GPU VRAM limitations and related resource constraints), grouped by method family, together with the maximum dataset size each method could process successfully.

### 2.9 Benchmarking computational efficiency and scalability across varying data sizes

The ultra-large (1.85 million cells), single-cell-resolution lymph node CosMx dataset presents a unique opportunity to benchmark the computational efficiency, resource-friendliness and the scalability of domain segmentation algorithms. To provide a controlled comparison, we first benchmarked runtime and peak memory on four progressively larger subsets of the human lymph node dataset (5,521; 7,319; 9,999; and 19,718 cells or pseudo-spots), holding the gene panel fixed across all subsets to ensure that scaling trends primarily reflect growth in the number of observations rather than differences in feature dimensionality (Fig. 9a). As expected, both wall-clock time and peak memory increased with dataset size for nearly all methods, but the rate of increase was highly method dependent. In particular, DeepLinc and STAGATE showed the steepest runtime growth, becoming noticeably slower at higher cell counts, while DeepLinc, BANKSY, GraphST, and SEDR exhibited among the highest peak memory footprints, indicating poorer practical scalability under these settings (Fig. 9a). In contrast, MENDER, CellCharter, NicheCompass, and scNiche displayed comparatively stable growth curves over this size range, suggesting more favorable scaling behavior in both time and memory.

To probe scalability beyond 2 *×* 10^4^ cells and better approximate modern high-throughput SRT workloads, we further constructed large-scale stress-test inputs by uniformly sampling subdatasets of 20k, 100k, 200k, 400k, 1M, and 1.8M cells from the full CosMx human lymph node field of view (Fig. 9b). These subdatasets progressively expand tissue coverage and morphological complexity, thereby increasing both the number of spatial neighbors that must be modeled and the overall heterogeneity that algorithms must accommodate. All large-scale runs were performed under a fixed computational budget (128 GB RAM and a single NVIDIA GeForce RTX 4090 GPU with 24 GB VRAM) to ensure comparability of feasibility outcomes. Under this standardized, resource-constrained setting, scalability limits diverged sharply across methods (Fig. 9c). Only three methods (MENDER, NicheCompass, and CellCharter) remained fully scalable to 1.8M cells, whereas Nicheformer and Novae were feasible up to 1M cells. At intermediate scales, STAGATE and SpaceFlow completed up to 400k cells, BayesSpace up to 200k, and BANKSY and scNiche up to 100k. In contrast, several methods, including CytoCommunity, GraphST, SpaGCN, DeepLinc, SEDR, and STACI, failed once the dataset size increased beyond 20k, typically due to RAM/CPU memory or GPU VRAM constraints and associated downstream failures (e.g., graph construction, neighborhood aggregation, or model training instability). Collectively, these results highlight a widening gap between the rapidly increasing scale of single-cell-resolution spatial platforms and the computational readiness of many current niche identification methods: while technological advances now routinely deliver 10^5^-10^6^ cells per experiment, most existing algorithms remain constrained by super-linear time/memory growth and resource-intensive optimization, underscoring the need for scalable algorithmic designs (e.g., linear-time neighborhood construction, minibatch/streaming training, and memory-aware implementations) to enable routine, accurate niche inference on million-cell spatial atlases.

## 3 Discussion

The advent of spatially resolved transcriptomics has provided an unprecedented lens into cellular biology, enabling us to move beyond simple cell typing to investigate how cells interact with their surrounding microenvironments, communicate with neighbors, and modulate their transcriptional programs in response to extrinsic stimuli. Historically, niche identification has been treated as synonymous with domain segmentation, a paradigm that assumes tissue architectures can be partitioned into discrete, non-overlapping domains characterized by relatively homogeneous cellular and transcriptomic profiles. In this work, we demonstrate that while key functional lineages often exhibit distinct spatial preferences when viewed in isolation, these boundaries become significantly attenuated when integrated into the full cell-type compositional landscape of the tissue. This reduced signal-to-noise ratio creates a fundamental conflict for existing domain segmentation frameworks; these algorithms are often forced to choose between competing objectives: isolating stochastic cell clusters, capturing collective paracrine responses, maintaining transcriptomic homogeneity, or identifying abrupt shifts in cellular composition. Consequently, few algorithms can natively deliver segmentation results that recapitulate expert-curated annotations, which are typically guided by the distinct, mutually exclusive distributions of core lineages informed by extrinsic domain knowledge.

A critical observation emerged from our benchmark is the existence of both horizontal (cross the entire tissue slice) and vertical (local spatial spots) discrepancies in spatial transcriptomic data. In the specific context of the human lymph node, we observed a clear, mutually exclusive spatial enrichment of key immunological lineages: including T cells, naïve and memory B cells, germinal center (GC) B cells, and plasma cells. Collectively, these distinct cellular territories span nearly the entire tissue area, forming a complex mosaic of functional zones. While these disjoint lineage distributions provide powerful structural cues for spatial partitioning, existing algorithms prioritize vertical transcriptomic variance at the individual spot level. At this granular scale, the signal-to-noise ratio is frequently compromised by the infiltration of peripheral cell lineages, leading to high compositional entropy that obscures niche boundaries. By focusing predominantly on localized heterogeneity, these methods largely fail to leverage the global architectural logic of lineage distribution, leading to a significant disconnect between computational output and biological structure.

Furthermore, our results suggest that the foundational assumption that transcriptomic profiles are inherently more similar within a domain than across domains, requires a more nuanced re-evaluation. The spatial transcriptomic signal is a convolution of two primary factors: the cell-type composition of a given locale and the specific transcriptional state of individual constituent cells. While we do observe niche-specific expression patterns for individual cell types, these localized shifts are often subtle and limited to a few genes and pathways. When compounded by the stochastic infiltration of peripheral cell types, the resulting intra-niche transcriptomic homogeneity is markedly attenuated, frequently falling below the threshold required for robust unsupervised segmentation. This dilution of inter-niche contrast is further exacerbated by the high resolution of the CosMx platform, where transcriptomic profiles are recorded at cellular resolution rather than being physically aggregated into spatial spots. Naturally, at single-cell resolution, the transcriptomic similarity between neighboring cells of disparate lineages cannot eclipse the inherent biological affinity between distal cells of the same type. Consequently, in the absence of deliberate spatial smoothing, via pseudo-spotting for instance, many algorithms that excelled at spot-level segmentation struggle to maintain performance at single-cell resolution. These observations call for a more sophisticated utilization of the transcriptomic homogeneity property in future niche identification frameworks. Specifically, transcriptomic variance should be modeled conditionally within specific cell types, while compositional similarity should be contextualized and limited to functional subsets of lineages rather than being treated as a globally inclusive metric.

Nonetheless, we observed that several methods, while exhibiting limitations in their default configurations, demonstrated significant performance gains through expert-guided refinements. Specifically, MENDER, STACI, and CellCharter showed a substantial increase in accuracy following the application of local transcriptomic smoothing, while GraphST benefited from the targeted use of highly variable genes (HVGs). Furthermore, GraphST, MENDER, and STACI achieved their peak performance when the analysis was constrained to core immunological lineages, a strategy facilitated by the integration of domain-specific prior knowledge. Intriguingly, the utilization of spatially variable genes (SVGs) or specific core lineage markers did not yield a uniform performance improvement across the benchmarked algorithms. This discrepancy raises critical questions regarding the efficacy of traditional gene selection strategies in the context of niche identification, suggesting that global transcriptomic features may occasionally outperform targeted marker sets in resolving complex microenvironments. Finally, we observed that the performance of spatial segmentation is highly context-dependent. While most algorithms were incapable of resolving the germinal center (GC) niche from the surrounding follicle at the full ROI scale, a targeted sub-sampling of the follicles enabled MENDER, GraphST, STAGATE, and STACI to successfully segment the GC zone. This suggests that the signal-to-noise ratio of functional niches is often obscured by distal background transcriptomes, and that localized spatial focus can restore algorithmic sensitivity.

It is important to emphasize that our findings do not invalidate the utility of current domain segmentation algorithms. Rather, we propose that domain segmentation and niche identification, while interrelated, represent fundamentally distinct analytical objectives. From our perspective, domain segmentation primarily targets the identification of discrete, macroscopic tissue compartments defined by structural homogeneity. In contrast, niche identification seeks to resolve dynamic, functional microenvironments characterized by complex cellular stoichiometry and plastic transcriptional states. The high performance of these algorithms in compartmentalized tissues, such as the DLPFC data, demonstrates that when tissue domains are mechanistically and spatially distinct, current frameworks are fully capable of generating high-quality, biologically concordant segmentations. Our work, however, underscores the necessity of distinguishing these structural assignments from functional niche discovery. We advocate for a strategic shift in how we approach spatially diffuse tissues, calling for the development of specialized methodologies tailored specifically to the unique challenges of niche identification.

Several limitations of the current study delineate clear directions for future research. First, establishing a consensus ground truth for spatial niches remains a formidable challenge; manual annotation is not only labor-intensive but inherently reflects expert-specific interpretative biases regarding spatial resolution and biological priority. Second, although our benchmark spans diverse niche geometries and technologies, each dataset is currently analyzed independently and the evaluation remains centered on spatial transcriptomics. Future benchmarks should therefore encompass both joint multi-slice modeling across conditions, technologies, and developmental stages [57] and complementary spatial modalities, including proteins, chromatin accessibility, and metabolites. These molecular layers may capture regulatory and functional organization that is only indirectly reflected in RNA abundance, while multi-task frameworks such as STAX provide a route toward modeling tissue architecture consistently across diverse spatial omics data types [58]. Third, the selection of an optimal algorithm is often dictated by application-specific trade-offs, such as prioritizing boundary precision over the resolution of continuous gradients, or favoring model interpretability over raw concordance. This motivates the continued development of evaluation frameworks that move beyond structural agreement to connect niche assignments with downstream functional validation. Consequently, we intend to expand our benchmarking suite to include increasingly complex niche architectures, providing a robust foundation for the next generation of spatial biology tools.

## 4 Conclusion

In this study, we established a comprehensive benchmarking suite for spatial niche identification, evaluating 16 state-of-the-art algorithms across diverse architectural paradigms. To move beyond the study of mechanistically compartmentalized tissues, we generated a high-resolution reference by thoroughly annotating a human reactive follicular hyperplasia (RFH) lymph node using CosMx SMI data. Our annotation revealed distinct, mutually exclusive spatial territories of key lineages, facilitating the demarcation of non-overlapping functional niches. Simultaneously, we identified a follicular B-cell maturation niche characterized by a transcriptomic continuum that partially overlaps with the germinal center (GC) architecture. Our analysis reveals a critical mismatch in spatial data: while the global distribution of key lineages provides clear and decisive architectural cues, these signals are substantially attenuated when resolved as local cell-type compositions. This discrepancy between macro-scale organization and spot-level heterogeneity represents a significant hurdle for existing domain segmentation frameworks; consequently, few algorithms in their default configurations can fully recover the biological boundaries of these niches. Despite these limitations, we demonstrate that expert-guided strategies, specifically the strategic weighting of core functional lineages, can rescue segmentation performance for select methods, most notably GraphST and MENDER. Finally, cross-platform comparisons against structured tissues like the DLPFC indicate that the foundational assumption of intra-domain transcriptomic homogeneity is frequently violated in dynamic, non-compartmentalized environments. These findings underscore the necessity for a new generation of niche-aware algorithms that prioritize lineage-specific structural priors over global transcriptomic similarity. Coupled with our expert-annotated lymph node reference and benchmarking suite, this work provides both the framework and the foundation for the development of future niche identification paradigms.

Both the lymph node annotation and the benchmark suite could be found at: https://github.com/WYXNICK/spatial-niche-benchmark

## 5 Methods

### 5.1 Construction and annotation of four major lymph node niches

A representative region of interest (ROI) comprising 19,718 cells was selected from a CosMx spatial transcriptomics dataset of human lymph node (total *∼*1.85 million cells), encompassing all major functional compartments (B cell zone, Germinal Center, T cell zone, and Medulla). While the original dataset provided broad cell type annotations, we resolved cellular heterogeneity at a finer scale to achieve more accurate niche delineation. Cell type identities were refined through unsupervised clustering, differential expression (DE) analysis, and expert curation, ultimately yielding 16 distinct cellular subtypes. These include functionally specialized subsets such as naïve and memory B cells, dark zone (DZ) and light zone (LZ) GC B cells, CD4+ and CD8+ T cells, and plasma cells.

To validate these annotations, we performed hierarchical DE analysis. First, lineage-level DE confirmed that broad cell lineages (B cells, T cells, Myeloid cells) exhibit distinct molecular profiles. Second, subtype-level DE within each lineage validated specific subtypes based on canonical marker genes (e.g., *IGHD* and *TCL1A* for naïve B cells; *TNFRSF13B* for memory B cells). This systematic validation ensures that each assigned label corresponds to biologically accurate cell identities.

To overcome the sparsity and positional noise inherent in single-cell spatial coordinates, we utilized continuous spatial density fields rather than discrete cell positions for robust geometric partitioning. Gaussian Kernel Density Estimation (KDE) was computed for lineage-defining cell populations, specifically B cells, GC B cells, T cells, and Plasma cells, using the SciPy library [59]. For each cell type, the continuous spatial density *D*(*x*) was calculated with a bandwidth factor of 0.1 to preserve local geometric features while smoothing stochastic dropout noise, and subsequently normalized to the range [0, 1]:

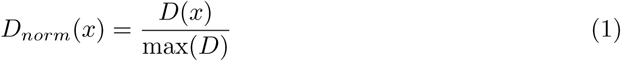

To guide accurate segmentation, structural boundaries were visualized by mapping the density co-occurrence of adjacent cell lineages (e.g., B cells vs. T cells at the follicle-paracortex interface). A subtractive color mixing model was employed to render these transitions, mimicking physical ink mixing on paper. Each cell type was assigned a base color (e.g., *C_A_* = [0, 0, 1] for B cells, *C_B_* = [1, 0, 0] for T cells). The “ink intensity” *I_k_* for a cell type at any spatial

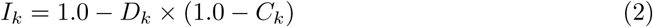

The final RGB color was computed by simulating ink overlay on a white background (*BG* = [1, 1, 1]):

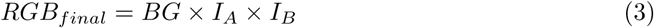

This density-overlay visualization encodes local cell abundance by color intensity and highlights regions of spatial overlap through color mixing, enabling clear identification of transition zones. Final ground-truth niche masks (B cell zone, GC, T cell zone, and medulla) were manually delineated from these density-gradient maps and verified against morphological landmarks on H&E staining. The GC annotation was additionally used as the ground truth for island niches.

### 5.2 Annotation of gradient niche

To characterize the continuous spatial transition during lymphocyte maturation, B-cell maturation zone was annotated by modeling the topological interface between naïve and memory B cells. We computed the transition intensity between these two cell populations and used it as a gradient norm, highlighting where the naïve*→*memory shift is most pronounced.

To guide manual annotation of the B-cell maturation zone, we visualized the spatial interface between naïve and memory B cells. A k-nearest neighbor graph (*k* = 30) was constructed, and potential fields were computed through signal diffusion. Binary signal fields were initialized for Naïve B cells (*f_A_*), Memory B cells (*f_B_*), and their union (*f_Sum_*). An adaptive-bandwidth affinity matrix was computed based on local density (using the *k*-th nearest neighbor distance as local scale *σ*), and signals were smoothed via iterative diffusion on the resulting transition matrix to yield continuous potential fields (*P_A_, P_B_, P_Sum_*). The gradient of each potential field (*∇P*) was computed in PCA space, and the transition gradient (*G_trans_*) was calculated to isolate the internal interface:

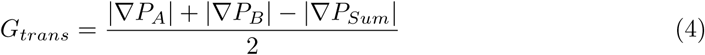

This geometric subtraction removes the gradient signal at the overall tissue boundary, specifically retaining the internal interface where the naïve*→*memory shift is most pronounced. The final transition score was weighted by local potential intensity and log-transformed:

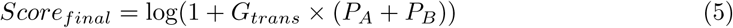

This score serves as the gradient norm to quantify the transition intensity at each spatial location. To facilitate interpretation of the complex gradient distribution, the transition score was thresholded at the 75th percentile to highlight the core transition zone for subsequent manual annotation.

Based on this transition intensity visualization, the B-cell maturation zone boundaries were manually drawn. The resulting annotated regions anatomically corresponded to the germinal center periphery and interfollicular zones, validating the biological relevance of this approach.

### 5.3 Simulation of biologically informed spatial niche data

We simulated a single-cell-resolution lymph node-like spatial transcriptomics dataset using SRTsim [55], with the manually annotated human lymph node reference serving as the template for expression characteristics and niche-aware structure. In brief, SRTsim fits gene-wise count models from the reference data and then generates synthetic counts on a user-defined spatial layout, while preserving spatial patterns through a reference-to-simulation mapping step that transfers expression structure between nearby locations.

We generated a synthetic tissue layout with four canonical niches—Medulla, T cell zone, B cell zone, and Germinal center—and specified biologically informed niche-wise cell-type composition priors to reflect lymph node organization. Under the baseline no-diffusion setting (i.e., maximal dominant-cell enrichment), each niche was assigned two dominant cell types with proportions chosen to mirror lymph node biology: in the Medulla, plasma cell 35.18% and macrophage 20.10%; in the T cell zone, CD4 T cell 47.95% and CD8 T cell 21.92%; in the B cell zone, Naïve B cell 58.21% and Memory B cell 31.34%; and in the Germinal center, GC B DZ 55.28% and GC B LZ 27.64%. Importantly, the no-diffusion baseline does not imply that niches are composed exclusively of dominant populations; rather, it represents the strongest dominant-cell enrichment while retaining non-dominant cell types to keep the simulation biologically realistic.

To quantify method sensitivity to compositional mixing within niches, we generated four datasets by progressively reducing dominant-cell proportion to 80/60/40% while preserving a non-trivial dominant signal. Let *p*_0_(*c | n*) denote the baseline cell-type probability for cell type *c* in niche *n*, and let *D_n_* denote the dominant cell types for niche *n*. For each factor

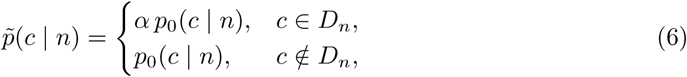

and then redistribute the removed mass to non-dominant types using a controlled mixing term Δ*_n_*(*c*), followed by renormalization,

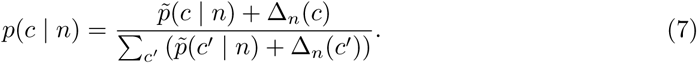

In practice, we constructed Δ*_n_*(*c*) to reflect realistic tissue heterogeneity by modeling it as a weighted mixture of spatial crosstalk and background noise. Specifically, 70% of the displaced probability mass was reallocated to dominant cell types characteristic of spatially adjacent compartments, while the remaining 30% was distributed among non-dominant background populations. This design ensures that lowering purity introduces spatially coherent interference without collapsing niche identity.

Expression was simulated with SRTsim’s domain-aware workflow, implemented with *createSRT*, *srtsim_fit*, and *srtsim_count*. We used a domain-specific simulation scheme so that gene-wise parameters were fit within niches rather than globally, and we simulated UMI-like overdispersed counts with a negative binomial (NB) marginal model (marginal = “nb”, sim_scheme = “domain”). Spatial pattern transfer was performed by mapping each simulated location to a neighborhood of reference locations using *k* = 50 nearest neighbors, and we generated counts using local aggregation with nn_num = 5 and nn_func = “mean” to stabilize the reference-to-simulation assignment. Across diffusion settings, the spatial layout and gene panel were held fixed; only the niche-wise cell-type composition priors were perturbed, allowing downstream benchmarking to isolate sensitivity to within-niche compositional diffusion rather than differences in spatial geometry or gene coverage.

### 5.4 Benchmarking methods

To provide a comprehensive evaluation of the current landscape of spatial niche identification, we benchmarked 16 representative algorithms, categorizing them into four distinct methodological frameworks based on their underlying mathematical and architectural principles: (i) Probabilistic & Statistical methods, (ii) Graph Neural Network (GNN) & Contrastive Learning approaches, (iii) Deep Generative models, and (iv) Foundation Models. A detailed description of each method is provided in the subsequent sections.

To ensure a rigorous and fair comparison, we standardized the experimental protocols across two distinct dimensions: input feature selection and cluster resolution. First, regarding input features, we implemented a stratified strategy to decouple algorithmic performance from data preprocessing. For our primary manually annotated Human Lymph Node dataset, we systematically evaluated the robustness of each method across four specific input scenarios: using all detected genes, the top 2,000 highly variable genes (HVGs), the top 2,000 spatially variable genes (SVGs), and a curated set of 2,000 genes prioritized via core-lineage-based differential expression. Conversely, for the remaining validation datasets, we adhered to the optimal feature selection strategy recommended by the original authors of each method (typically HVGs, SVGs, or full expression profiles) to prioritize their baseline performance. Second, to harmonize the granularity of niche identification, we aligned the output resolution with the ground truth. For methods requiring a predefined cluster number, *k* was set to four to match the intended output cardinality of the primary benchmark. For resolution-based clustering, the resolution was adjusted only until four valid non-empty domains were obtained; manual cell-level reference labels and agreement metrics were not used for hyperparameter tuning, model selection, seed selection, or best-run selection. Complete method-specific software versions, inputs, preprocessing procedures, neighborhood and graph construction, clustering settings, non-default parameters, computational devices, prespecified random seeds, and failure or fall-back rules are provided in Supplementary Table S1. A run was considered technically failed only if it encountered an execution or numerical error, returned missing or misaligned labels, or did not produce four valid non-empty domains under the declared procedure; low reference agreement was retained as a valid result. For the baseline runs of MENDER and CytoCommunity, the discrete cell-state inputs were the 16 curated cell-type labels defined in Fig. 2, together with the spatial coordinates; no separate expression-based cell clustering was used for these runs.

#### 5.4.1 Probabilistic & Statistical

##### BayesSpace

BayesSpace [15] is a fully Bayesian statistical framework that models low-dimensional gene expression representations using a Markov Random Field (MRF) prior. By employing a Potts model to encourage shared cluster assignments among neighbors, it explicitly enforces spatial smoothness to resolve continuous tissue structure despite technical noise.

##### BANKSY

BANKSY [16] leverages a spatially aware feature augmentation strategy to unify cell typing and domain segmentation without complex graph-based priors. It constructs a composite representation by concatenating a cell’s intrinsic profile with the weighted average of its local neighborhood, enabling the scalable identification of microenvironmental contexts via standard clustering.

##### MENDER

MENDER [17] introduces a scalable framework that avoids deep learning by explicitly encoding cellular context statistics. The method constructs a multi-range neighborhood representation by quantifying the distributional frequencies of cell states within multiple concentric spatial rings centered on each cell. These representations capture local heterogeneity across spatial scales and are subsequently clustered using standard clustering algorithms to delineate spatial domains.

#### 5.4.2 GNN & Contrastive

##### SpaGCN

SpaGCN [11] is a Graph Convolutional Network (GCN) [60] approach that integrates gene expression, spatial coordinates, and histology to identify spatial domains. It constructs a weighted graph based on physical and morphological proximity, utilizing graph convolution to aggregate information from neighboring spots and smooth technical variability.

##### GraphST

GraphST [19] combines graph neural networks with self-supervised contrastive learning to generate discriminative spot embeddings. By minimizing the embedding distance between spatially adjacent spots while maximizing global separability, the model effectively denoises data and delineates fine-grained tissue structures.

##### STAGATE

STAGATE [21] features a graph attention auto-encoder designed to learn low-dimensional latent embeddings by fusing gene expression and spatial information. Its core innovation is an adaptive attention mechanism that dynamically weighs the importance of neighboring spots, capturing non-linear spatial dependencies to sharpen domain boundaries.

##### CytoCommunity

CytoCommunity [20] is a graph pooling-based learning framework that identifies Tissue Cellular Neighborhoods (TCNs) directly from cell phenotypes and spatial distributions. Unlike multistep pipelines, it employs a graph neural network to learn a soft assignment matrix that maps individual cells directly to TCNs. This end-to-end architecture enables the de novo discovery of condition-specific multicellular communities of variable sizes.

##### SpaceFlow

SpaceFlow [22] generates spatially consistent low-dimensional embeddings by integrating spatial regularization into a deep graph network architecture. By incorporating a regularization term into the objective function that minimizes the latent distance between spatially adjacent cells, the framework effectively fuses gene expression patterns with spatial information.

#### 5.4.3 Deep Generative

##### NicheCompass

NicheCompass [12] integrates graph deep learning with biological prior knowledge to characterize tissue niches based on cellular communication potential. Unlike methods that rely solely on transcriptomic similarity, it constructs spatial graphs where edges encode potential ligand-receptor interactions derived from reference databases. By employing a message-passing mechanism over these interaction graphs, the model learns interpretable latent embeddings that explicitly capture signaling activity, enabling the identification of niches defined by distinct intercellular communication patterns rather than just cell type composition.

##### scNiche

scNiche [24] establishes a computational framework to identify cell niches at single-cell resolution by modeling the interplay between cells and their local microenvironments. Diverging from spot-level aggregation, scNiche constructs robust niche representations by integrating spatial proximity with the phenotypic features of neighboring cells, thereby enabling the precise delineation of spatial domains and quantitatively linking microenvironmental context to cellular phenotypes.

##### SEDR

SEDR [23] couples a deep autoencoder with a Variational Graph Autoencoder (VGAE) to generate low-dimensional latent representations. Transcriptomic features are learned via a masked self-supervised mechanism and subsequently integrated with spatial neighborhood graphs within the VGAE. This architecture enforces spatial coherence by aggregating information from adjacent spots, yielding a joint embedding that captures both molecular heterogeneity and tissue topology.

##### STACI

STACI [26] integrates gene expression profiles with nuclear morphological features via a graph-based autoencoder. The framework relies on over-parameterization within the latent space as a built-in mechanism to disentangle biological heterogeneity from technical batch effects, eliminating the need for external correction steps. By learning a joint representation that fuses transcriptomic and morphological modalities, STACI effectively captures complex tissue states and functional alterations.

##### DeepLinc

DeepLinc [25] utilizes a deep generative framework based on a Variational Graph Autoencoder (VGAE) to reconstruct cell interaction landscapes de novo. By integrating spatial connectivity with gene expression profiles, latent representations encoding spatially driven heterogeneity are learned from the data. Within this generative process, missing proximal and distal interactions are imputed while noise is filtered, yielding robust embeddings that facilitate the identification of cell niches defined by their specific interaction environments.

##### CellCharter

CellCharter [27] delineates spatial niches by modeling the composition of local cellular neighborhoods. A spatial graph is first constructed via Delaunay triangulation to define connectivity, after which a variational autoencoder (VAE) is used to learn a low-dimensional latent embedding of each cell’s expression profile, and composite feature vectors are generated by concatenating each cell’s intrinsic profile (latent embedding) with the aggregated features of its spatial neighbors. These spatially augmented representations are subsequently clustered using a Gaussian Mixture Model (GMM) to systematically identify distinct tissue microenvironments conserved across samples.

#### 5.4.4 Foundation Models

##### Novae

Novae [28] is a graph-based foundation model pre-trained on nearly 30 million spatially resolved cells spanning 18 tissues. It represents cells together with their local spatial neighborhoods and learns transferable graph representations through a prototype-based self-supervised objective adapted from SwAV. The resulting representation supports zero-shot spatial-domain assignment across gene panels, tissues, and technologies, as well as batch-aware integration and hierarchical domain resolution.

##### Nicheformer

Nicheformer [29] is a transformer-based foundation model pre-trained on SpatialCorpus-110M, which comprises more than 110 million human and mouse cells from dissociated and image-based spatial transcriptomic assays. Each cell is represented as an expression-ranked sequence of gene tokens augmented with organism, modality, and assay tokens, and the model is trained by reconstructing masked tokens. Physical coordinates are not used during pre-training; spatial niche and region labels are instead learned from the pre-trained cell representation through task-specific linear probing or fine-tuning.

### 5.5 Evaluation metrics

In this study, we quantified algorithm performance across Ground Truth Accuracy (ARI, AMI, Homogeneity, Completeness, Macro-F1), Biological Consistency (Celltype Cosine Sim), Spatial Structure (Spatial Connectivity), and Embedding Quality (Silhouette Score). In parallel, we evaluated the practical scalability of each method by measuring Computational Efficiency (Runtime, Peak Memory).

#### ARI

The Adjusted Rand Index (ARI) [61] measures the concordance between predicted spatial domains and ground truth annotations by correcting the standard Rand Index for chance agreement. Ranging from −1 to 1, a score of 1 indicates a perfect match, 0 implies random labeling, and negative values suggest systematic disagreement. This adjustment ensures robustness against variations in cluster size and number, providing a rigorous standard for partition similarity.

#### AMI

Adjusted Mutual Information (AMI) [62] extends the concept of NMI by correcting for the expected mutual information arising from chance, thereby addressing the selection bias inherent in comparing partitions with high cardinality. By subtracting the expected background score of random clusterings, AMI provides a stringent, unbiased assessment of biological fidelity, even when the number of detected niches is large relative to the sample size.

#### Homogeneity

Homogeneity [63] measures the purity of inferred niches by quantifying whether each niche contains samples from a single reference class. It is defined as

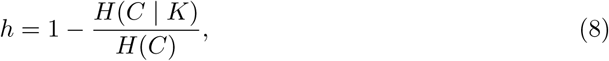

where *C* denotes the reference annotations and *K* denotes the predicted niche assignments. The score ranges from 0 to 1, with higher values indicating purer niches and stronger agreement with the reference labels.

#### Completeness

Completeness [63] evaluates the coherence of reference classes by assessing whether samples from the same class are assigned to a single inferred niche. It is defined as

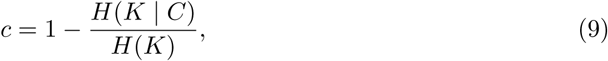

where *C* denotes the reference annotations and *K* denotes the predicted niche assignments. The score ranges from 0 to 1, with higher values indicating reduced fragmentation of reference classes across inferred niches.

#### Macro-F1 Score

To account for the arbitrary labeling inherent in unsupervised clustering, we first aligned the predicted spatial domains with ground truth annotations using the Hungarian algorithm to maximize the overlap between clusters. The Macro-F1 score was subsequently calculated as the harmonic mean of precision and recall, averaged equally across all unique ground truth categories. This metric ensures a balanced assessment that penalizes poor performance on rare niches:

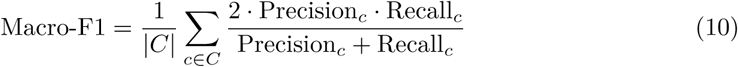

#### Composition Similarity

Following the alignment of predicted domains to ground truth niches via the Hungarian algorithm, we assessed biological consistency using Cell Type Cosine Similarity. For each matched pair of ground truth and predicted domains, we constructed cell type abundance vectors (*v_gt_*and *v_pred_*, respectively). The metric is defined as the mean cosine similarity between these aligned vectors, quantifying how accurately the algorithm reconstructs the cellular composition of each microenvironment:

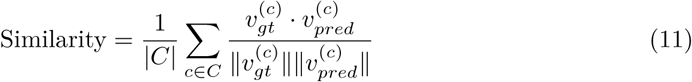

#### Spatial Connectivity

Connectivity quantifies the local spatial smoothness of the resulting partition. It is calculated as the average proportion of a cell’s spatial neighbors (*N_i_*) that are assigned to the same domain as the cell itself. This metric rewards spatial coherence and effectively penalizes fragmented classifications:

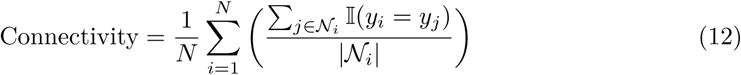

#### Silhouette Score

Silhouette Score [64] evaluates the quality of the learned latent representations by measuring intra-cluster cohesion and inter-cluster separation in the embedding space. For each cell, the score compares its average distance to other cells within the same predicted cluster with the average distance to cells in the nearest neighboring cluster. The Silhouette Score ranges from *−*1 to 1, with higher values indicating more compact and well-separated clusters.

#### Aggregate Performance Score and Sensitivity Analysis

To provide a compact summary of performance across complementary evaluation objectives, we additionally calculated an aggregate score for the primary lymph-node benchmark shown in Fig. 4e. For method *m* and metric *j*, metrics for which higher values indicate better performance—ARI, AMI, homogeneity, completeness, Macro-F1, composition similarity, spatial connectivity, and silhouette score—were min–max normalized across the 16 evaluated methods:

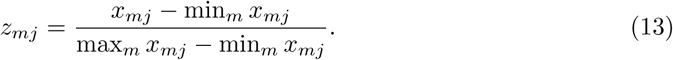

Runtime and peak memory were treated as cost metrics and reverse-normalized so that higher normalized values consistently indicated better performance:

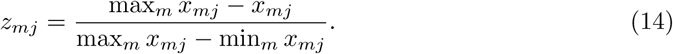

The five reference-agreement metrics were averaged to define the accuracy dimension:

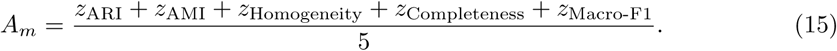

Biological consistency (*B_m_*), spatial structure (*S_m_*), and embedding quality (*E_m_*) were represented by the normalized composition similarity, spatial connectivity, and silhouette score, respectively. Computational efficiency (*C_m_*) was defined as the mean of the reverse-normalized runtime and peak memory:

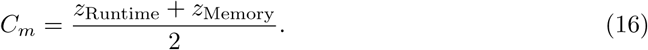

The aggregate score was then calculated as

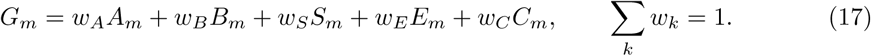

The score reported in Fig. 4e used equal weights for the five evaluation dimensions, with

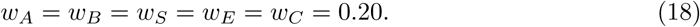

Thus, the five conceptual dimensions were weighted equally, whereas the ten individual component metrics were not assigned equal weights. The aggregate score was used as a descriptive multi-criteria summary; individual component metrics remained the primary basis for interpreting method performance.

All 16 methods included in this comparison completed successfully and had valid values for the required component metrics, and therefore no missing-value imputation was applied. If a required component metric is unavailable or a run fails, the aggregate score is considered not evaluable; missing components are not assigned zero values or compensated for by redistributing their weights. Failed scalability runs are reported separately and are not converted into aggregate-score penalties.

To assess the robustness of the aggregate ranking to metric redundancy and weighting choices, we first calculated pairwise Spearman correlations among the ten component metrics across the 16 methods. We then recomputed aggregate scores under six prespecified weighting schemes representing different evaluation priorities: equal-category, equal-metric, accuracy-focused, biology/spatial-focused, efficiency-focused, and scientific-performance-only weighting. The corresponding weights (*w_A_, w_B_, w_S_, w_E_, w_C_*) were (0.20, 0.20, 0.20, 0.20, 0.20), (0.50, 0.10, 0.10, 0.10, 0.20), (0.50, 0.125, 0.125, 0.125, 0.125), (0.25, 0.25, 0.25, 0.15, 0.10), (0.25, 0.125, 0.125, 0.10, 0.40), and (0.35, 0.25, 0.25, 0.15, 0), respectively. As a broader sensitivity analysis, we additionally evaluated 10,000 plausible combinations of the five category weights, constrained to sum to one with each weight ranging from 0.10 to 0.40. For each combination, aggregate scores were recalculated and methods were reranked. Ranking robustness was summarized using the median rank, interquartile range, 5th–95th percentile range, and the frequency with which each method appeared among the top three.

## 6 Declarations

### 6.1 Ethics approval and consent to participate

Not applicable. This study used publicly available spatial transcriptomics datasets. The original data collection and ethical approvals were obtained by the original data generators.

### 6.2 Consent for publication

Not applicable.

### 6.3 Availability of data and materials

The CosMx Human Lymph Node FFPE dataset used in this study is publicly available from NanoString/Bruker Spatial Biology at https://nanostring.com/products/ cosmx-spatial-molecular-imager/ffpe-dataset/cosmx-human-lymph-node-ffpe-dataset/. The 10x Genomics Visium DLPFC dataset (including slice 151673) was obtained from https://research.libd.org/spatialLIBD/. The MOSTA E16.5 mouse brain dataset was retrieved from the STOmicsDB MOSTA portal at https://db.cngb.org/stomics/mosta/. All benchmarking code, together with the manual niche annotations for the human lymph node reference, is publicly available at https://github.com/WYXNICK/spatial-niche-benchmark.

### 6.6 Authors’ contributions

Y.W., J.C., H.X., and Y.C. conceived the study. L.Y. and C.W. investigated the data and methods. Y.C. performed the data annotation. Y.W. implemented the framework, conducted the experiments, and analyzed the results. Y.W., H.X., and Y.C. wrote the manuscript. J.C. and H.X. revised the manuscript. H.X. and J.C. supervised the project. H.X. acquired funding for this research. All authors reviewed and approved the manuscript.

### 6.7 Acknowledgements

We thank all members who contributed to this study for their valuable support and discussion.

## References

[1] Scadden, D.T.: Nice neighborhood: emerging concepts of the stem cell niche. Cell 157(1), 41–50 (2014)

[2] Morrison, S.J., Spradling, A.C.: Stem cells and niches: mechanisms that promote stem cell maintenance throughout life. Cell 132(4), 598–611 (2008)

[3] De Martin, A., Stanossek, Y., Pikor, N.B., Ludewig, B.: Protective fibroblastic niches in secondary lymphoid organs. Journal of Experimental Medicine 221(1), 20221220 (2023)

[4] Alexandre, Y.O., Mueller, S.N.: Stromal cell networks coordinate immune response generation and maintenance. Immunological reviews 283(1), 77–85 (2018)

[5] Kicheva, A., Briscoe, J.: Control of tissue development by morphogens. Annual review of cell and developmental biology 39(1), 91–121 (2023)

[6] Ståhl, P.L., Salmén, F., Vickovic, S., Lundmark, A., Navarro, J.F., Magnusson, J., Giacomello, S., Asp, M., Westholm, J.O., Huss, M., et al.: Visualization and analysis of gene expression in tissue sections by spatial transcriptomics. Science 353(6294), 78–82 (2016)

[7] Chen, K.H., Boettiger, A.N., Moffitt, J.R., Wang, S., Zhuang, X.: Spatially resolved, highly multiplexed rna profiling in single cells. Science 348(6233), 6090 (2015)

[8] Clevers, H.: The intestinal crypt, a prototype stem cell compartment. Cell 154(2), 274–284 (2013)

[9] Sato, T., Van Es, J.H., Snippert, H.J., Stange, D.E., Vries, R.G., Van Den Born, M., Barker, N., Shroyer, N.F., Van De Wetering, M., Clevers, H.: Paneth cells constitute the niche for lgr5 stem cells in intestinal crypts. Nature 469(7330), 415–418 (2011)

[10] Moses, L., Pachter, L.: Museum of spatial transcriptomics. Nature methods 19(5), 534–546 (2022)

[11] Hu, J., Li, X., Coleman, K., Schroeder, A., Ma, N., Irwin, D.J., Lee, E.B., Shinohara, R.T., Li, M.: Spagcn: Integrating gene expression, spatial location and histology to identify spatial domains and spatially variable genes by graph convolutional network. Nature methods 18(11), 1342–1351 (2021)

[12] Birk, S., Bonafonte-Pardàs, I., Feriz, A.M., Boxall, A., Agirre, E., Memi, F., Maguza, A., Yadav, A., Armingol, E., Fan, R., et al.: Quantitative characterization of cell niches in spatially resolved omics data. Nature Genetics, 1–13 (2025)

[13] Bhate, S.S., Barlow, G.L., Schürch, C.M., Nolan, G.P.: Tissue schematics map the specialization of immune tissue motifs and their appropriation by tumors. Cell Systems 13(2), 109–130 (2022)

[14] Armingol, E., Officer, A., Harismendy, O., Lewis, N.E.: Deciphering cell–cell interactions and communication from gene expression. Nature Reviews Genetics 22(2), 71–88 (2021)

[15] Zhao, E., Stone, M.R., Ren, X., Guenthoer, J., Smythe, K.S., Pulliam, T., Williams, S.R., Uytingco, C.R., Taylor, S.E., Nghiem, P., et al.: Spatial transcriptomics at subspot resolution with bayesspace. Nature biotechnology 39(11), 1375–1384 (2021)

[16] Singhal, V., Chou, N., Lee, J., Yue, Y., Liu, J., Chock, W.K., Lin, L., Chang, Y.-C., Teo, E.M.L., Aow, J., et al.: Banksy unifies cell typing and tissue domain segmentation for scalable spatial omics data analysis. Nature genetics 56(3), 431–441 (2024)

[17] Yuan, Z.: Mender: fast and scalable tissue structure identification in spatial omics data. Nature Communications 15(1), 207 (2024)

[18] Scarselli, F., Gori, M., Tsoi, A.C., Hagenbuchner, M., Monfardini, G.: The graph neural network model. IEEE transactions on neural networks 20(1), 61–80 (2008)

[19] Long, Y., Ang, K.S., Li, M., Chong, K.L.K., Sethi, R., Zhong, C., Xu, H., Ong, Z., Sachaphibulkij, K., Chen, A., et al.: Spatially informed clustering, integration, and deconvolution of spatial transcriptomics with graphst. Nature Communications 14(1), 1155 (2023)

[20] Hu, Y., Rong, J., Xu, Y., Xie, R., Peng, J., Gao, L., Tan, K.: Unsupervised and supervised discovery of tissue cellular neighborhoods from cell phenotypes. Nature Methods 21(2), 267–278 (2024)

[21] Dong, K., Zhang, S.: Deciphering spatial domains from spatially resolved transcriptomics with an adaptive graph attention auto-encoder. Nature communications 13(1), 1739 (2022)

[22] Ren, H., Walker, B.L., Cang, Z., Nie, Q.: Identifying multicellular spatiotemporal organization of cells with spaceflow. Nature communications 13(1), 4076 (2022)

[23] Xu, H., Fu, H., Long, Y., Ang, K.S., Sethi, R., Chong, K., Li, M., Uddamvathanak, R., Lee, H.K., Ling, J., et al.: Unsupervised spatially embedded deep representation of spatial transcriptomics. Genome Medicine 16(1), 12 (2024)

[24] Qian, J., Shao, X., Bao, H., Fang, Y., Guo, W., Li, C., Li, A., Hua, H., Fan, X.: Identification and characterization of cell niches in tissue from spatial omics data at single-cell resolution. Nature Communications 16(1), 1693 (2025)

[25] Li, R., Yang, X.: De novo reconstruction of cell interaction landscapes from single-cell spatial transcriptome data with deeplinc. Genome biology 23(1), 124 (2022)

[26] Zhang, X., Wang, X., Shivashankar, G., Uhler, C.: Graph-based autoencoder integrates spatial transcriptomics with chromatin images and identifies joint biomarkers for alzheimer’s disease. Nature Communications 13(1), 7480 (2022)

[27] Varrone, M., Tavernari, D., Santamaria-Martínez, A., Walsh, L.A., Ciriello, G.: Cellcharter reveals spatial cell niches associated with tissue remodeling and cell plasticity. Nature genetics 56(1), 74–84 (2024)

[28] Blampey, Q., Benkirane, H., Bercovici, N., Mulder, K., Gessain, G., Ginhoux, F., André, F., Cournède, P.-H.: Novae: a graph-based foundation model for spatial transcriptomics data. Nature Methods 22(12), 2539–2550 (2025)

[29] Tejada-Lapuerta, A., Schaar, A.C., Gutgesell, R., Palla, G., Halle, L., Minaeva, M., Vornholz, L., Dony, L., Drummer, F., Richter, T., et al.: Nicheformer: a foundation model for single-cell and spatial omics. Nature methods, 1–14 (2025)

[30] He, S., Bhatt, R., Brown, C., Brown, E.A., Buhr, D.L., Chantranuvatana, K., Danaher, P., Dunaway, D., Garrison, R.G., Geiss, G., et al.: High-plex imaging of rna and proteins at subcellular resolution in fixed tissue by spatial molecular imaging. Nature biotechnology 40(12), 1794–1806 (2022)

[31] Maynard, K.R., Collado-Torres, L., Weber, L.M., Uytingco, C., Barry, B.K., Williams, S.R., Catallini, J.L., Tran, M.N., Besich, Z., Tippani, M., et al.: Transcriptome-scale spatial gene expression in the human dorsolateral prefrontal cortex. Nature neuroscience 24(3), 425–436 (2021)

[32] Chen, A., Liao, S., Cheng, M., Ma, K., Wu, L., Lai, Y., Qiu, X., Yang, J., Xu, J., Hao, S., et al.: Spatiotemporal transcriptomic atlas of mouse organogenesis using dna nanoball-patterned arrays. Cell 185(10), 1777–1792 (2022)

[33] Liao, S., Padera, T.P.: Lymphatic function and immune regulation in health and disease. Lymphatic research and biology 11(3), 136–143 (2013)

[34] Grant, S.M., Lou, M., Yao, L., Germain, R.N., Radtke, A.J.: The lymph node at a glance–how spatial organization optimizes the immune response. Journal of cell science 133(5), 241828 (2020)

[35] Mohseni, S., Shojaiefard, A., Khorgami, Z., Alinejad, S., Ghorbani, A., Ghafouri, A.: Peripheral lymphadenopathy: approach and diagnostic tools. Iranian journal of medical sciences 39(2 Suppl), 158 (2014)

[36] Dasoveanu, D.C., Shipman, W.D., Chia, J.J., Chyou, S., Lu, T.T.: Regulation of lymph node vascular–stromal compartment by dendritic cells. Trends in immunology 37(11), 764–777 (2016)

[37] De Silva, N.S., Klein, U.: Dynamics of b cells in germinal centres. Nature reviews immunology 15(3), 137–148 (2015)

[38] Pereira, J.P., Kelly, L.M., Cyster, J.G.: Finding the right niche: B-cell migration in the early phases of t-dependent antibody responses. International immunology 22(6), 413–419 (2010)

[39] Allen, C.D., Okada, T., Cyster, J.G.: Germinal-center organization and cellular dynamics. Immunity 27(2), 190–202 (2007)

[40] Rodda, L.B., Lu, E., Bennett, M.L., Sokol, C.L., Wang, X., Luther, S.A., Barres, B.A., Luster, A.D., Ye, C.J., Cyster, J.G.: Single-cell rna sequencing of lymph node stromal cells reveals niche-associated heterogeneity. Immunity 48(5), 1014–1028 (2018)

[41] Girard, J.-P., Moussion, C., Förster, R.: Hevs, lymphatics and homeostatic immune cell trafficking in lymph nodes. Nature Reviews Immunology 12(11), 762–773 (2012)

[42] Phan, T.G., Green, J.A., Gray, E.E., Xu, Y., Cyster, J.G.: Immune complex relay by subcapsular sinus macrophages and noncognate b cells drives antibody affinity maturation. Nature immunology 10(7), 786–793 (2009)

[43] Linton, M.F., Babaev, V.R., Huang, J., Linton, E.F., Tao, H., Yancey, P.G.: Macrophage apoptosis and efferocytosis in the pathogenesis of atherosclerosis. Circulation Journal 80(11), 2259–2268 (2016)

[44] Gallo, P., Gonçalves, R., Mosser, D.M.: The influence of igg density and macrophage fc (gamma) receptor cross-linking on phagocytosis and il-10 production. Immunology letters 133(2), 70–77 (2010)

[45] Kawai, Y., Ouchida, R., Yamasaki, S., Dragone, L., Tsubata, T., Wang, J.-Y.: Laptm5 promotes lysosomal degradation of intracellular cd3*ζ* but not of cell surface cd3*ζ*. Immunology and cell biology 92(6), 527–534 (2014)

[46] Gulati, G.S., D’Silva, J.P., Liu, Y., Wang, L., Newman, A.M.: Profiling cell identity and tissue architecture with single-cell and spatial transcriptomics. Nature reviews Molecular cell biology 26(1), 11–31 (2025)

[47] Zhang, W., Brown, E.L., Usmani, A., Earland, N., Kang, M., Olelewe, C., Viswanathan, A., Chauhan, P.S., Steen, C.B., Jeon, H.S., et al.: Non-invasive profiling of the tumour microenvironment with spatial ecotypes. Nature, 1–12 (2026)

[48] McInnes, L., Healy, J., Melville, J.: Umap: Uniform manifold approximation and projection for dimension reduction. arXiv preprint arXiv:1802.03426 (2018)

[49] Cover, T., Hart, P.: Nearest neighbor pattern classification. IEEE transactions on information theory 13(1), 21–27 (1967)

[50] Hess, N.J., Jiang, S., Li, X., Guan, Y., Tapping, R.I.: Tlr10 is a b cell intrinsic suppressor of adaptive immune responses. The Journal of Immunology 198(2), 699–707 (2017)

[51] Cosgrove, J., Novkovic, M., Albrecht, S., Pikor, N.B., Zhou, Z., Onder, L., Mörbe, U., Cupovic, J., Miller, H., Alden, K., et al.: B cell zone reticular cell microenvironments shape cxcl13 gradient formation. Nature communications 11(1), 3677 (2020)

[52] Lyubchenko, T., Dal Porto, J., Cambier, J.C., Holers, V.M.: Coligation of the b cell receptor with complement receptor type 2 (cr2/cd21) using its natural ligand c3dg: activation without engagement of an inhibitory signaling pathway. The Journal of Immunology 174(6), 3264–3272 (2005)

[53] Palanichamy, A., Barnard, J., Zheng, B., Owen, T., Quach, T., Wei, C., Looney, R.J., Sanz, I., Anolik, J.H.: Novel human transitional b cell populations revealed by b cell depletion therapy. The Journal of Immunology 182(10), 5982–5993 (2009)

[54] Zhang, X., Ma, C., Lu, Y., Wang, J., Yun, H., Jiang, H., Wu, M., Feng, X., Gai, W., Xu, G., et al.: Rack1 regulates b-cell development and function by binding to and stabilizing the transcription factor pax5. Cellular & Molecular Immunology 21(11), 1282–1295 (2024)

[55] Zhu, J., Shang, L., Zhou, X.: Srtsim: spatial pattern preserving simulations for spatially resolved transcriptomics. Genome biology 24(1), 39 (2023)

[56] Kleshchevnikov, V., Shmatko, A., Dann, E., Aivazidis, A., King, H.W., Li, T., Elmentaite, R., Lomakin, A., Kedlian, V., Gayoso, A., et al.: Cell2location maps fine-grained cell types in spatial transcriptomics. Nature biotechnology 40(5), 661–671 (2022)

[57] Zhou, X., Dong, K., Zhang, S.: Integrating spatial transcriptomics data across different conditions, technologies and developmental stages. Nature Computational Science 3(10), 894–906 (2023)

[58] Guo, Z.-H., Huang, D.-S., Zhang, S.: Multi-task spatial omics analytics for high-precision modeling of tissue architecture with stax. bioRxiv, 2025–12 (2025)

[59] Virtanen, P., Gommers, R., Oliphant, T.E., Haberland, M., Reddy, T., Cournapeau, D., Burovski, E., Peterson, P., Weckesser, W., Bright, J., et al.: Scipy 1.0: fundamental algorithms for scientific computing in python. Nature methods 17(3), 261–272 (2020)

[60] Kipf, T.: Semi-supervised classification with graph convolutional networks. arXiv preprint arXiv:1609.02907 (2016)

[61] Hubert, L., Arabie, P.: Comparing partitions. Journal of classification 2(1), 193–218 (1985)

[62] Vinh, N.X., Epps, J., Bailey, J.: Information theoretic measures for clusterings comparison: Variants, properties, normalization and correction for chance. Journal of Machine Learning Research 11(95), 2837–2854 (2010)

[63] Rosenberg, A., Hirschberg, J.: V-measure: A conditional entropy-based external cluster evaluation measure. In: Proceedings of the 2007 Joint Conference on Empirical Methods in Natural Language Processing and Computational Natural Language Learning (EMNLP-CoNLL), pp. 410–420 (2007)

[64] Rousseeuw, P.J.: Silhouettes: a graphical aid to the interpretation and validation of cluster analysis. Journal of computational and applied mathematics 20, 53–65 (1987)

