## Supplementary Figures for "Benchmarking niche identification via domain segmentation for spatial transcriptomics data"

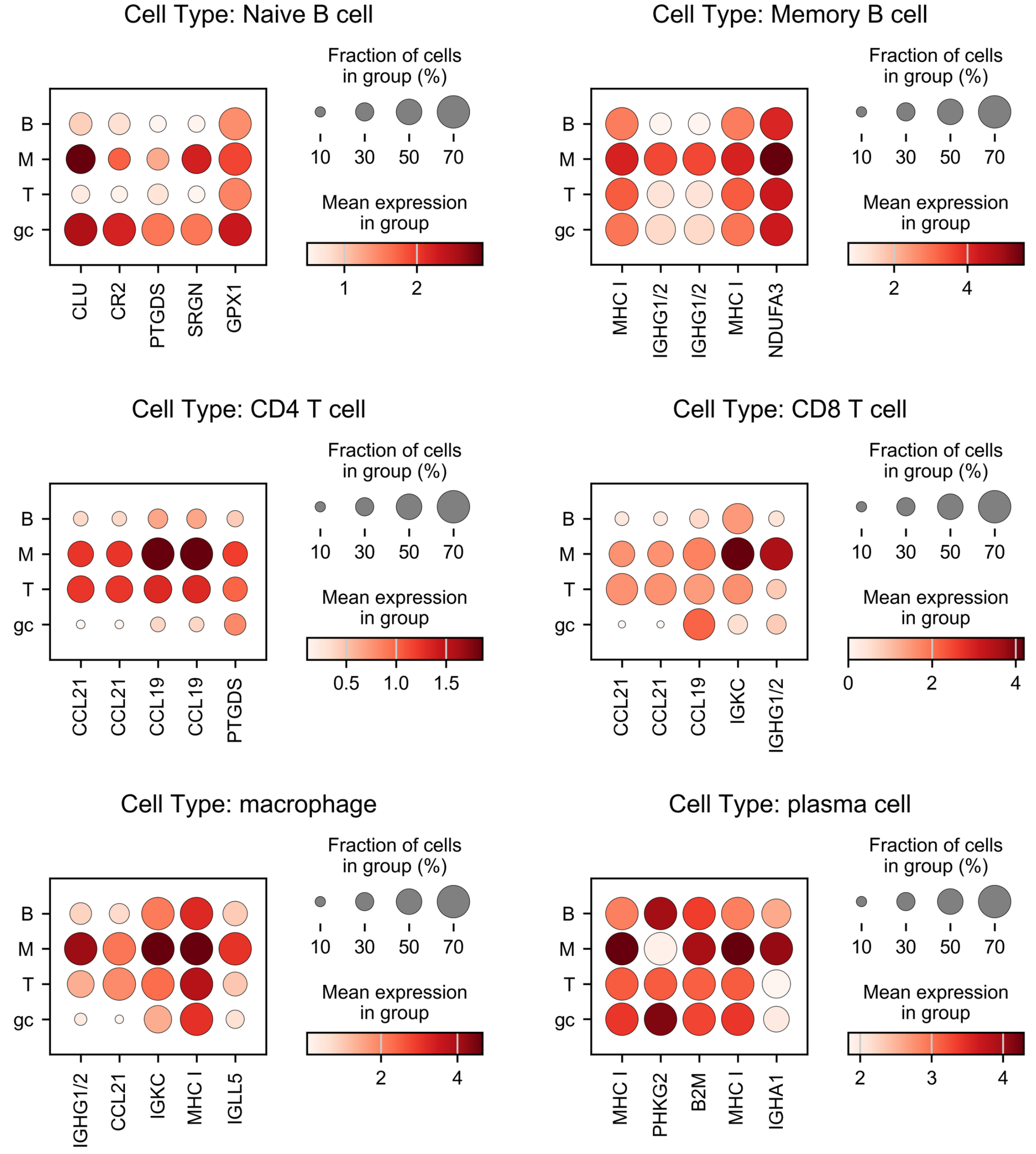


**Supplementary Fig. S1.** Gene expression patterns of major cell types across lymph node niches. Dot plots show the expression of representative marker genes for major immune cell types (Naive B cell, Memory B cell, CD4 T cell, CD8 T cell, macrophage, and plasma cell) across four annotated niches: T cell Zone (T), B cell Zone (B), Medulla (M), and Germinal Center (GC). Dot color indicates the mean (scaled) expression level of each gene within the niche-specific subset of the given cell type, and dot size indicates the fraction of cells expressing the gene in that group (%).

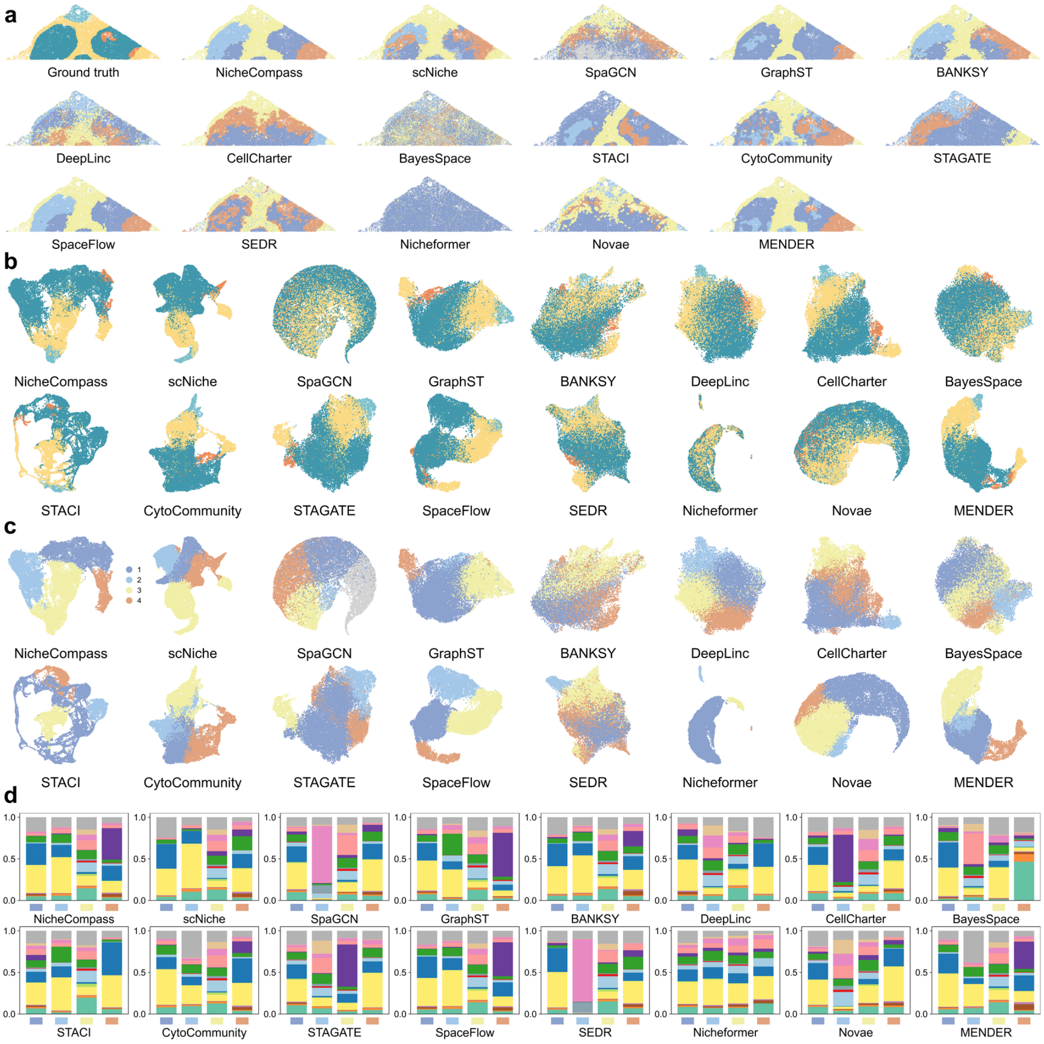


**e**

| | Ground Truth Accuracy | | | | | Biological Consistency | Spatial Structure | Embedding Quality | Computational Efficiency | | Aggregate |
|---|---|---|---|---|---|---|---|---|---|---|---|
| **Method** | ARI | AMI | Homogeneity | Completeness | Macro-F1 | Composition Similarity | Spatial Connectivity | Silhouette Score | Time (s) | Memory (GB) | Total |
| **MENDER** | 0.37 | 0.34 | 0.41 | 0.30 | 0.46 | 0.74 | 0.96 | 0.15 | 158 | 0.16 | 0.860 |
| **CytoCommunity** | 0.20 | 0.25 | 0.32 | 0.21 | 0.37 | 0.74 | 0.86 | 0.15 | 832 | 6.48 | 0.736 |
| **Novae** | 0.16 | 0.17 | 0.19 | 0.15 | 0.44 | 0.82 | 0.86 | 0.00 | 70.9 | 0.32 | 0.729 |
| **scNiche** | 0.25 | 0.37 | 0.47 | 0.31 | 0.39 | 0.69 | 0.88 | 0.10 | 96.9 | 0.49 | 0.720 |
| **BANKSY** | 0.22 | 0.32 | 0.40 | 0.26 | 0.40 | 0.72 | 0.94 | 0.08 | 204 | 6.02 | 0.720 |
| **GraphST** | 0.53 | 0.44 | 0.46 | 0.42 | 0.44 | 0.66 | 0.97 | 0.11 | 57.7 | 15.7 | 0.706 |
| **NicheCompass** | 0.24 | 0.32 | 0.39 | 0.27 | 0.38 | 0.69 | 0.97 | 0.04 | 660 | 0.27 | 0.683 |
| **STAGATE** | 0.13 | 0.18 | 0.20 | 0.16 | 0.34 | 0.69 | 0.91 | 0.11 | 368 | 0.96 | 0.662 |
| **SpaceFlow** | 0.27 | 0.33 | 0.41 | 0.28 | 0.39 | 0.69 | 0.97 | 0.01 | 540 | 1.01 | 0.660 |
| **SEDR** | 0.31 | 0.33 | 0.39 | 0.29 | 0.43 | 0.70 | 0.86 | 0.06 | 57.5 | 9.28 | 0.652 |
| **CellCharter** | 0.20 | 0.24 | 0.28 | 0.21 | 0.31 | 0.65 | 0.90 | 0.11 | 1374 | 0.19 | 0.628 |
| **STACI** | 0.05 | 0.20 | 0.22 | 0.18 | 0.32 | 0.67 | 0.96 | 0.05 | 288 | 3.56 | 0.602 |
| **Nicheformer** | 0.01 | 0.00 | 0.00 | 0.00 | 0.22 | 0.68 | 0.88 | 0.04 | 2704 | 3.21 | 0.470 |
| **DeepLinc** | 0.12 | 0.19 | 0.23 | 0.15 | 0.36 | 0.79 | 0.73 | -0.03 | 14440 | 25.0 | 0.417 |
| **SpaGCN** | 0.09 | 0.13 | 0.17 | 0.11 | 0.29 | 0.68 | 0.74 | -0.09 | 312 | 7.60 | 0.373 |
| **BayesSpace** | 0.12 | 0.09 | 0.11 | 0.08 | 0.30 | 0.65 | 0.43 | 0.00 | 1727 | 0.18 | 0.310 |

**Supplementary Fig. S2.** Benchmarking results on the manually annotated human lymph node dataset using the **HVG-2000** augmentation strategy. (a) Spatial niche assignments for the ground truth and each method, visualized in tissue coordinates. (b-c) UMAP visualization of method-specific latent embeddings, with cells colored by ground-truth niche labels (b) and by each method's predicted niche labels (c). (d) Cell-type composition of niches for each method, shown as the proportion of annotated cell types within each niche. (e) Multi-criteria quantitative comparison across ground-truth accuracy, biological consistency, spatial structure, embedding quality, computational efficiency, and the aggregated overall score.

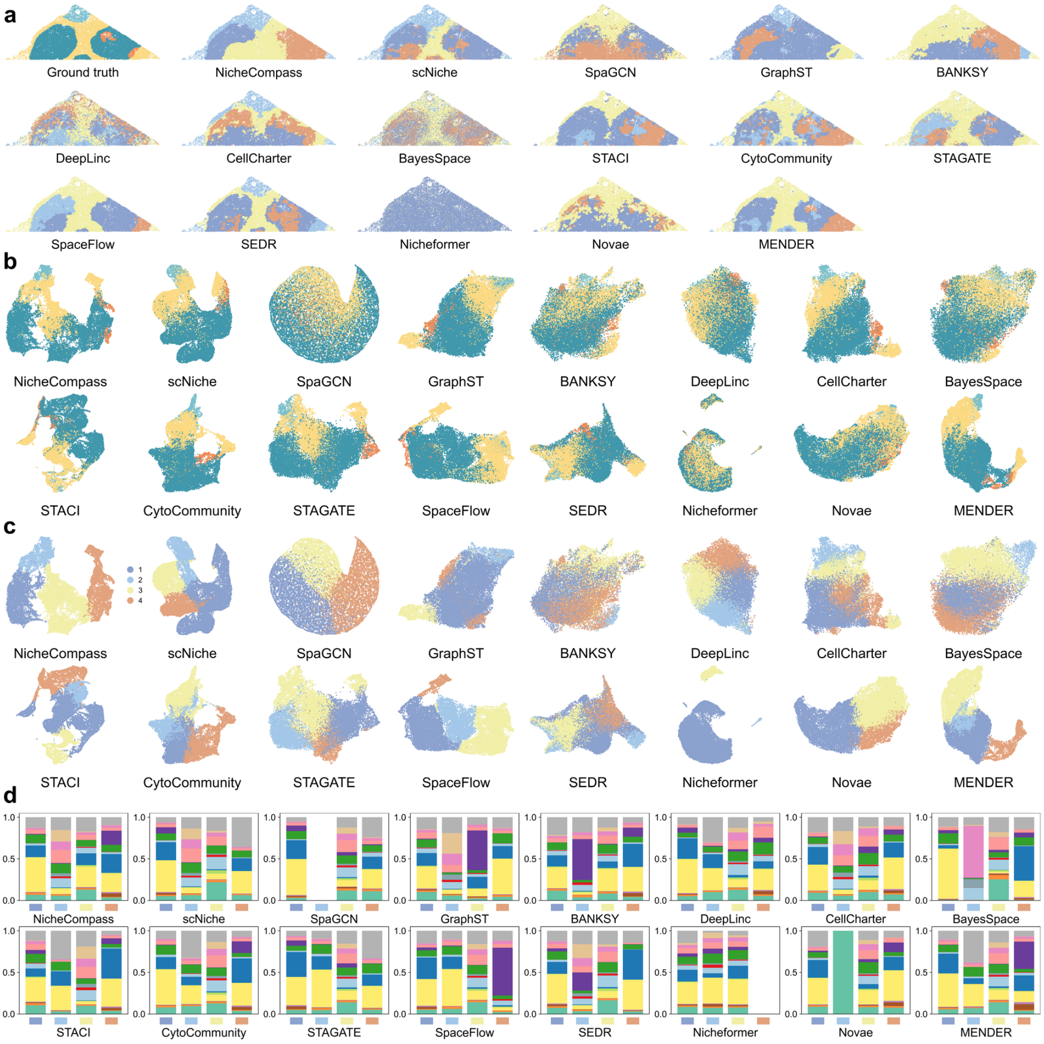


| Method | Ground Truth Accuracy: ARI | AMI | Homogeneity | Completeness | Macro-F1 | Biological Consistency: Composition Similarity | Spatial Structure: Spatial Connectivity | Embedding Quality: Silhouette Score | Computational Efficiency: Time (s) | Memory (GB) | Aggregate: Total |
|---|---|---|---|---|---|---|---|---|---|---|---|
| CellCharter | 0.22 | 0.27 | 0.34 | 0.23 | 0.47 | 0.85 | 0.89 | 0.11 | 1372 | 0.19 | 0.864 |
| MENDER | 0.37 | 0.34 | 0.41 | 0.30 | 0.46 | 0.74 | 0.96 | 0.15 | 124 | 0.15 | 0.857 |
| SEDR | 0.36 | 0.42 | 0.50 | 0.36 | 0.53 | 0.80 | 0.94 | 0.10 | 57.1 | 9.28 | 0.857 |
| scNiche | 0.31 | 0.34 | 0.41 | 0.30 | 0.45 | 0.79 | 0.89 | 0.05 | 107 | 0.49 | 0.801 |
| NicheCompass | 0.13 | 0.25 | 0.31 | 0.21 | 0.42 | 0.81 | 0.98 | 0.06 | 970 | 0.27 | 0.800 |
| CytoCommunity | 0.20 | 0.25 | 0.32 | 0.21 | 0.37 | 0.74 | 0.86 | 0.15 | 825 | 6.48 | 0.728 |
| STAGATE | 0.20 | 0.33 | 0.41 | 0.27 | 0.34 | 0.67 | 0.90 | 0.10 | 363 | 0.96 | 0.687 |
| SpaceFlow | 0.32 | 0.40 | 0.46 | 0.34 | 0.36 | 0.63 | 0.97 | -0.01 | 518 | 1.06 | 0.638 |
| GraphST | 0.10 | 0.23 | 0.23 | 0.22 | 0.39 | 0.72 | 0.97 | 0.02 | 45.8 | 15.4 | 0.620 |
| Novae | 0.15 | 0.15 | 0.16 | 0.14 | 0.37 | 0.65 | 0.88 | 0.10 | 73.8 | 0.10 | 0.619 |
| STACI | 0.11 | 0.28 | 0.33 | 0.25 | 0.30 | 0.68 | 0.95 | -0.03 | 229 | 3.52 | 0.586 |
| SpaGCN | 0.19 | 0.21 | 0.23 | 0.19 | 0.31 | 0.82 | 0.86 | -0.13 | 349 | 7.60 | 0.586 |
| BANKSY | 0.04 | 0.14 | 0.16 | 0.12 | 0.30 | 0.63 | 0.95 | 0.08 | 208 | 6.02 | 0.567 |
| Nicheformer | 0.00 | 0.00 | 0.00 | 0.00 | 0.22 | 0.70 | 0.89 | 0.03 | 2807 | 3.23 | 0.508 |
| BayesSpace | 0.15 | 0.14 | 0.17 | 0.12 | 0.34 | 0.65 | 0.48 | 0.00 | 1690 | 0.19 | 0.375 |
| DeepLinc | 0.12 | 0.12 | 0.15 | 0.10 | 0.32 | 0.72 | 0.68 | -0.01 | 16608 | 24.8 | 0.312 |

**Supplementary Fig. S3.** Benchmarking results on the manually annotated human lymph node dataset using the **SVG-2000** augmentation strategy. (a) Spatial niche assignments for the ground truth and each method, visualized in tissue coordinates. (b-c) UMAP visualization of method-specific latent embeddings, with cells colored by ground-truth niche labels (b) and by each method's predicted niche labels (c). (d) Cell-type composition of niches for each method, shown as the proportion of annotated cell types within each niche. (e) Multi-criteria quantitative comparison across ground-truth accuracy, biological consistency, spatial structure, embedding quality, computational efficiency, and the aggregated overall score.

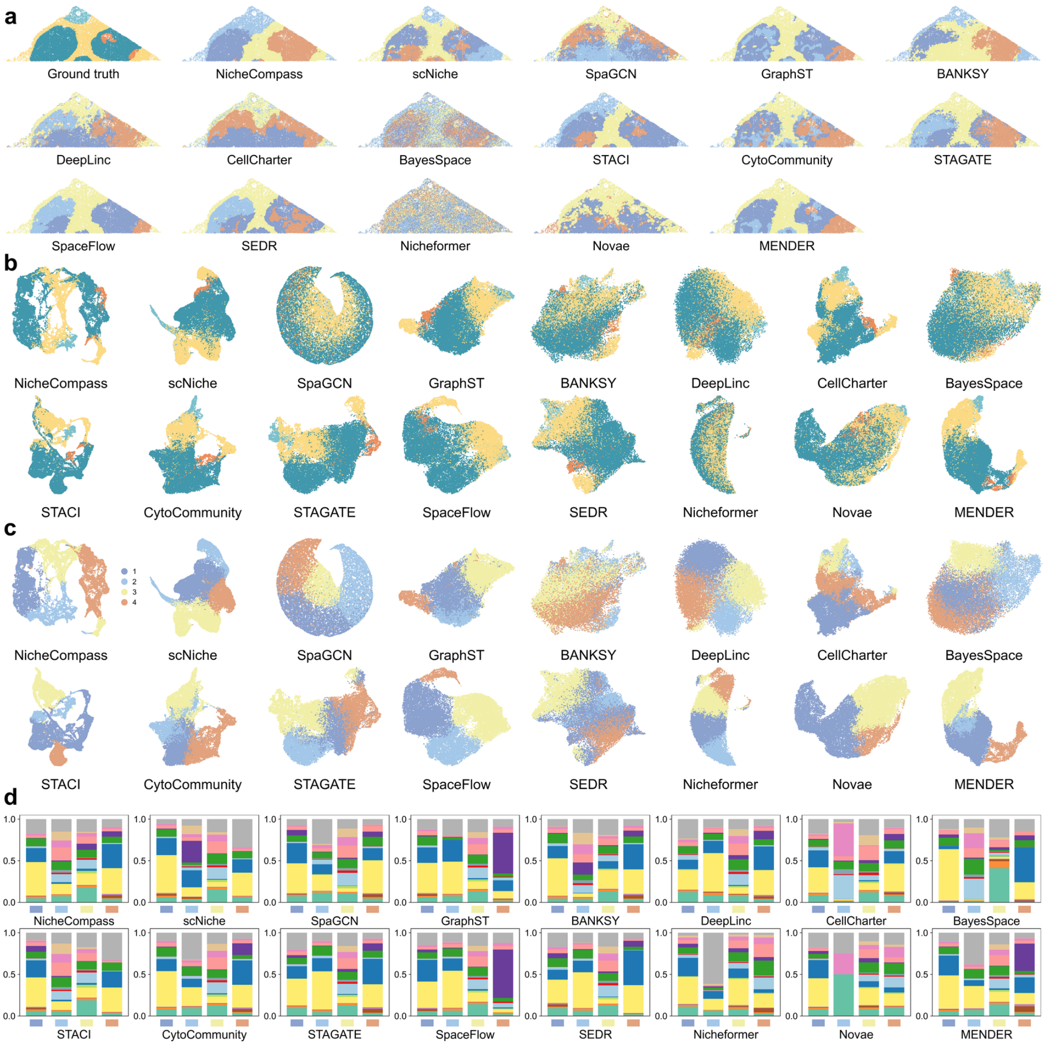


e

| | Ground Truth Accuracy | | | | | Biological Consistency | Spatial Structure | Embedding Quality | Computational Efficiency | | Aggregate |
|---|---|---|---|---|---|---|---|---|---|---|---|
| Method | ARI | AMI | Homogeneity | Completeness | Macro-F1 | Composition Similarity | Spatial Connectivity | Silhouette Score | Time (s) | Memory (GB) | Total |
| STACI | 0.31 | 0.33 | 0.38 | 0.29 | 0.48 | 0.80 | 0.96 | 0.12 | 314 | 3.56 | 0.895 |
| NicheCompass | 0.28 | 0.38 | 0.49 | 0.32 | 0.47 | 0.83 | 0.97 | 0.04 | 2698 | 0.28 | 0.889 |
| MENDER | 0.37 | 0.34 | 0.41 | 0.30 | 0.46 | 0.74 | 0.96 | 0.15 | 161 | 0.17 | 0.882 |
| BANKSY | 0.12 | 0.27 | 0.33 | 0.23 | 0.42 | 0.80 | 0.94 | 0.08 | 82.4 | 6.02 | 0.808 |
| STAGATE | 0.23 | 0.35 | 0.44 | 0.29 | 0.38 | 0.70 | 0.90 | 0.12 | 661 | 0.96 | 0.773 |
| scNiche | 0.32 | 0.33 | 0.41 | 0.28 | 0.46 | 0.71 | 0.88 | 0.06 | 172 | 0.49 | 0.756 |
| CytoCommunity | 0.20 | 0.25 | 0.32 | 0.21 | 0.37 | 0.74 | 0.86 | 0.15 | 778 | 6.48 | 0.754 |
| CellCharter | 0.21 | 0.25 | 0.29 | 0.22 | 0.36 | 0.72 | 0.90 | 0.12 | 14.0 | 0.14 | 0.753 |
| SEDR | 0.22 | 0.35 | 0.44 | 0.29 | 0.32 | 0.67 | 0.94 | 0.12 | 72.8 | 9.28 | 0.715 |
| SpaceFlow | 0.32 | 0.40 | 0.47 | 0.35 | 0.37 | 0.63 | 0.97 | 0.01 | 669 | 1.01 | 0.680 |
| GraphST | 0.33 | 0.36 | 0.42 | 0.31 | 0.39 | 0.65 | 0.94 | 0.04 | 58.6 | 15.5 | 0.649 |
| Novae | 0.18 | 0.15 | 0.16 | 0.15 | 0.33 | 0.66 | 0.88 | 0.09 | 521 | 0.09 | 0.634 |
| SpaGCN | 0.14 | 0.20 | 0.25 | 0.16 | 0.30 | 0.72 | 0.81 | -0.15 | 170 | 4.56 | 0.500 |
| BayesSpace | 0.14 | 0.13 | 0.16 | 0.11 | 0.31 | 0.67 | 0.44 | 0.01 | 1904 | 0.18 | 0.419 |
| DeepLinc | 0.12 | 0.18 | 0.23 | 0.15 | 0.30 | 0.67 | 0.82 | -0.03 | 15340 | 24.3 | 0.352 |
| Nicheformer | 0.02 | 0.01 | 0.01 | 0.01 | 0.24 | 0.62 | 0.37 | 0.02 | 499 | 1.83 | 0.301 |

**Supplementary Fig. S4.** Benchmarking results on the manually annotated human lymph node dataset using the **curated genes** strategy. (a) Spatial niche assignments for the ground truth and each method, visualized in tissue coordinates. (b-c) UMAP visualization of method-specific latent embeddings, with cells colored by ground-truth niche labels (b) and by each method's predicted niche labels (c). (d) Cell-type composition of niches for each method, shown as the proportion of annotated cell types within each niche. (e) Multi-criteria quantitative comparison across ground-truth accuracy, biological consistency, spatial structure, embedding quality, computational efficiency, and the aggregated overall score.

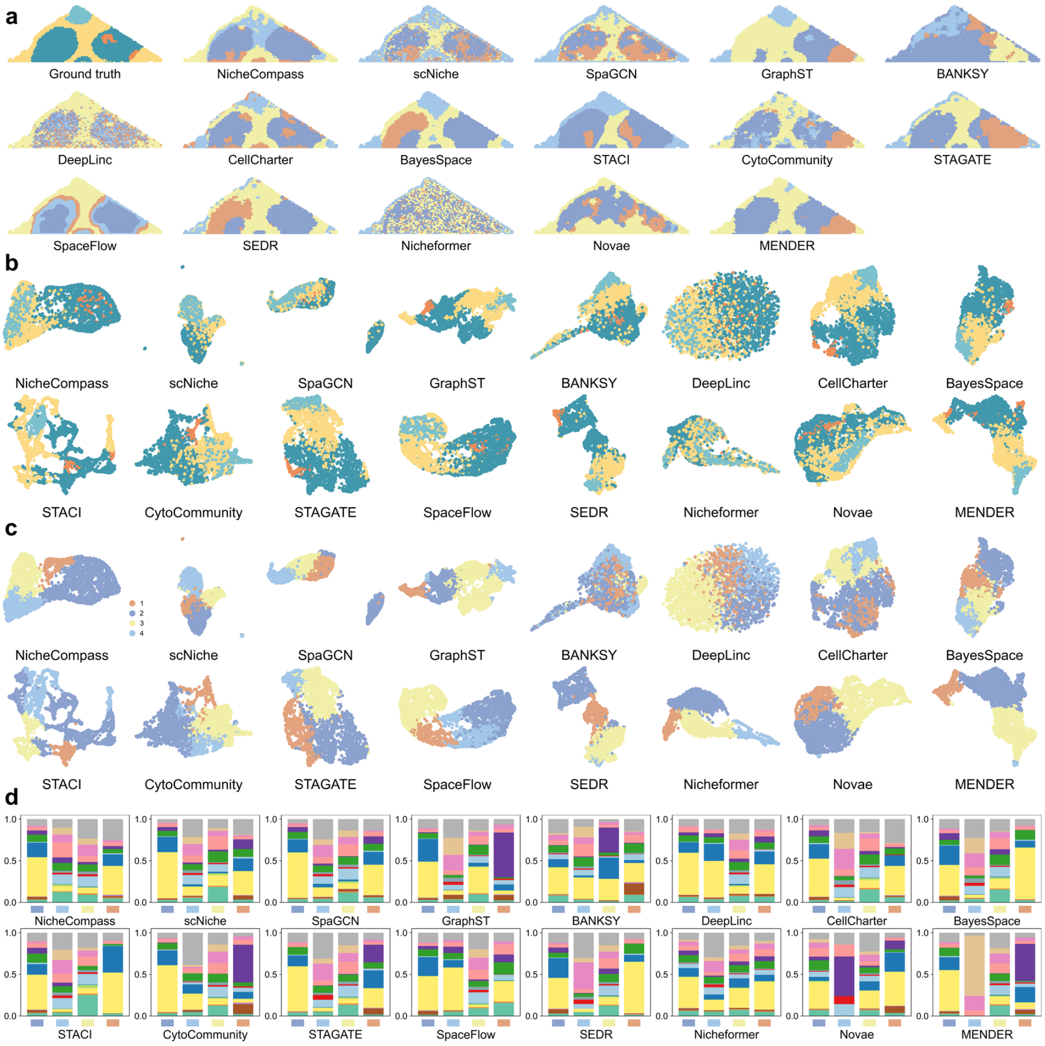


**e**

| Method | Ground Truth Accuracy: ARI | AMI | Homogeneity | Completeness | Macro-F1 | Biological Consistency: Composition Similarity | Spatial Structure: Spatial Connectivity | Embedding Quality: Silhouette Score | Computational Efficiency: Time (s) | Memory (GB) | Aggregate: Total |
|---|---|---|---|---|---|---|---|---|---|---|---|
| **MENDER** | 0.50 | 0.41 | 0.41 | 0.42 | 0.52 | 0.80 | 0.94 | 0.11 | 14.2 | 0.55 | 0.960 |
| **BayesSpace** | 0.35 | 0.44 | 0.52 | 0.39 | 0.49 | 0.79 | 0.93 | 0.06 | 223 | 3.05 | 0.883 |
| **CellCharter** | 0.40 | 0.40 | 0.45 | 0.36 | 0.53 | 0.79 | 0.86 | 0.06 | 8.71 | 1.75 | 0.873 |
| **STAGATE** | 0.36 | 0.37 | 0.42 | 0.33 | 0.48 | 0.78 | 0.91 | 0.07 | 115 | 1.88 | 0.862 |
| **GraphST** | 0.20 | 0.29 | 0.32 | 0.28 | 0.51 | 0.75 | 0.95 | 0.14 | 153 | 1.58 | 0.861 |
| **STACI** | 0.34 | 0.41 | 0.44 | 0.38 | 0.44 | 0.75 | 0.93 | 0.05 | 86.9 | 1.42 | 0.837 |
| **CytoCommunity** | 0.38 | 0.31 | 0.35 | 0.28 | 0.48 | 0.73 | 0.80 | 0.13 | 67.0 | 1.22 | 0.790 |
| **SEDR** | 0.34 | 0.39 | 0.44 | 0.35 | 0.40 | 0.72 | 0.90 | 0.03 | 17.3 | 3.13 | 0.756 |
| **NicheCompass** | 0.43 | 0.41 | 0.45 | 0.38 | 0.42 | 0.75 | 0.88 | -0.18 | 227 | 0.89 | 0.670 |
| **BANKSY** | 0.11 | 0.15 | 0.16 | 0.14 | 0.42 | 0.68 | 0.90 | 0.03 | 25.2 | 1.82 | 0.640 |
| **SpaGCN** | 0.21 | 0.24 | 0.28 | 0.21 | 0.37 | 0.76 | 0.76 | -0.13 | 24.4 | 1.80 | 0.597 |
| **scNiche** | 0.22 | 0.23 | 0.27 | 0.20 | 0.41 | 0.78 | 0.64 | -0.09 | 253 | 3.64 | 0.580 |
| **SpaceFlow** | 0.38 | 0.47 | 0.54 | 0.42 | 0.36 | 0.63 | 0.87 | -0.16 | 16.5 | 2.82 | 0.558 |
| **Novae** | 0.02 | 0.08 | 0.08 | 0.08 | 0.29 | 0.61 | 0.86 | -0.14 | 32.8 | 1.57 | 0.400 |
| **DeepLinc** | 0.17 | 0.22 | 0.25 | 0.19 | 0.30 | 0.66 | 0.50 | -0.03 | 10374 | 0.95 | 0.331 |
| **Nicheformer** | 0.03 | 0.03 | 0.04 | 0.03 | 0.29 | 0.70 | 0.46 | -0.12 | 907 | 14.0 | 0.227 |

**Supplementary Fig. S5.** Benchmarking results on the manually annotated human lymph node dataset using the **pseudo-spot aggregation** strategy. (a) Spatial niche assignments for the ground truth and each method, visualized in tissue coordinates. (b-c) UMAP visualization of method-specific latent embeddings, with cells colored by ground-truth niche labels (b) and by each method's predicted niche labels (c). (d) Cell-type composition of niches for each method, shown as the proportion of annotated cell types within each niche. (e) Multi-criteria quantitative comparison across ground-truth accuracy, biological consistency, spatial structure, embedding quality, computational efficiency, and the aggregated overall score.

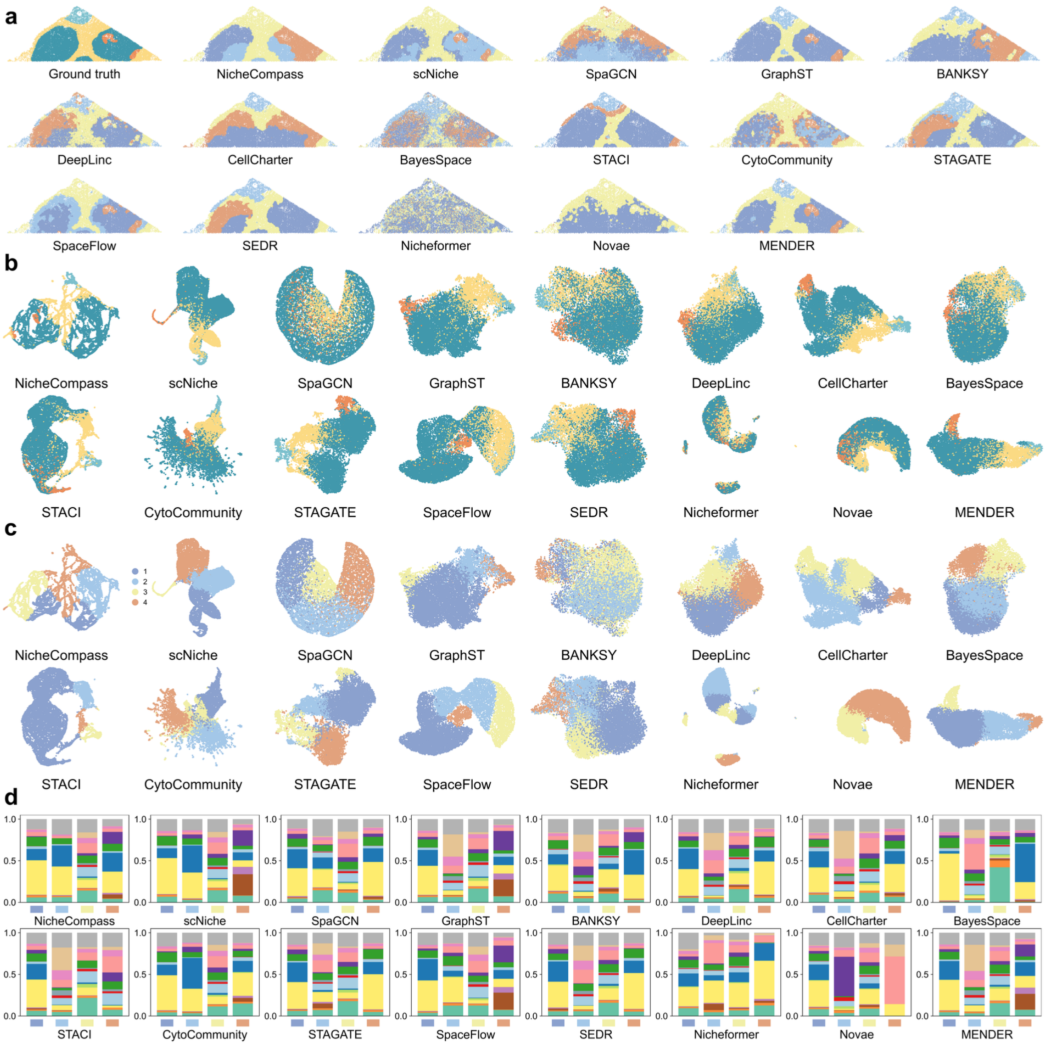


**e**

| | Ground Truth Accuracy | | | | | Biological Consistency | Spatial Structure | Embedding Quality | Computational Efficiency | | Aggregate |
|---|---|---|---|---|---|---|---|---|---|---|---|
| **Method** | ARI | AMI | Homogeneity | Completeness | Macro-F1 | Composition Similarity | Spatial Connectivity | Silhouette Score | Time (s) | Memory (GB) | Total |
| **MENDER** | 0.55 | 0.48 | 0.51 | 0.45 | 0.78 | 0.97 | 0.96 | 0.25 | 112 | 0.67 | 0.966 |
| **GraphST** | 0.61 | 0.51 | 0.51 | 0.51 | 0.77 | 0.96 | 0.97 | 0.09 | 52.5 | 6.40 | 0.841 |
| **CellCharter** | 0.26 | 0.34 | 0.39 | 0.30 | 0.58 | 0.88 | 0.96 | 0.18 | 15.0 | 0.67 | 0.822 |
| **scNiche** | 0.36 | 0.43 | 0.51 | 0.38 | 0.55 | 0.79 | 0.92 | 0.23 | 74.1 | 1.39 | 0.795 |
| **SEDR** | 0.32 | 0.42 | 0.51 | 0.35 | 0.51 | 0.83 | 0.97 | 0.17 | 86.0 | 3.93 | 0.781 |
| **SpaceFlow** | 0.43 | 0.47 | 0.55 | 0.41 | 0.55 | 0.81 | 0.94 | 0.16 | 985 | 1.81 | 0.751 |
| **STAGATE** | 0.34 | 0.37 | 0.45 | 0.31 | 0.48 | 0.82 | 0.93 | 0.19 | 610 | 2.32 | 0.747 |
| **STACI** | 0.50 | 0.44 | 0.43 | 0.45 | 0.55 | 0.82 | 0.98 | -0.05 | 323 | 2.56 | 0.678 |
| **CytoCommunity** | 0.26 | 0.33 | 0.42 | 0.28 | 0.40 | 0.71 | 0.83 | 0.26 | 1425 | 2.94 | 0.612 |
| **NicheCompass** | 0.25 | 0.36 | 0.45 | 0.29 | 0.39 | 0.72 | 0.98 | 0.08 | 1930 | 0.78 | 0.597 |
| **BANKSY** | 0.15 | 0.18 | 0.21 | 0.16 | 0.43 | 0.82 | 0.95 | 0.11 | 112 | 11.9 | 0.587 |
| **Novae** | 0.26 | 0.19 | 0.17 | 0.21 | 0.34 | 0.58 | 0.94 | 0.09 | 247 | 0.67 | 0.545 |
| **SpaGCN** | 0.14 | 0.19 | 0.25 | 0.16 | 0.30 | 0.73 | 0.84 | -0.11 | 101 | 2.22 | 0.445 |
| **DeepLinc** | 0.30 | 0.31 | 0.38 | 0.27 | 0.50 | 0.84 | 0.85 | -0.08 | 2372 | 11.9 | 0.394 |
| **BayesSpace** | 0.21 | 0.22 | 0.28 | 0.19 | 0.36 | 0.69 | 0.58 | 0.02 | 1039 | 0.67 | 0.359 |
| **Nicheformer** | 0.02 | 0.01 | 0.01 | 0.01 | 0.28 | 0.67 | 0.57 | 0.05 | 2372 | 2.82 | 0.211 |

**Supplementary Fig. S6.** Benchmarking results on the manually annotated human lymph node dataset using the **core-cell-type** strategy. (a) Spatial niche assignments for the ground truth and each method, visualized in tissue coordinates. (b-c) UMAP visualization of method-specific latent embeddings, with cells colored by ground-truth niche labels (b) and by each method's predicted niche labels (c). (d) Cell-type composition of niches for each method, shown as the proportion of annotated cell types within each niche. (e) Multi-criteria quantitative comparison across ground-truth accuracy, biological consistency, spatial structure, embedding quality, computational efficiency, and the aggregated overall score.

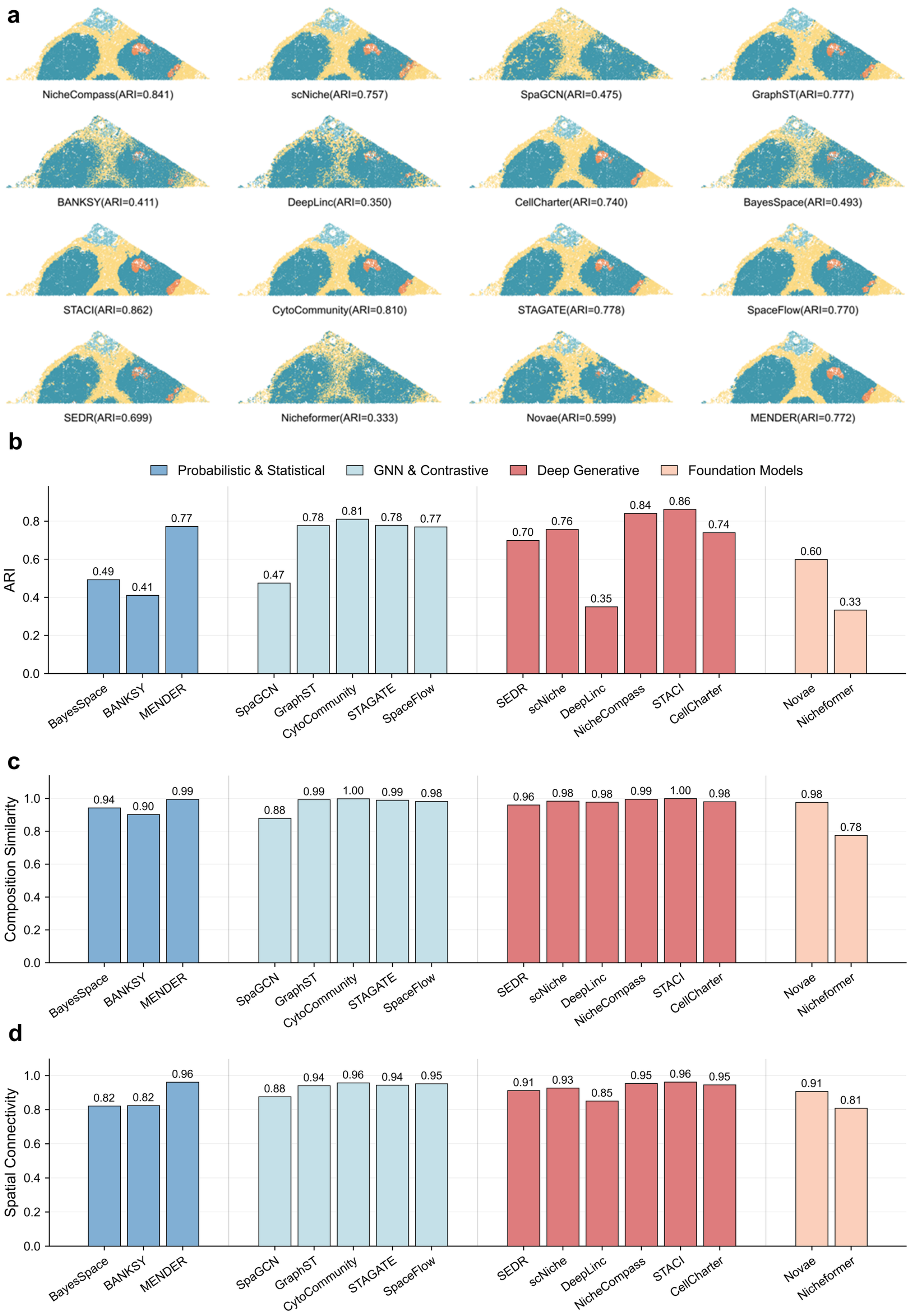


**Supplementary Fig. S7.** Embedding-supervised enhancement of niche assignments on the manually annotated human lymph node dataset. (a) Spatial niche maps after supervised refinement of each method's latent embedding using ground-truth niche labels. For each method, a kNN classifier was trained on the method-specific embedding with stratified cross-validation to generate out-of-fold niche predictions for all cells, which were then visualized in tissue coordinates; ARI values are reported for the refined results. (b) Adjusted Rand Index (ARI) of the embedding-supervised niche assignments across methods, with bar colors indicating method categories (Probabilistic & Statistical, GNN & Contrastive, Deep Generative, and Foundation Models). (c) Composition Similarity of the refined niche assignments. (d) Spatial Connectivity of the refined niche assignments.

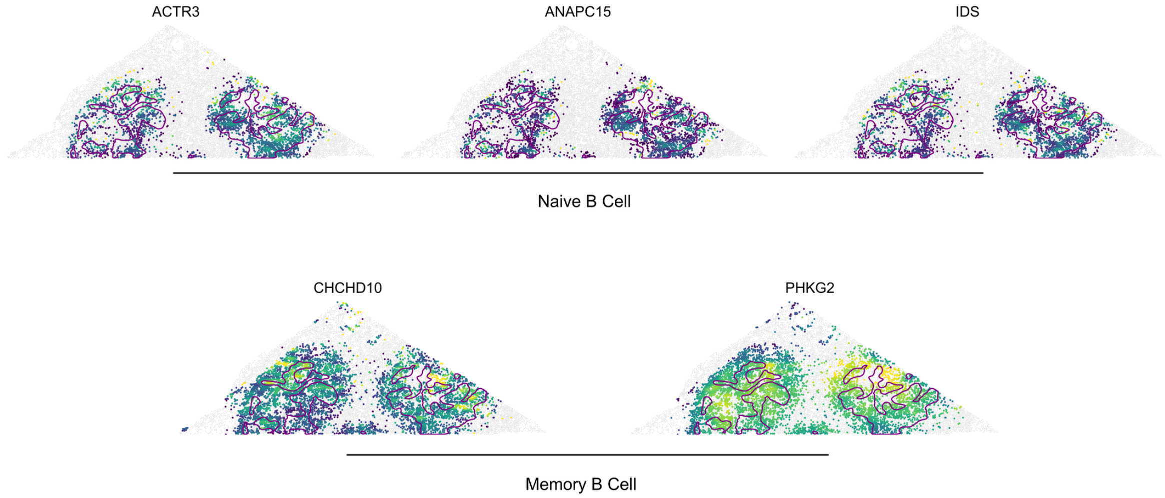


**Supplementary Fig. S8.** Expression of representative marker genes within the B-cell maturation zone: ACTR3, ANAPC15 and IDS (Naive B-associated), and CHCHD10, PHKG2 (Memory B-associated).

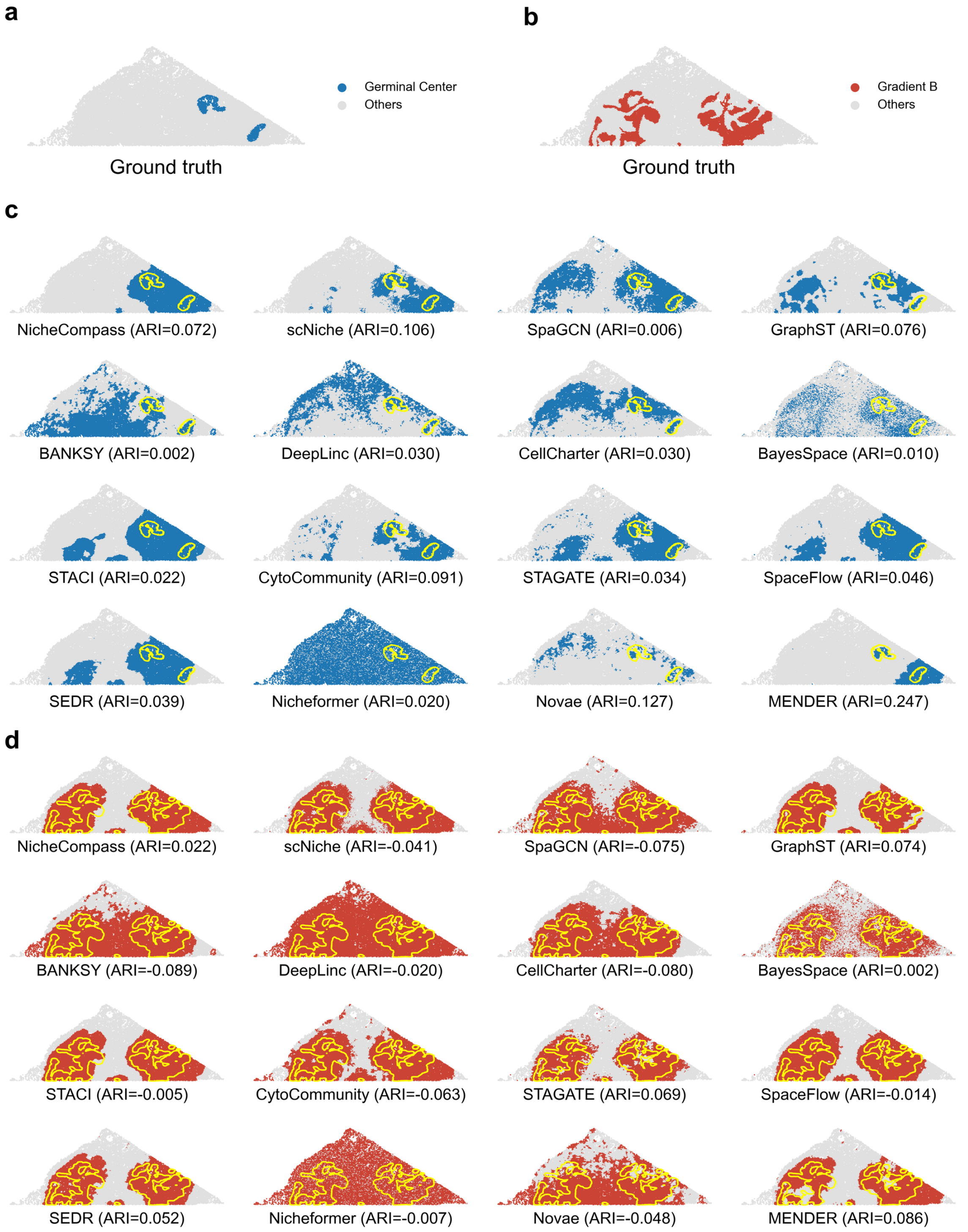


**Supplementary Fig. S9.** Performance of niche-detection methods in identifying an island niche (Germinal Center) and a gradient niche (B-cell maturation zone) in the annotated human lymph node dataset (n = 19,718 cells). (a) Ground-truth annotation for the island-like Germinal Center (GC) niche. (b) Ground-truth annotation for the B-cell maturation zone niche. (c) Spatial predictions from 16 methods for GC. (d) Spatial predictions from 16 methods for B-cell maturation zone. For each method, clusters with spatial overlap exceeding a predefined threshold with the corresponding ground-truth region were merged and reported as the method's predicted niche region (GC: overlap threshold = 0.30; B-cell maturation zone: overlap threshold = 0.10). Adjusted Rand Index (ARI) values are shown for each method. Yellow contours delineate the ground-truth niche boundaries; gray indicates all other cells.

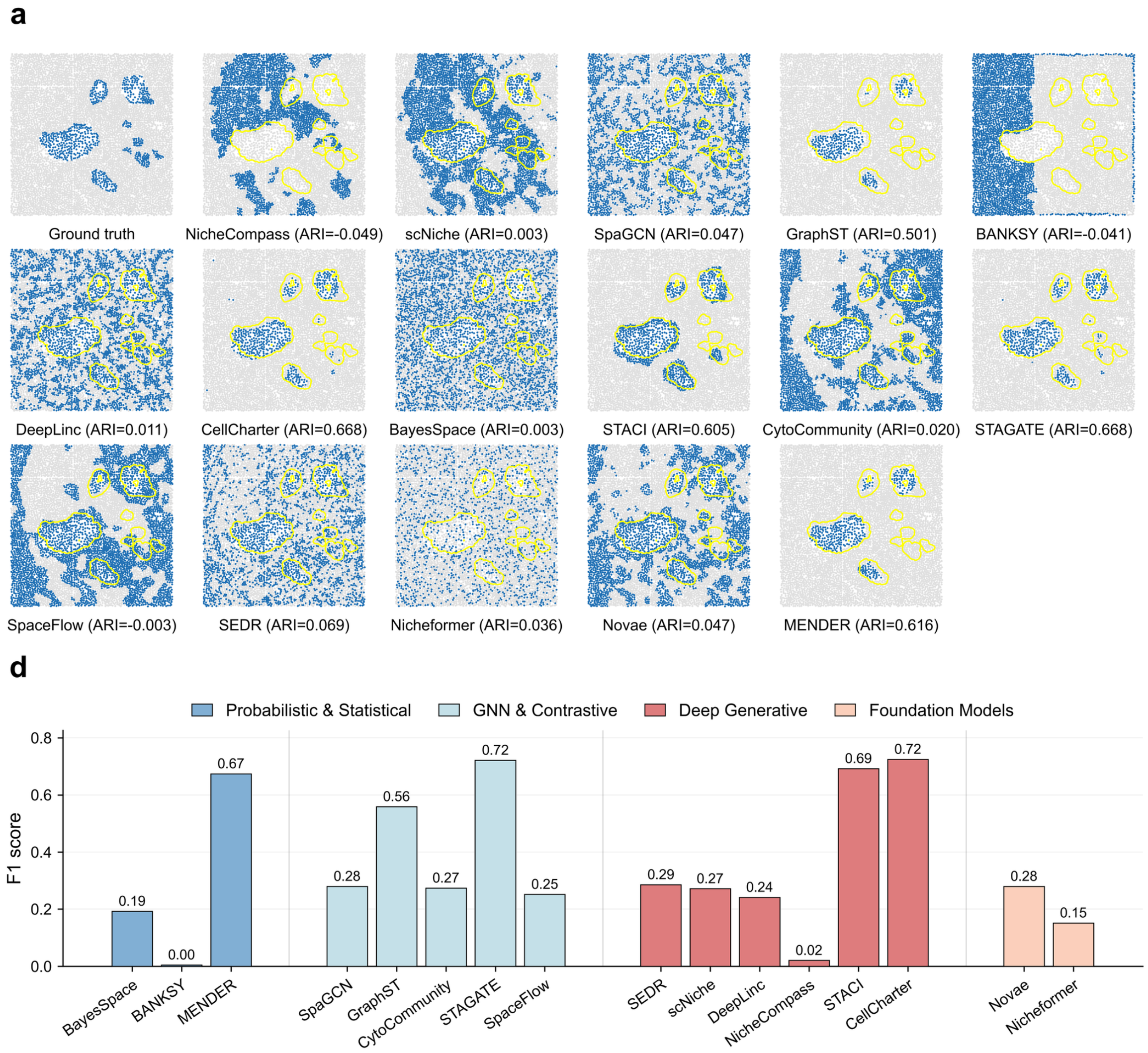


**Supplementary Fig. S10.** Detection of an island-like Germinal Center niche in an additional B cell zone region (n = 7,319 cells) from the original CosMx human lymph node dataset. (a) Spatial maps comparing the ground-truth Germinal Center (GC) island annotation with predictions from 16 niche-detection methods. Predicted GC regions were obtained by merging all clusters whose overlap with the GC ground-truth region exceeded a predefined threshold; yellow contours delineate the ground-truth GC islands and gray indicates non-GC cells. Method names and ARI values are shown for each panel. (b) F1 scores for GC island detection across methods. Bar colors denote method categories (Probabilistic & Statistical, GNN & Contrastive, Deep Generative, and Foundation Models).

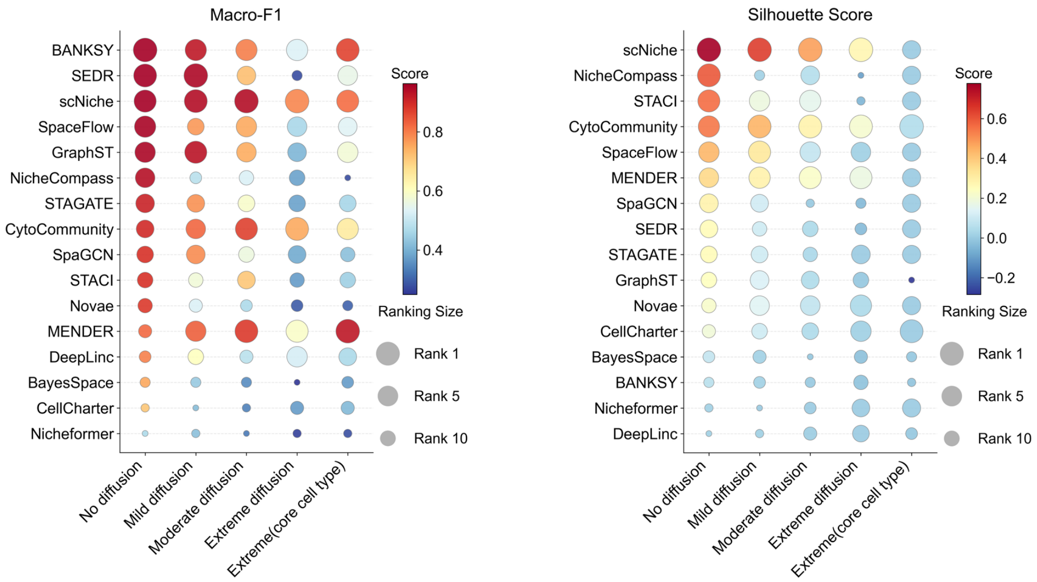


**Supplementary Fig. S11.** Bubble-matrix summary of additional evaluation metrics for simulated lymph node benchmarking across cell-type compositional diffusion settings. This figure complements panel (f) of the main simulation benchmark by reporting the remaining two metrics: Macro-F1 (left) and Silhouette Score (right), evaluated under no, mild, moderate, and extreme diffusion, as well as an additional condition applying the core-cell-type refinement strategy under extreme diffusion. Bubble color encodes the metric value and bubble size indicates the relative rank within each condition; methods are ordered by performance in the no-diffusion setting.

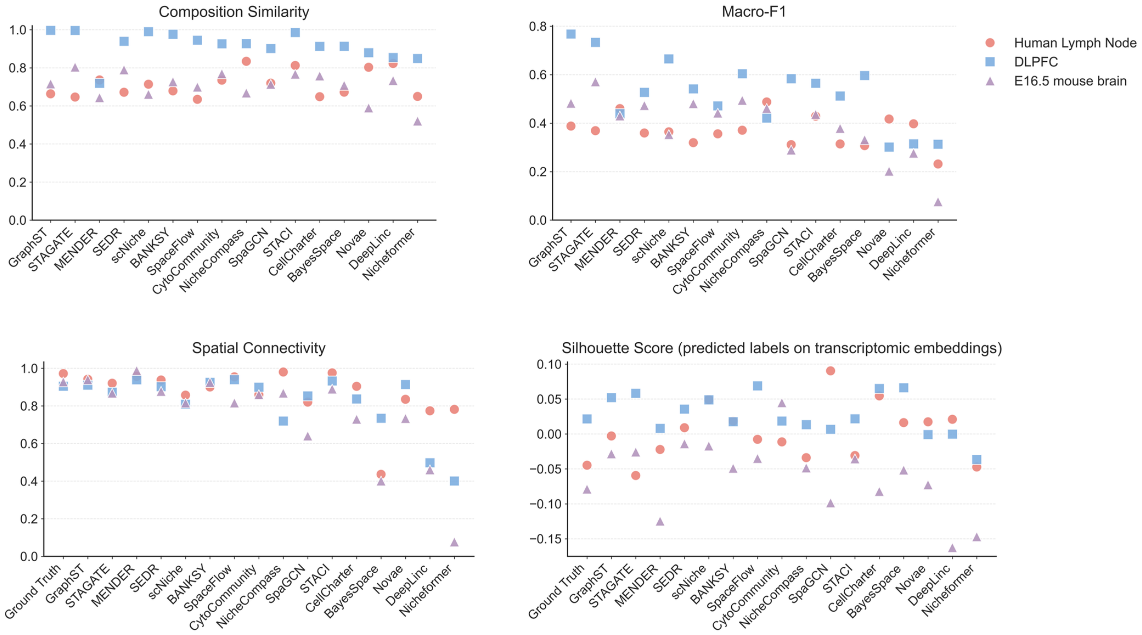


**Supplementary Fig. S12.** Cross-dataset comparison of additional evaluation metrics for niche/anatomical-domain detection methods. This figure complements panel (e) of the main cross-technology benchmark by reporting the remaining metrics across the manually annotated human lymph node reference, DLPFC slice 151673, and the MOSTA E16.5 mouse brain dataset (marker shape/color indicates dataset). Shown are Composition Similarity (top left), Macro-F1 (top right), Spatial Connectivity (bottom left), and Silhouette Index (SI) score (bottom right). The SI score was computed on a shared transcriptomic UMAP for each dataset by calculating silhouette scores using (i) the ground-truth labels and (ii) each method's predicted labels. Methods are ordered consistently across panels to facilitate comparison.

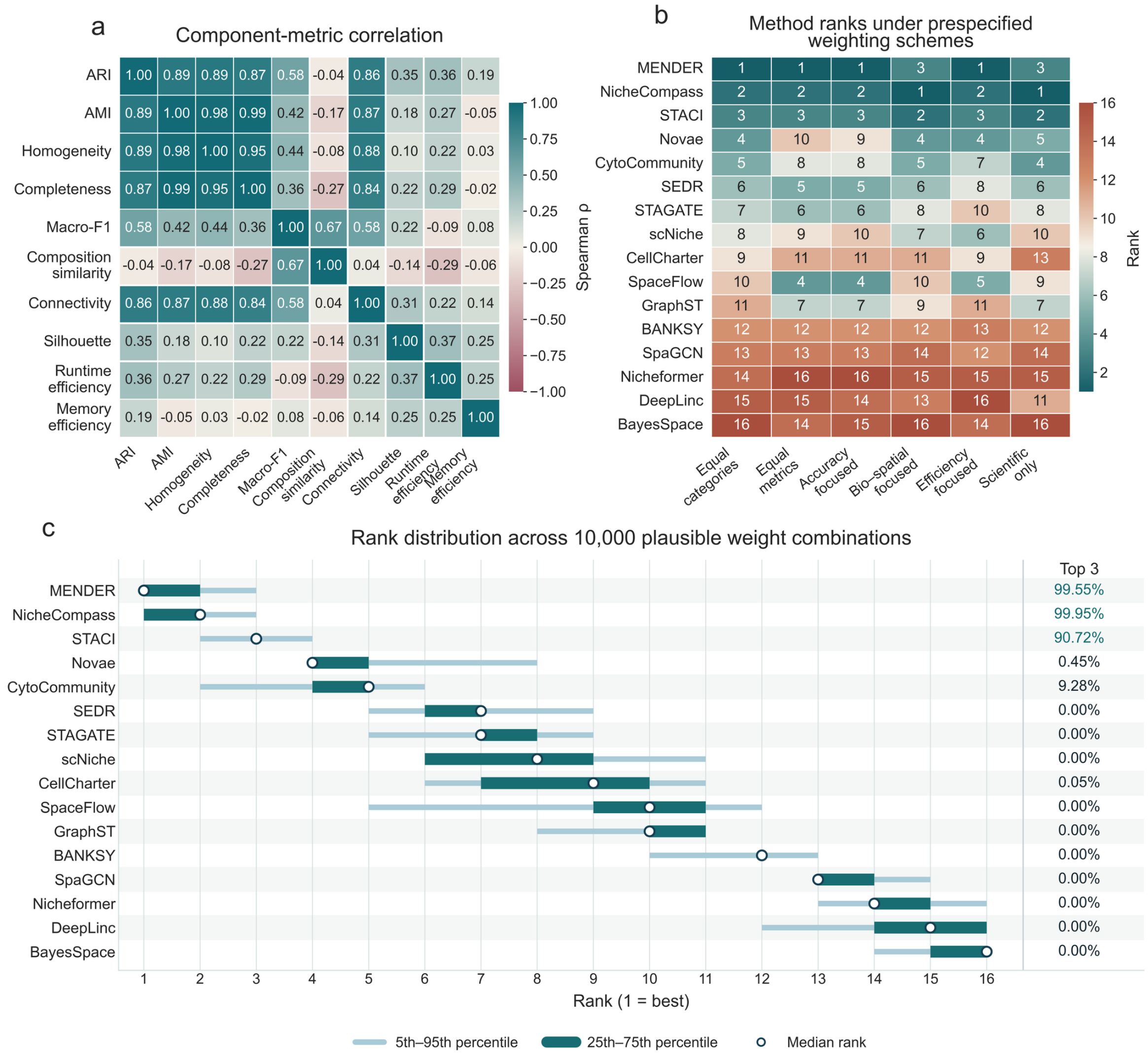


**Supplementary Fig. S13.** Robustness of aggregate performance rankings across alternative weighting schemes. (a) Spearman correlation matrix of the ten component metrics across the 16 methods in the primary lymph-node benchmark. Runtime and memory are represented as reverse-normalized efficiency measures, such that higher values indicate better performance. (b) Method ranks under six prespecified weighting schemes representing different evaluation priorities. Lower rank indicates better aggregate performance. (c) Rank distributions across 10,000 plausible combinations of the five category weights. Light bars indicate the 5th–95th percentile range, dark bars the 25th–75th percentile range, and open circles the median rank; values at right indicate the percentage of weight combinations in which each method ranked among the top three.

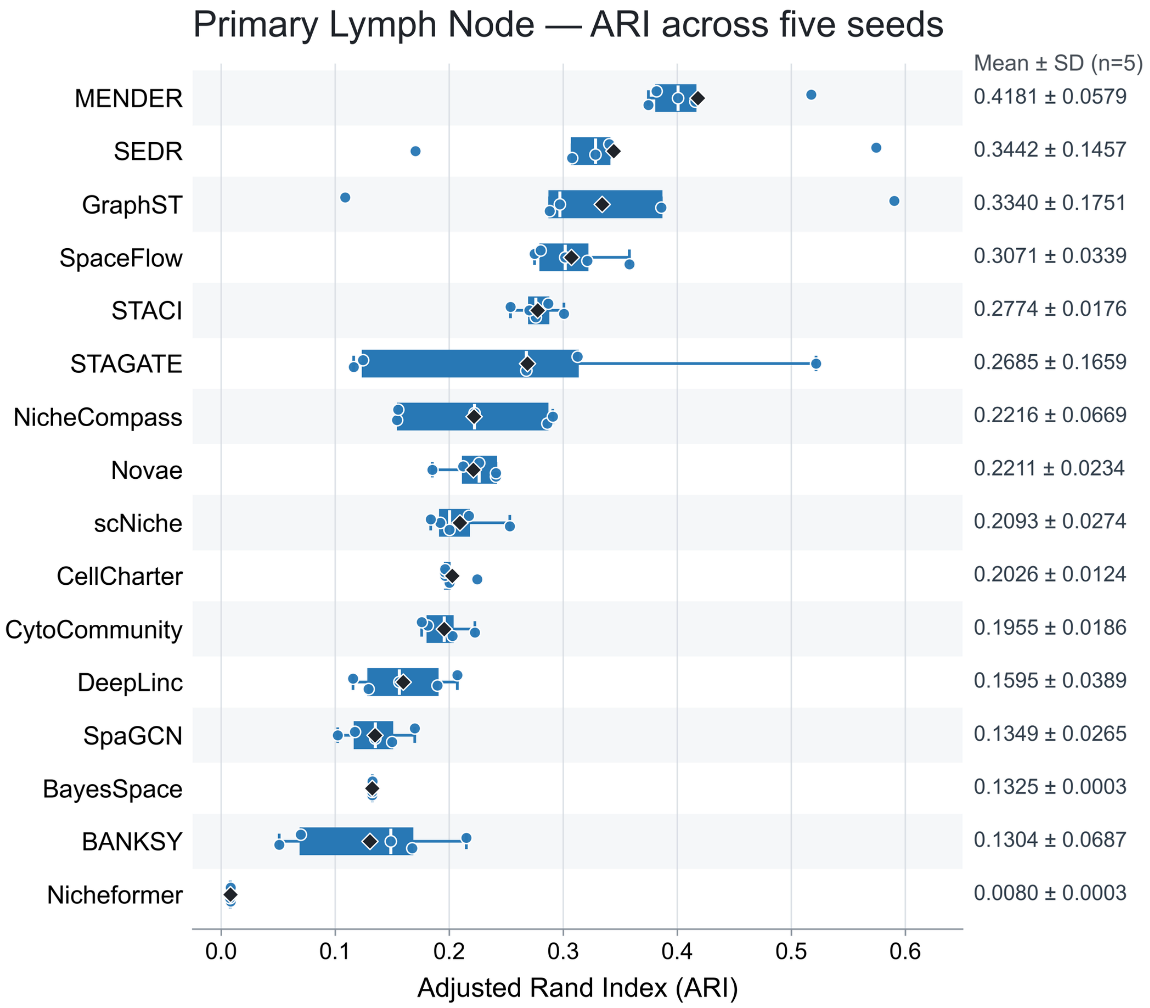


**Supplementary Fig. S14.** Random-seed reproducibility of ARI in the primary human lymph-node benchmark. Adjusted Rand Index (ARI) distributions for the 16 evaluated methods across five prespecified random seeds. Input data, preprocessing, target domain number, and method-specific hyperparameters were held fixed across repeats, with only the random seed varied. Individual points denote the five runs, boxplots summarize the corresponding distributions, and diamonds indicate the mean ARI; method-wise mean ± SD is reported at right. Methods are ordered by mean ARI. Complete run-level results and method-wise summary statistics are provided in Supplementary Tables S2 and S3, respectively.

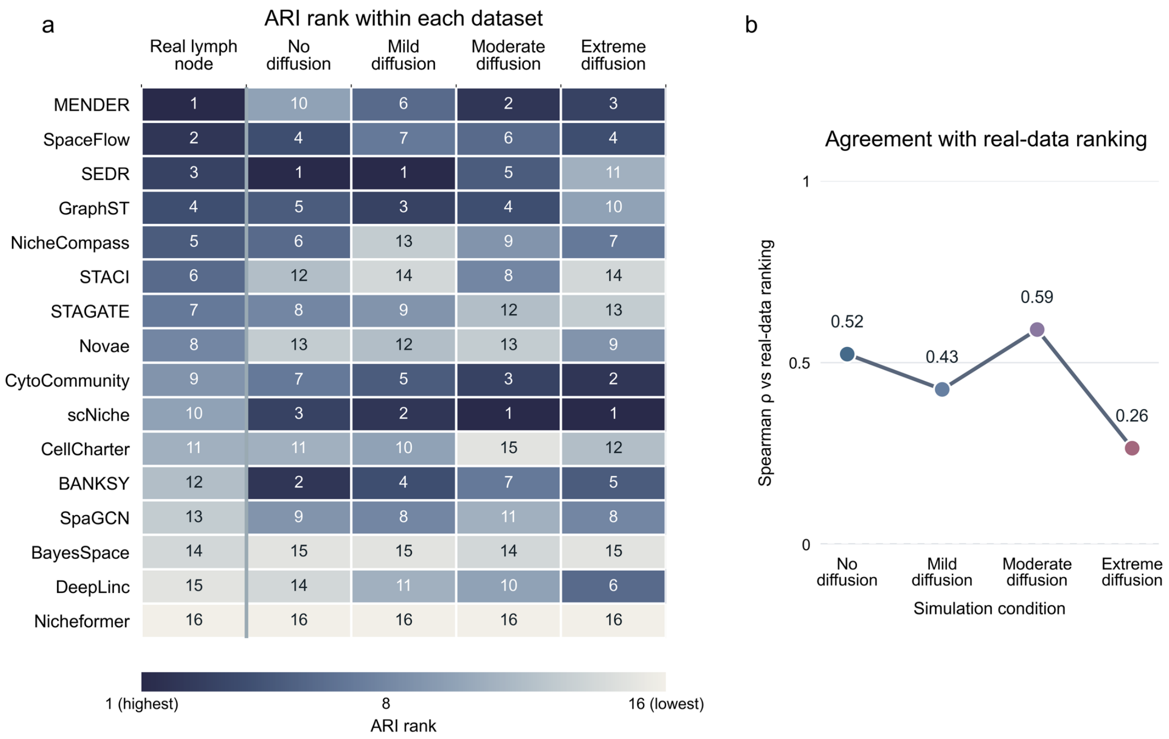


| | Real lymph node | No diffusion | Mild diffusion | Moderate diffusion | Extreme diffusion |
|---|---|---|---|---|---|
| MENDER | 1 | 10 | 6 | 2 | 3 |
| SpaceFlow | 2 | 4 | 7 | 6 | 4 |
| SEDR | 3 | 1 | 1 | 5 | 11 |
| GraphST | 4 | 5 | 3 | 4 | 10 |
| NicheCompass | 5 | 6 | 13 | 9 | 7 |
| STACI | 6 | 12 | 14 | 8 | 14 |
| STAGATE | 7 | 8 | 9 | 12 | 13 |
| Novae | 8 | 13 | 12 | 13 | 9 |
| CytoCommunity | 9 | 7 | 5 | 3 | 2 |
| scNiche | 10 | 3 | 2 | 1 | 1 |
| CellCharter | 11 | 11 | 10 | 15 | 12 |
| BANKSY | 12 | 2 | 4 | 7 | 5 |
| SpaGCN | 13 | 9 | 8 | 11 | 8 |
| BayesSpace | 14 | 15 | 15 | 14 | 15 |
| DeepLinc | 15 | 14 | 11 | 10 | 6 |
| Nicheformer | 16 | 16 | 16 | 16 | 16 |

**Supplementary Fig. S15.** Comparison of method rankings between the real and simulated lymph-node benchmarks. (a) ARI ranks of the 16 methods in the primary human lymph-node benchmark and under no, mild, moderate, and extreme cell-type compositional diffusion. Methods were ranked independently within each dataset, with rank 1 indicating the highest ARI and rank 16 the lowest. (b) Spearman rank correlation ($\rho$) between the method ranking in each simulated diffusion condition and that in the real lymph-node benchmark. Higher values indicate greater concordance with the real-data ranking.

a

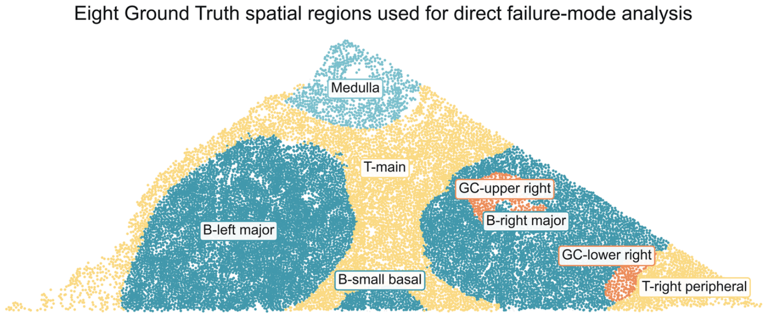


b

Class-level niche recovery and structural failure modes

| Method | B-cell zone (3 GT regions) | T-cell zone (2 GT regions) | Medulla (1 GT region) | Germinal center (2 GT regions) |
|---|---|---|---|---|
| NicheCompass | 2/3 recovered Split → 2 labels | 1/2 recovered Split → 2 labels | 0/1 recovered Hidden with T | 0/2 recovered Hidden with B |
| scNiche | 3/3 recovered Split → 3 labels | 1/2 recovered Other hidden with B | 0/1 recovered Hidden with T | 0/2 recovered Hidden with B/T |
| SpaGCN | 0/3 recovered Split → 3 labels | 0/2 recovered Split → 2 labels | 0/1 recovered Hidden with T | 0/2 recovered Hidden with B/T |
| GraphST | 1/3 recovered Split → 2 labels | 2/2 recovered | 0/1 recovered Hidden with B/T | 0/2 recovered Hidden with B |
| BANKSY | 0/3 recovered Split → 2 labels | 1/2 recovered Split → 2 labels | 0/1 recovered Split → 2 labels | 0/2 recovered Hidden with B/T |
| DeepLinc | 1/3 recovered Split → 3 labels | 0/2 recovered Split → 2 labels | 0/1 recovered Hidden with B/T | 0/2 recovered Hidden with B |
| CellCharter | 0/3 recovered Split → 2 labels | 1/2 recovered Split → 3 labels | 0/1 recovered Hidden with T | 1/2 recovered Lower-right GC Other hidden with B/T |
| BayesSpace | 0/3 recovered Split → 2 labels | 0/2 recovered Split → 3 labels | 0/1 recovered Hidden with T | 0/2 recovered Not recovered: 2 |
| STACI | 3/3 recovered Split → 2 labels | 0/2 recovered Split → 2 labels | 0/1 recovered Hidden with T | 0/2 recovered Hidden with B |
| CytoCommunity | 1/3 recovered Split → 3 labels | 1/2 recovered Split → 2 labels | 0/1 recovered Hidden with T | 0/2 recovered Hidden with B/T |
| STAGATE | 2/3 recovered Split → 2 labels | 2/2 recovered | 0/1 recovered Hidden with B/T | 0/2 recovered Hidden with B |
| SpaceFlow | 2/3 recovered Split → 2 labels | 2/2 recovered | 0/1 recovered Hidden with T | 0/2 recovered Hidden with B |
| SEDR | 2/3 recovered Split → 2 labels | 2/2 recovered Split → 2 labels | 0/1 recovered Hidden with B/T | 0/2 recovered Hidden with B |
| Nicheformer | 0/3 recovered Hidden with T | 0/2 recovered Hidden with B | 0/1 recovered Hidden with B/T | 0/2 recovered Hidden with B/T |
| Novae | 0/3 recovered Partial: 2 | 0/2 recovered Split → 3 labels | 0/1 recovered Hidden with T | 0/2 recovered Hidden with B/T |
| MENDER | 3/3 recovered | 1/2 recovered Other hidden with B | 0/1 recovered Hidden with B/T | 1/2 recovered Upper-right GC Other hidden with B/T |

Recovered | Partial recovery | Split | Merged/hidden | Partial | Missed/fragmented

Split: ≥2 labels, each with ≥15% Ground Truth coverage and ≥15% label purity.

c

Region-level niche recovery and failure modes

| Method | B-cell zone | | | T-cell zone | | Medulla | Germinal center | |
|---|---|---|---|---|---|---|---|---|
| | Left major | Right major | Small basal | Main | Right peripheral | Medulla | Upper right | Lower right |
| NicheCompass | R 0.93 | R 0.84 | F 0.40 | R 0.68 | H 0.23 | H 0.50 | H 0.11 | H 0.07 |
| scNiche | R 0.71 | R 0.66 | R 0.79 | R 0.79 | H 0.29 | H 0.28 | H 0.13 | H 0.09 |
| SpaGCN | F 0.44 | F 0.60 | H 0.16 | F 0.64 | F 0.41 | H 0.36 | H 0.19 | M 0.36 |
| GraphST | F 0.45 | F 0.55 | R 0.72 | R 0.80 | R 0.73 | H 0.22 | H 0.29 | P 0.57 |
| BANKSY | H 0.51 | F 0.64 | H 0.08 | F 0.45 | R 0.80 | F 0.45 | H 0.06 | H 0.61 |
| DeepLinc | F 0.40 | R 0.76 | M 0.28 | F 0.53 | P 0.52 | H 0.20 | H 0.43 | P 0.51 |
| CellCharter | H 0.46 | H 0.44 | H 0.09 | F 0.45 | R 0.87 | H 0.48 | H 0.09 | R 0.78 |
| BayesSpace | M 0.28 | M 0.39 | M 0.35 | H 0.30 | M 0.22 | H 0.44 | F 0.29 | M 0.17 |
| STACI | R 0.76 | R 0.86 | R 0.77 | F 0.62 | M 0.38 | H 0.49 | H 0.11 | H 0.06 |
| CytoCommunity | R 0.68 | F 0.47 | H 0.48 | R 0.67 | H 0.41 | H 0.32 | H 0.37 | H 0.14 |
| STAGATE | R 0.68 | R 0.85 | P 0.61 | R 0.79 | R 0.87 | H 0.22 | H 0.12 | H 0.08 |
| SpaceFlow | R 0.82 | R 0.88 | F 0.46 | R 0.82 | R 0.88 | H 0.28 | H 0.13 | H 0.08 |
| SEDR | R 0.77 | R 0.90 | P 0.61 | R 0.78 | R 0.92 | H 0.23 | H 0.12 | H 0.07 |
| Nicheformer | H 0.51 | H 0.39 | H 0.04 | H 0.39 | H 0.09 | H 0.08 | H 0.03 | M 0.06 |
| Novae | P 0.52 | P 0.46 | M 0.08 | F 0.33 | F 0.43 | H 0.47 | F 0.40 | H 0.35 |
| MENDER | R 0.83 | R 0.78 | R 0.72 | R 0.76 | H 0.57 | H 0.24 | R 0.71 | H 0.20 |

R Recovered | P Partial | F Fragmented | H Merged/hidden | M Missed

Recovered: recall and precision ≥0.50; F1 ≥0.65. Substantial overlap: ≥15%.

d

| | B-cell zone | T-cell zone | Medulla | Germinal center |
|---|---|---|---|---|
| NicheCompass | Split ×2 0.972 | Split ×2 0.916 | NA Merged/hidden | NA Merged/hidden |
| scNiche | Split ×3 0.901 | Single 0.943 | NA Merged/hidden | NA Merged/hidden |
| SpaGCN | Split ×3 0.924 | Split ×2 0.892 | NA Merged/hidden | NA Merged/hidden |
| GraphST | Split ×2 0.989 | Single 0.960 | NA Merged/hidden | NA Merged/hidden |
| BANKSY | Split ×2 0.937 | Split ×2 0.869 | Split ×2 0.605 | NA Merged/hidden |
| DeepLinc | Split ×3 0.901 | Split ×2 0.862 | NA Merged/hidden | NA Merged/hidden |
| CellCharter | Split ×2 0.982 | Split ×3 0.760 | NA Merged/hidden | NA No match |
| BayesSpace | Split ×2 0.871 | Split ×3 0.684 | NA Merged/hidden | NA Missed |
| STACI | Split ×2 0.984 | Split ×2 0.930 | NA Merged/hidden | NA Merged/hidden |
| CytoCommunity | Split ×3 0.924 | Split ×2 0.911 | NA Merged/hidden | NA Merged/hidden |
| STAGATE | Split ×2 0.982 | Single 0.986 | NA Merged/hidden | NA Merged/hidden |
| SpaceFlow | Split ×2 0.975 | Single 0.953 | NA Merged/hidden | NA Merged/hidden |
| SEDR | Split ×2 0.983 | Split ×2 0.873 | NA Merged/hidden | NA Merged/hidden |
| Nicheformer | NA Merged/hidden | NA Merged/hidden | NA Merged/hidden | NA Merged/hidden |
| Novae | NA No recovery | Split ×3 0.820 | NA Merged/hidden | NA Merged/hidden |
| MENDER | Single 0.994 | Single 0.956 | NA Merged/hidden | Single 0.611 |

Single-domain recovery | Split recovery | Not evaluable (NA)

**Supplementary Fig. S16.** Decomposition of niche-recovery failure modes and conditional composition fidelity. (a) Spatial decomposition of the four manually annotated reference niches into eight connected regions used for direct failure-mode analysis, comprising three B-cell-zone regions, two T-cell-zone regions, one Medulla region, and two germinal-center regions. (b) Class-level recovery summary across the 16 methods, indicating the number of constituent reference regions recovered and the corresponding structural outcome, including recovery, partial recovery, splitting, merging or hiding, and missing or fragmentation. (c) Region-level assessment of the eight connected reference regions. Each entry reports the recovery status together with the best spatial-component F1 score, enabling localization of specific regional failures. (d) Cell-type composition similarity evaluated conditional on a reliable spatial correspondence. Single-domain and split recoveries are assigned composition-similarity values, whereas niches without an identifiable correspondence because of merging, hiding, missing, or insufficient overlap are reported as not evaluable (NA) rather than assigned an artificial similarity score.

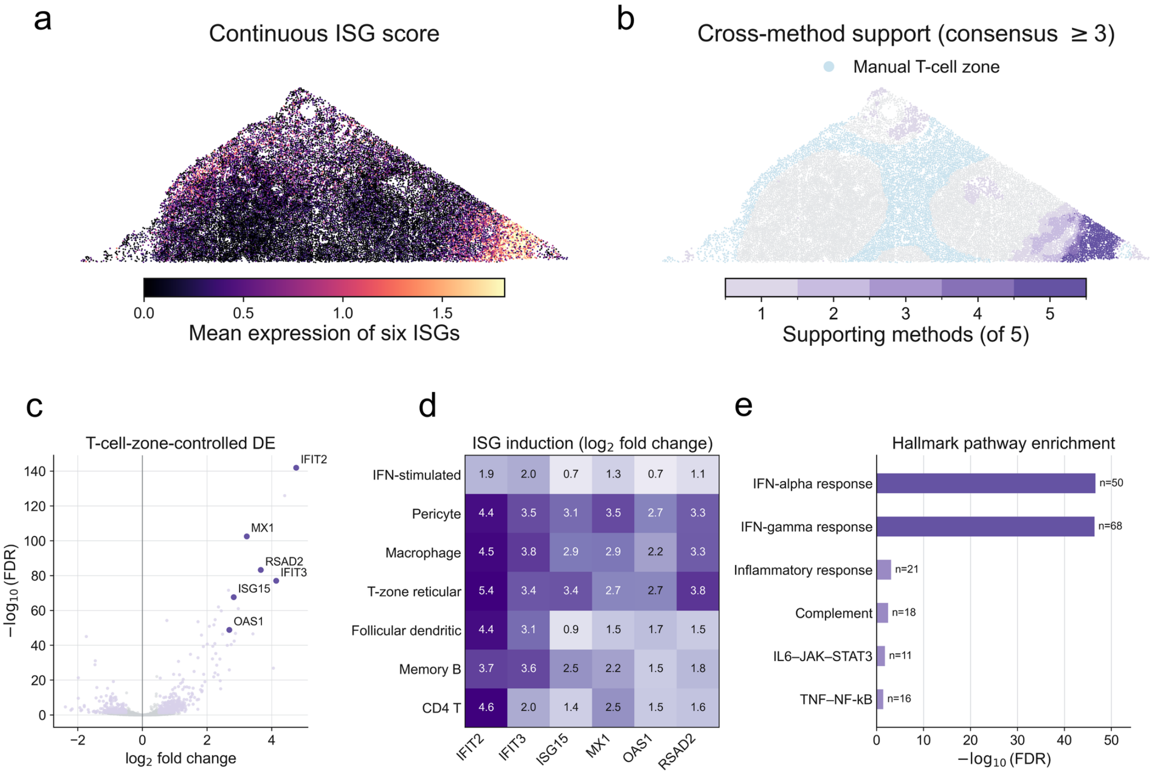


**Supplementary Fig. S17.** Biological characterization of a method-defined interferon-responsive microenvironment in the human lymph node. (a) Continuous spatial interferon-stimulated-gene (ISG) score, calculated from the mean expression of *IFIT2*, *IFIT3*, *ISG15*, *MX1*, *OAS1*, and *RSAD2*. (b) Cell-level cross-method support for the ISG-associated domains identified by GraphST, BANKSY, CellCharter, SpaceFlow, and MENDER. The consensus core was defined as cells supported by at least three of the five methods; 91.5% of consensus-core cells were located within the manually annotated T-cell zone and 81.1% belonged to the highest decile of the continuous ISG score. (c) Differential expression between consensus-core cells and the remaining cells within the manually annotated T-cell zone, controlling for baseline transcriptional differences between anatomical compartments. (d) Cell-type-resolved induction of the six representative ISGs, showing that the response extends across multiple immune and stromal populations rather than being restricted to a single annotated cell type. (e) Hallmark pathway enrichment of the T-cell-zone-controlled differential-expression signature. Interferon-alpha and interferon-gamma response programs were the dominant enriched pathways; the interferon-gamma signature is interpreted as a transcriptional response program and does not by itself identify the upstream cytokine source.

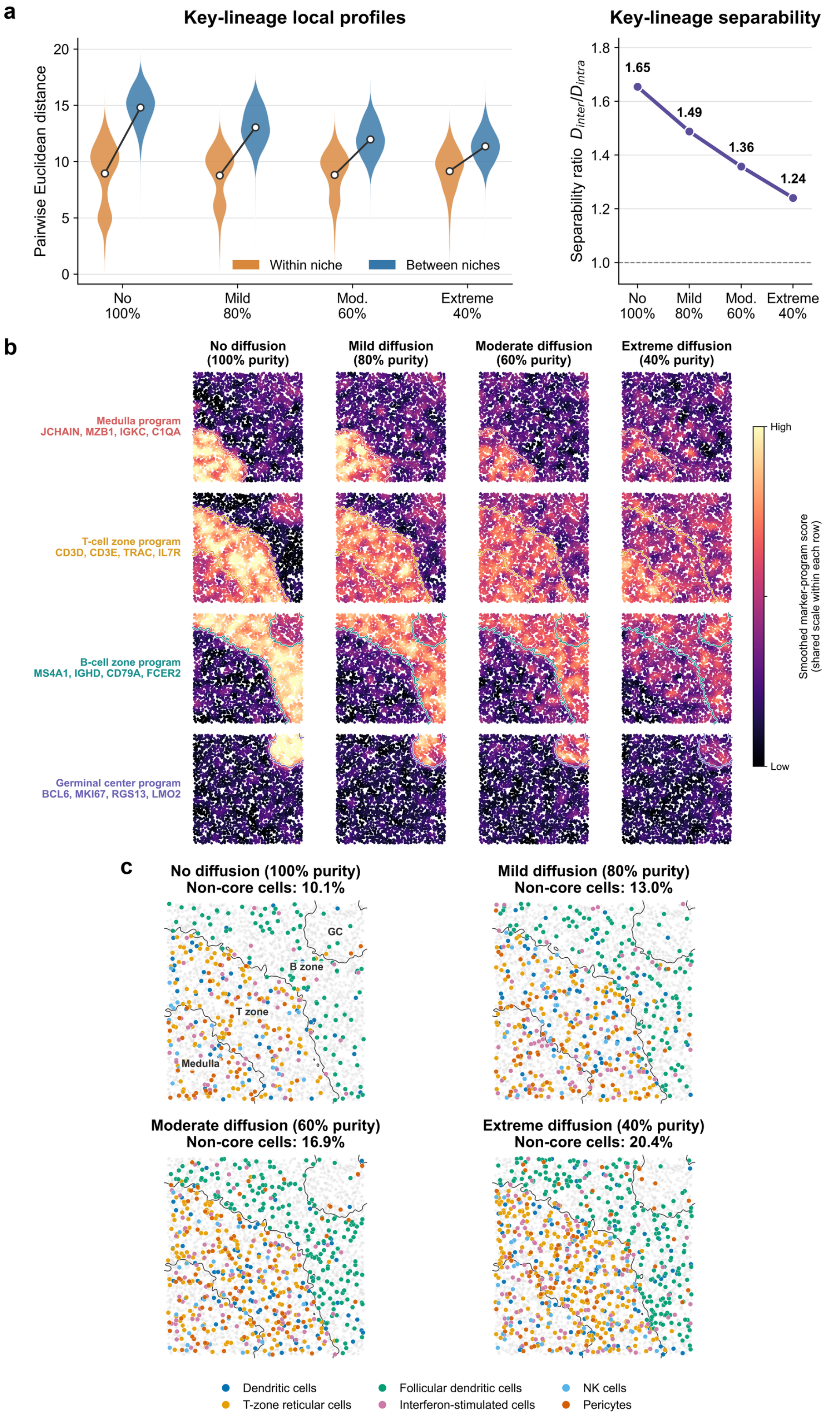


**Supplementary Fig. S18.** Cell-type compositional diffusion reduces molecular separability and increases non-core cell mixing in simulated lymph-node niches. (a) Within- and between-niche Euclidean distances between Gaussian-smoothed key-lineage expression profiles across increasing diffusion, with the corresponding separability ratio $D_{\mathrm{inter}}/D_{\mathrm{intra}}$. (b) Spatial scores of representative niche-marker programs, showing progressive loss of spatial confinement and niche contrast with increasing diffusion. (c) Spatial distributions of representative non-core cell populations. Percentages indicate their combined fraction, which increases and becomes more dispersed across niche boundaries with diffusion.

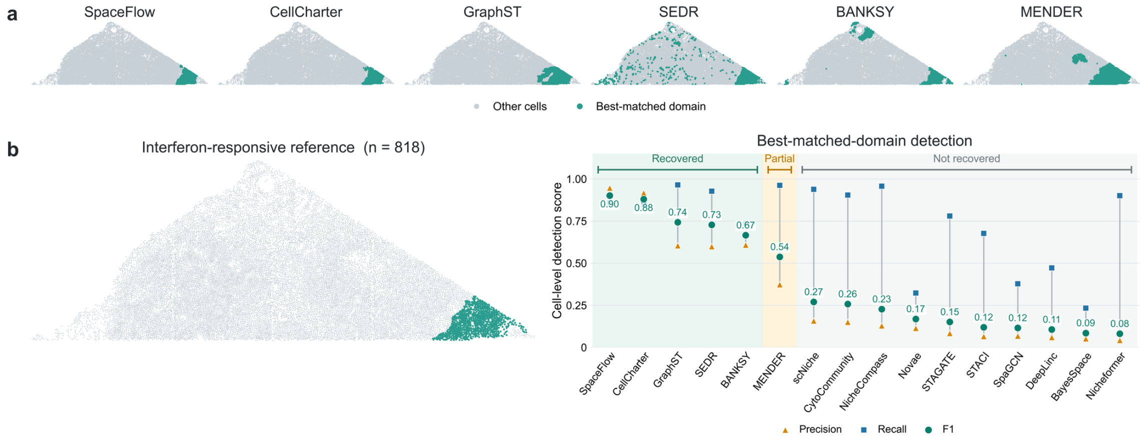


**Supplementary Fig. S19.** Benchmarking results of interferon-stimulation segmentation. (a) Spatial distributions of the best-matched domains identified by GraphST, BANKSY, CellCharter,SEDR, SpaceFlow, and MENDER, with all remaining cells shown in gray. (b) Binary interferon-responsive reference comprising 818 cells (left) and quantitative agreement of the best-matched domain from each of the 16 methods with this reference (right). Precision, recall, and F1 score are reported at the cell level, with methods ordered by F1 score. Background shading indicates recovered ($F1 \geq 0.65$), partial ($0.50 \leq F1 < 0.65$), and not recovered ($F1 < 0.50$) outcomes.

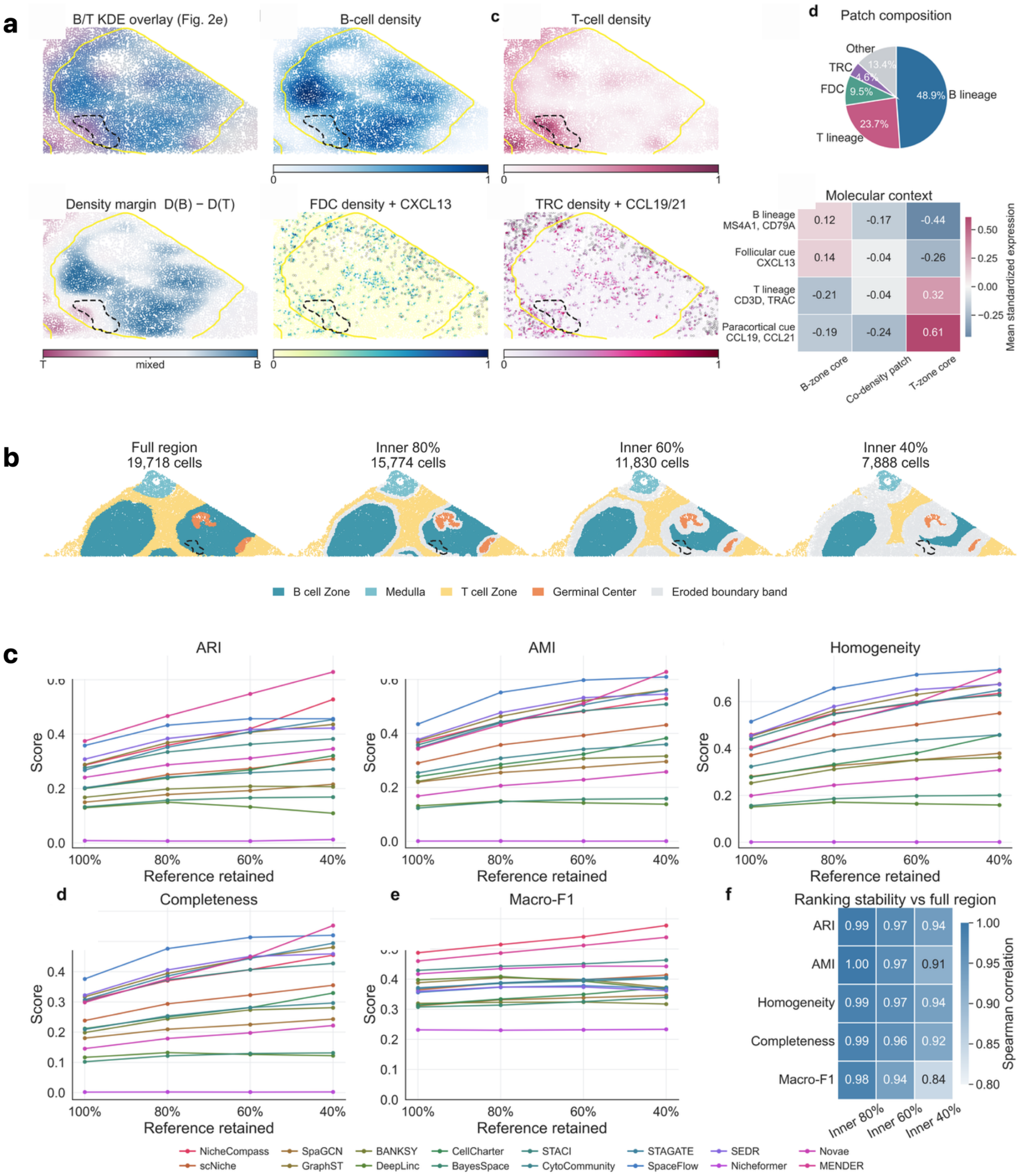


| | B-zone core | Co-density patch | T-zone core |
|---|---|---|---|
| B lineage MS4A1, CD79A | 0.12 | -0.17 | -0.44 |
| Follicular cue CXCL13 | 0.14 | -0.04 | -0.26 |
| T lineage CD3D, TRAC | -0.21 | -0.04 | 0.32 |
| Paracortical cue CCL19, CCL21 | -0.19 | -0.24 | 0.61 |

| | Inner 80% | Inner 60% | Inner 40% |
|---|---|---|---|
| ARI | 0.99 | 0.97 | 0.94 |
| AMI | 1.00 | 0.97 | 0.91 |
| Homogeneity | 0.99 | 0.97 | 0.94 |
| Completeness | 0.99 | 0.96 | 0.92 |
| Macro-F1 | 0.98 | 0.94 | 0.84 |

**Supplementary Fig. S20. Boundary uncertainty and robustness of the primary lymph-node benchmark.** (a) Characterization of the mixed B/T interface using lineage-density, stromal, and molecular context; dashed outlines indicate the examined mixed region and yellow contours denote the annotated B-cell-zone boundary. (b) Nested high-confidence niche cores obtained by progressively excluding cells near reference boundaries while preserving all four niche classes. (c) Benchmark performance on the full region and nested niche cores across ARI, AMI, homogeneity, completeness, and Macro-F1. The heatmap shows Spearman correlations between method rankings for each core set and the full-region ranking. Boundary exclusion increases absolute agreement while largely preserving relative method rankings.
